# Multi-Tissue Metabolomic Signatures of Five Longevity Interventions Converge on Ergothioneine and Lipid Remodeling in Male UM-HET3 Mice

**DOI:** 10.64898/2026.06.24.734388

**Authors:** Bretton Badenoch, Oliver Fiehn, Noa Rappaport, Sara Greenfield, Sriram Chandrasekaran, Richard A. Miller

**Affiliations:** Department of Molecular and Cellular Pathology, University of Michigan, Ann Arbor, MI, 48103, USA; UC Davis West Coast Metabolomics Center, University of California, Davis, CA, 95616, USA; Institute for Systems Biology, Seattle, WA, 98109, USA; Phenome Health, Seattle, WA, 98109, USA; Buck Institute for Research On Aging, Novato, CA, USA; Institute for Sustainability, Energy, and Environment at the University of Illinois Urbana-Champaign; Department of Biomedical Engineering, Center for Bioinformatics and Computational Medicine, Ann Arbor, MI, 48109, USA; Michigan Institute for Data and AI in Society, University of Michigan, Ann Arbor, MI, 48109, USA

## Abstract

The pace of aging can be delayed by mutations, dietary manipulations, and drugs, yet the metabolic mechanisms underlying longevity interventions remain poorly understood. Here we present a multi-tissue metabolomic analysis of male UM-HET3 mice treated from 4 to 12 months of age with five validated longevity interventions: rapamycin, acarbose, 17α-estradiol, canagliflozin, or caloric restriction. Using a feature-stabilized XGBoost pipeline applied to seven tissues, we show that metabolomic profiles can identify treated mice as likely recipients of a lifespan-extending intervention well before survival differences emerge. A leave-one-intervention-out procedure confirmed that models trained on any four interventions successfully classified mice from a fifth, unseen intervention, implying shared metabolic alterations across mechanistically distinct treatments. The most influential metabolites — defined as the minimum set explaining 50% of cumulative model gain — differed substantially across tissues. Only ergothioneine, a dietary antioxidant, ranked highly in more than two tissues: it was elevated by all five interventions in plasma and brain, and by four of five in muscle. Enrichment analyses further identified coordinated remodeling of lipid classes in plasma, perigonadal fat, and kidney. These findings reveal tissue-specific metabolic reprogramming shared across mechanistically distinct longevity interventions and, pending validation against interventions that do not extend lifespan, suggest a path toward metabolomic screening of candidate anti-aging drugs.

## Introduction

Aging is the main risk factor for most chronic diseases, including cardiovascular disease, neurodegeneration, metabolic dysfunction, and cancer.^1^ Interventions that slow the underlying biology of aging may therefore reduce the burden of most or all age-related conditions in parallel. The Interventions Testing Program (ITP), a multi-site initiative funded by the National Institute on Aging, has identified several compounds that extend lifespan in genetically heterogeneous UM-HET3 mice, including rapamycin, acarbose, 17α-estradiol, and canagliflozin.^2–5^ Caloric restriction (CR) also extends lifespan and healthspan in mice as well as in many other model organisms.^6^ The availability of multiple varieties of slow-aging mice permits a search for the metabolic mechanisms responsible for these benefits, and to determine whether molecular signatures are shared across different anti-aging interventions.

Metabolomics gives a sensitive readout of cellular and systemic physiology and captures the downstream result of genetic, transcriptomic, and proteomic changes. When used across multiple tissues, it can reveal both tissue-specific responses to longevity interventions and shared systemic signatures that may act as biomarkers of aging and mortality risk.^7^ Plasma metabolomics has been used to study aging and intervention responses.^7–10^ Our own work has shown that metabolite profiles from plasma can consistently discriminate control mice from those treated with any of four anti-aging drugs, or exposed to a CR diet.^8^ Analysis of the specific metabolites with highest impact on the calculated lifespan projection revealed that long-chain polyunsaturated triacylglycerols (TAG) are consistently elevated, and TAGs with short-chain fatty acids are diminished, in plasma of male UM-HET3 mice given anti-aging interventions.^7–9^

Machine learning methods, especially gradient-boosted trees such as XGBoost, are well suited to documenting shared features across diverse interventions, because they handle high-dimensional, correlated features and stay interpretable through feature importance.^11^ By training models to predict the extent of lifespan extension and then inspecting feature importance, we can find metabolites that are reliably changed by multiple, diverse longevity interventions. Because feature ranking can vary across runs that differ in random seed choice, particularly in high-dimensional settings,^12^ we have now developed a convergence-based method to find the minimum number of model iterations needed for stable feature rankings, thus greatly improving stability of feature ranking and reducing the effect of seed choice on biological interpretation.

Here we report a multi-tissue metabolomic analysis of genetically heterogeneous male UM-HET3 mice treated with rapamycin, acarbose, 17α-estradiol, canagliflozin, or caloric restriction from 4 to 12 months of age. Using a feature-stabilized XGBoost pipeline on metabolomic data from seven tissues, we identify metabolite signatures of longevity intervention shared by multiple interventions. Feature rankings defined several consistent tissue-specific alterations within classes of lipids. In addition, we noted consistent increases in plasma and brain levels of ergothioneine, a fungal metabolite that is absorbed from the gastrointestinal tract by the solute transport protein, SLC22A4/OCTN1.^13^ These results improve our understanding of shared metabolic changes underlying different longevity interventions and point to candidate mediators of their benefits. The models also have a practical benefit of being able to distinguish control and treated male mice after only 8 months of treatment, instead of performing a full lifespan assay.

## Results

### Estimation of lifespan increase from metabolite data from each of 7 tissues from 12-month-old male mice

We used an interpretable, tree-based machine learning framework (XGBoost) to estimate percentage lifespan increase in mice given anti-aging interventions or in untreated control populations, and to see which metabolites had the greatest influence on the predicted lifespan. Treated mice received one of five interventions from 4 months of age: rapamycin (Rapa, 14.7 ppm), acarbose (Aca, 1000 ppm), 17α-estradiol (17aE2, 14 ppm), canagliflozin (Cana, 180 ppm), or caloric restriction (CR) at 60% of the food intake of age-matched controls. At 12 months, we collected tissue from perigonadal fat, inguinal fat, brain, liver, muscle, and kidney, as well as plasma.

Treated male UM-HET3 mice could be distinguished from controls in all seven tissues (Panel A, Figures 1–4), using a procedure (the “novel intervention test” or NIT) that simulated evaluation of samples from a candidate drug whose effect on lifespan was not yet known. In this procedure, the model is constructed using data from untreated controls and from mice treated with four of the five treated groups, and then used to predict lifespan increase for each mouse in the fifth group, i.e. in the group not used for model training. The average percent increase in the estimated group is then compared to that in control mice using a t-test. The data for 17aE2 mice in Figure 1, for example, shows that the mean estimated lifespan in male 17aE2 mice is significantly higher than in control mice using a model trained on data from Cana, Aca, Rapa, and CR mice plus controls.

**Figure 1:**
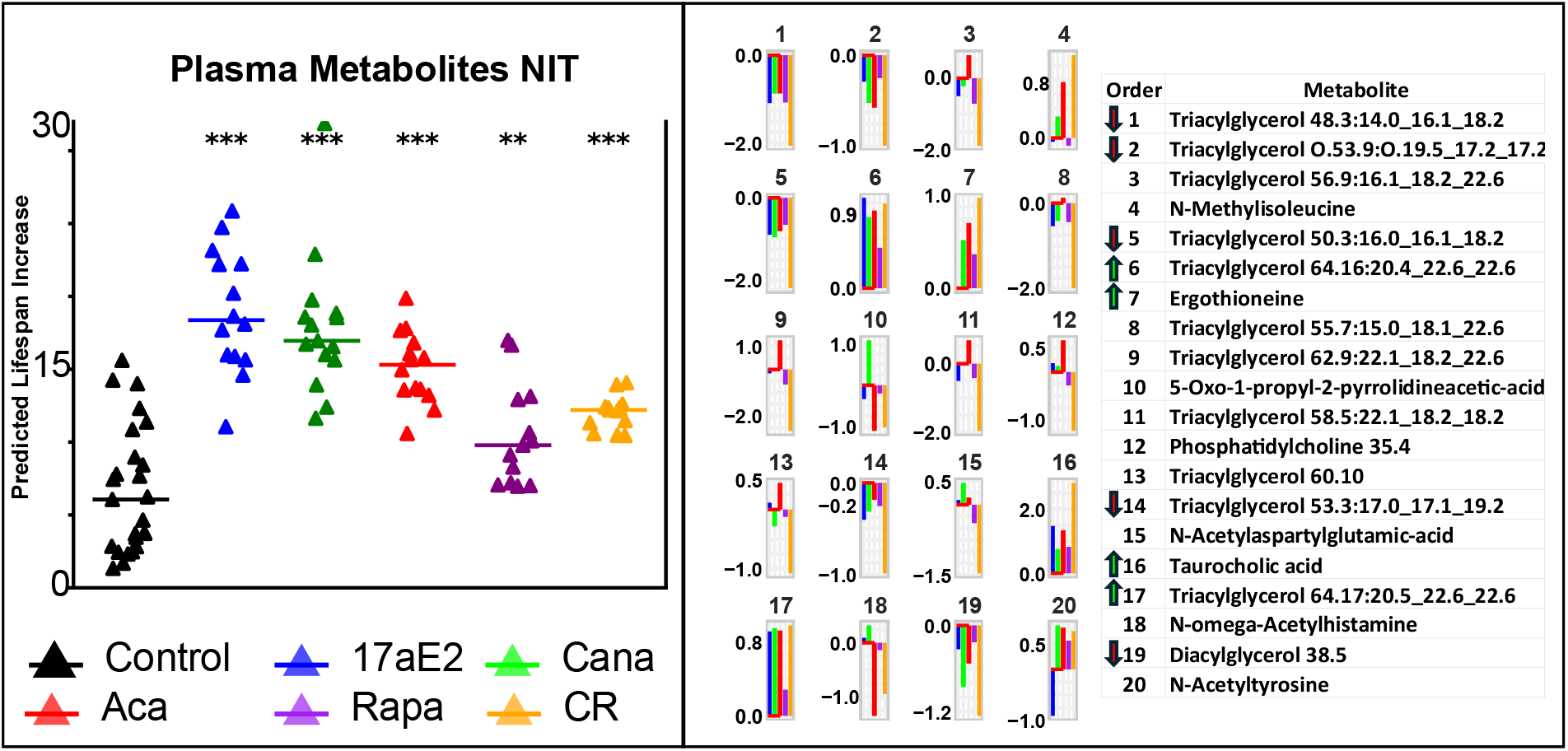
Novel Intervention Test and Feature Rank for Plasma. **(A)** Predicted percent lifespan increase for male UM-HET3 mice in each of five longevity intervention groups, estimated by XGBoost regression using a “novel intervention test” design. Each symbol represents one mouse. For each intervention group, a model was trained on control mice and the four remaining intervention groups, then applied to mice in the held-out group to generate predicted lifespan values; the 17α-estradiol group, for instance, was scored by a model trained on controls, acarbose, canagliflozin, caloric restriction, and rapamycin mice; within each held-out fold, the number of boosting rounds was selected by an inner tenfold cross-validation restricted to the training groups (leakage-free nested selection), and all other hyperparameters were fixed. Each held-out animal received a single predicted value from the trained model. Control predictions were generated by tenfold cross-validation across all mice (controls and treated combined), repeated over 20 iterations, with the per-animal median of the 20 predictions retained. Horizontal bars indicate group medians. Asterisks denote significance from Welch’s two-sample t-tests comparing each treated group to controls (*p < 0.05, **p < 0.01, ***p < 0.001). **(B)** Mean log_2_ fold changes for the 20 metabolites with highest XGBoost gain in plasma, shown as spark plots. Each bar represents the average log_2_ ratio of treated to control mice for the intervention indicated by color (see panel A legend). Feature ranks and identities correspond to the numbered key at right. Green up- or down-arrows denote metabolites that were, on average, consistently elevated or reduced across treated groups relative to controls.

Individual mice in the 17aE2 group have estimated lifespan increase values ranging from 12% to 26%, as shown in Figure 1. The results shown in Figure 1 differ from our previous report on plasma metabolites^8^ in that the current, improved procedure, shows values averaged over 1000 iterations each with its own random seed, a protocol that produces better stability than any one run of the model. Thus plasma metabolite patterns would have been able to predict extended lifespan for any of the five interventions shown in Figure 1, for male mice.

Figures 2, 3, and 4 (left panels) show that this procedure would, similarly, have been able to discriminate control mice from mice exposed to any of the five interventions if the models were trained on metabolite data from perigonadal or inguinal fat, brain, liver, muscle or kidney. The number of metabolites that were measured in each tissue and used for each tissue-specific ML model is listed in Supplemental Table 1.

**Table 1.**
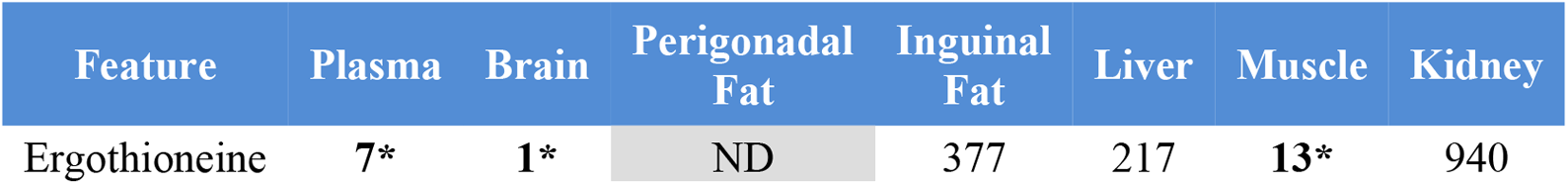
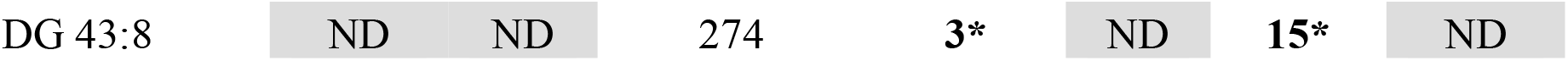
Metabolites shared as important features in two or more tissues at 50% feature explainability power. The bold numbers with the * represent the median XGBoost gain ranking of the metabolite after 1,000 iterations. “ND” means the metabolite was not detected by LC-MS or identified in post processing. These represent the median XGBoost gain ranking of the metabolite after 1,000 iterations. All bolded numbers were significant by a two-tailed t-test comparing pooled treated UM-HET3 males, and their controls. The number of metabolites it took for plasma, brain, PG fat, IG fat, liver, muscle, and kidney to explain 50% of the prediction are 10, 11, 5, 3, 2,15, and 4, respectively.

**Figure 2:**
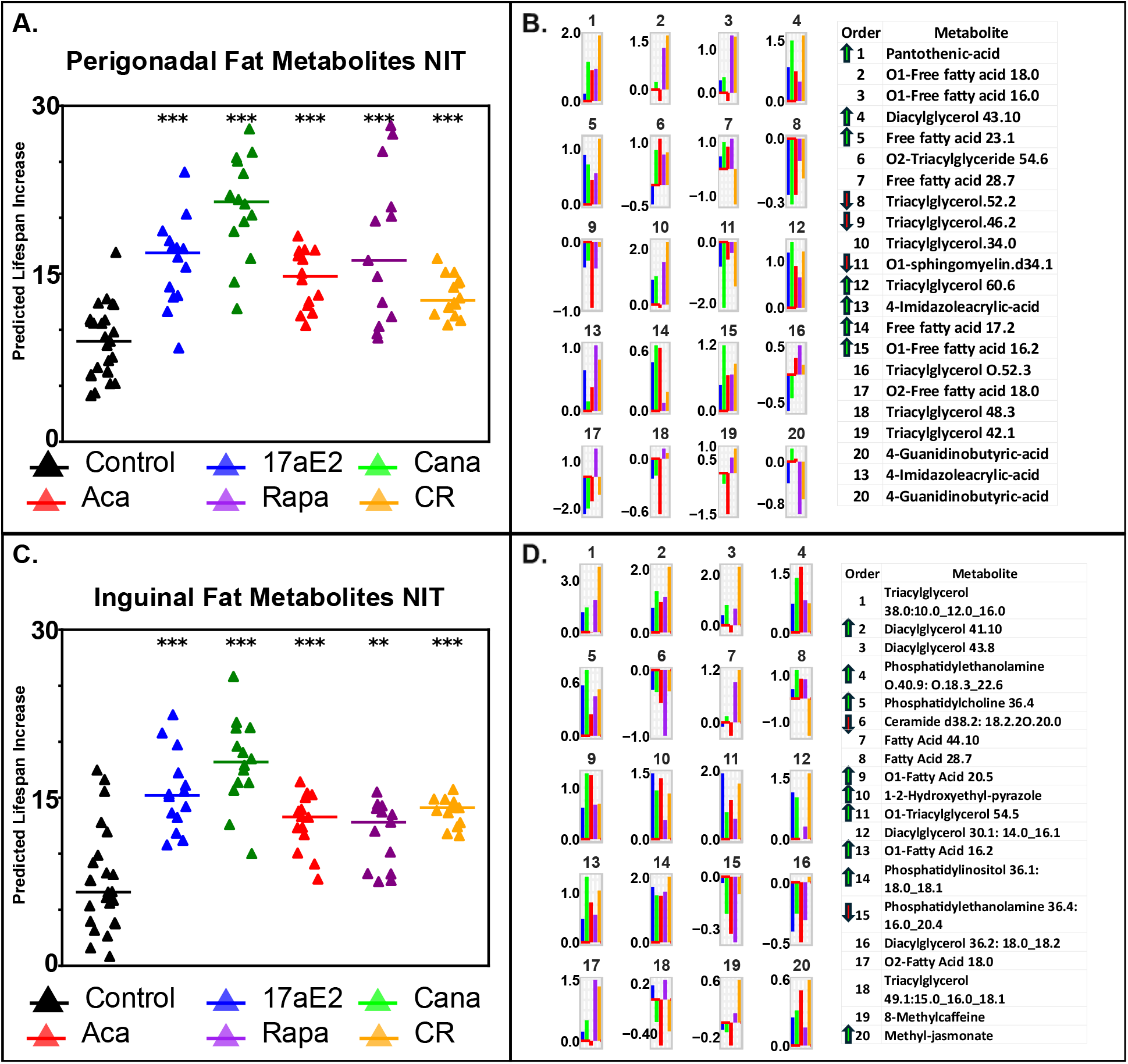
NIT and Feature Rank for Perigonadal and Inguinal Fat. See figure 1 legend.

**Figure 3:**
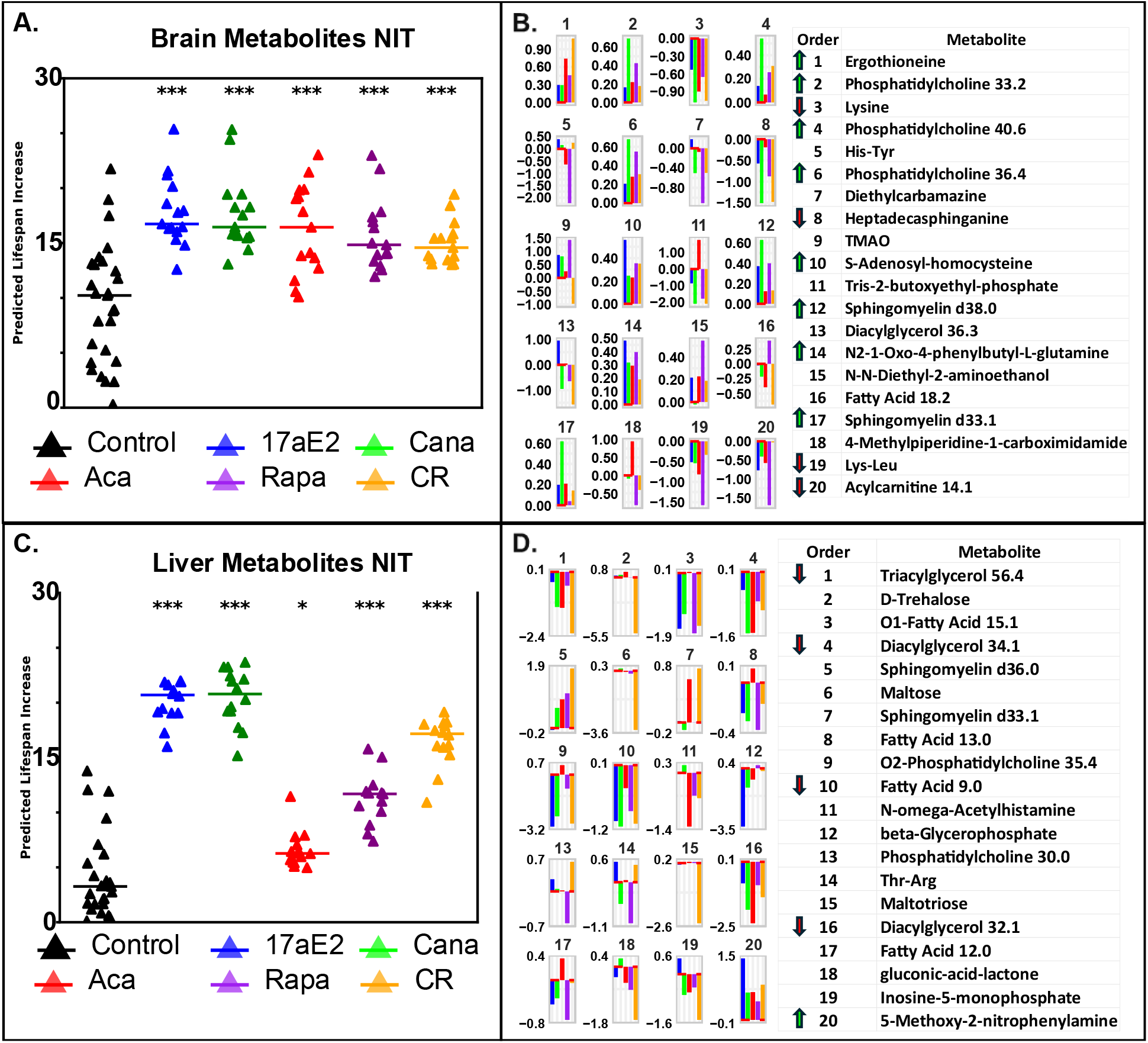
Novel Intervention Test and Feature Rank for Brain and Liver. See figure 1 legend.

**Figure 4:**
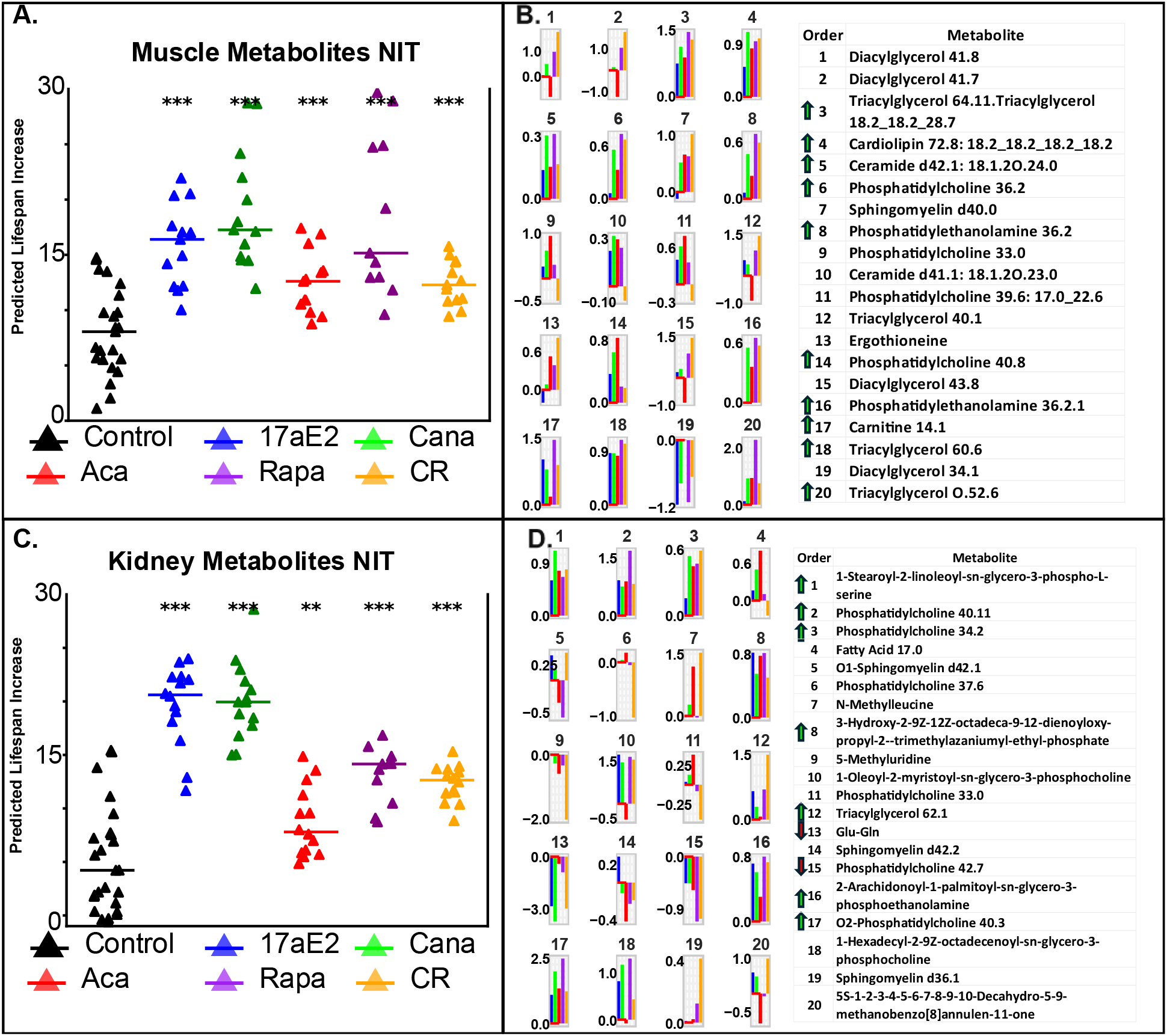
Novel Intervention Test and Feature Rank for Muscle and Kidney. See figure 1 legend.

### Ranking individual metabolites for effect on lifespan prediction

To see which metabolites had the largest influence on lifespan estimates in these models, we tabulated the top predictive metabolites as ranked using the XGBoost “Gain” function. Since each model requires initiation using a random seed, rank-order for individual metabolites varies across runs. We therefore conducted a series of experiments to determine how many model iterations were needed for the ranked listing of metabolite sets to converge. To quantify convergence, we computed two metrics at each iteration step (1, 2, 4, 10, 25, 50, 100, 200, 300, 400, 500, 600, 700, 800, 900, and 1,000 iterations): Spearman rank correlation (ρ_T_) between successive importance vectors, and total variation distance (D_T_), both evaluated on a fixed top-feature set defined at 1,000 iterations. We defined convergence as the first iteration at which ρ_T_>= 0.98 and D_T_ <= 0.02 were sustained for two successive steps. Under these criteria, brain and liver converged earliest at 300 iterations, plasma, gonadal fat, and inguinal fat converged at 400 iterations, and kidney and muscle required 500 iterations. By 1,000 iterations all tissues achieved ρ_T_>= 0.996 and D_T_ <= 0.008, indicating near-perfect rank stability. These results are summarized in Supplementary Table 6. We therefore concluded that calculating median values for Gain over a set of 1000 iterations would yield a stable rank ordered listing. Panel B of Figures 1 – 4 display the 20 most informative metabolites for each tissue in the form of sparkplots, i.e. log2-fold changes for each intervention for each of the 20 metabolites averaged over 1,000 iterations. These panels also include a table naming each of the 20 metabolites. Red arrows in the table indicate metabolites which are diminished in each of the five interventions, and green arrows indicate metabolites increased by all five interventions. Plasma had 9 such unidirectional features among the 20 top metabolites. The corresponding values in the other tissues were: PG fat 10, Ing fat 11, brain 12, liver 5, muscle 10, and kidney 9. Complete tables showing all rank-ordered metabolites are included in the Supplementary Table 5.

It clear from the tables of metabolites in Figure 1 – 4 that there is very little overlap across tissues; metabolites ranking in the top 20 by influence in any one tissue are seldom highly ranked in any other tissue, including plasma. To formalize this impression, we calculated, for each tissue, the number of metabolites needed to account for 50% of the model’s predictive power. For plasma, brain, PG fat, ING fat, liver, muscle, and kidney those numbers were respectively 10, 11, 5, 3, 2, 15, and 4. Only two metabolites were in the top features responsible for 50% of the model’s predictive power in two or more tissues: ergothioneine and DG 43:8 (Table 1). Ergothioneine was significantly increased when pooling male treated mice (Rapa, Aca, Cana, 17aE2, and CR vs UM-HET3 Control mice) in the brain, plasma, and muscle, separately (Table 1). DG 43:8 was significantly increased in ING fat and muscle (P < 0.05) (Table 1). Each of the five interventions led to an increase in ergothioneine in plasma and in brain, and each intervention except 17aE2 led to increased ergothioneine in muscle (Figure 6). DG 43:8 was decreased by Aca in ING fat and muscle.

We also tested the idea that specific classes of metabolites might be over-represented within the set with highest predictive power, by compiling lists of metabolites needed to achieve 90% of predictive power for each tissue (Supplemental Table 2) and then applying a binomial test for skew to see if specific classes were over-represented within the subset needed to achieve 90% predictive power in each tissue. We found DGs were overrepresented in the plasma, TAGs were overrepresented in plasma; and PC in the kidney (Supplemental Table 2). To see if there were consistent patterns in the direction of changes for these over-represented classes, we plotted each metabolite within these classes with respect to lipid length and saturation (Figure 5). The plasma result (Figure 5) confirmed our earlier report^8^ that anti-aging interventions increase levels of TAGs with long, polyunsaturated fatty acids and decrease levels of TAGs with short, saturated TAGs. Several of the tissue/class combinations showed a similar degree of consistency. DGs in plasma, for example, declined in treated mice. PCs in kidney generally increased, regardless of fatty acid chain length.

**Figure 5:**
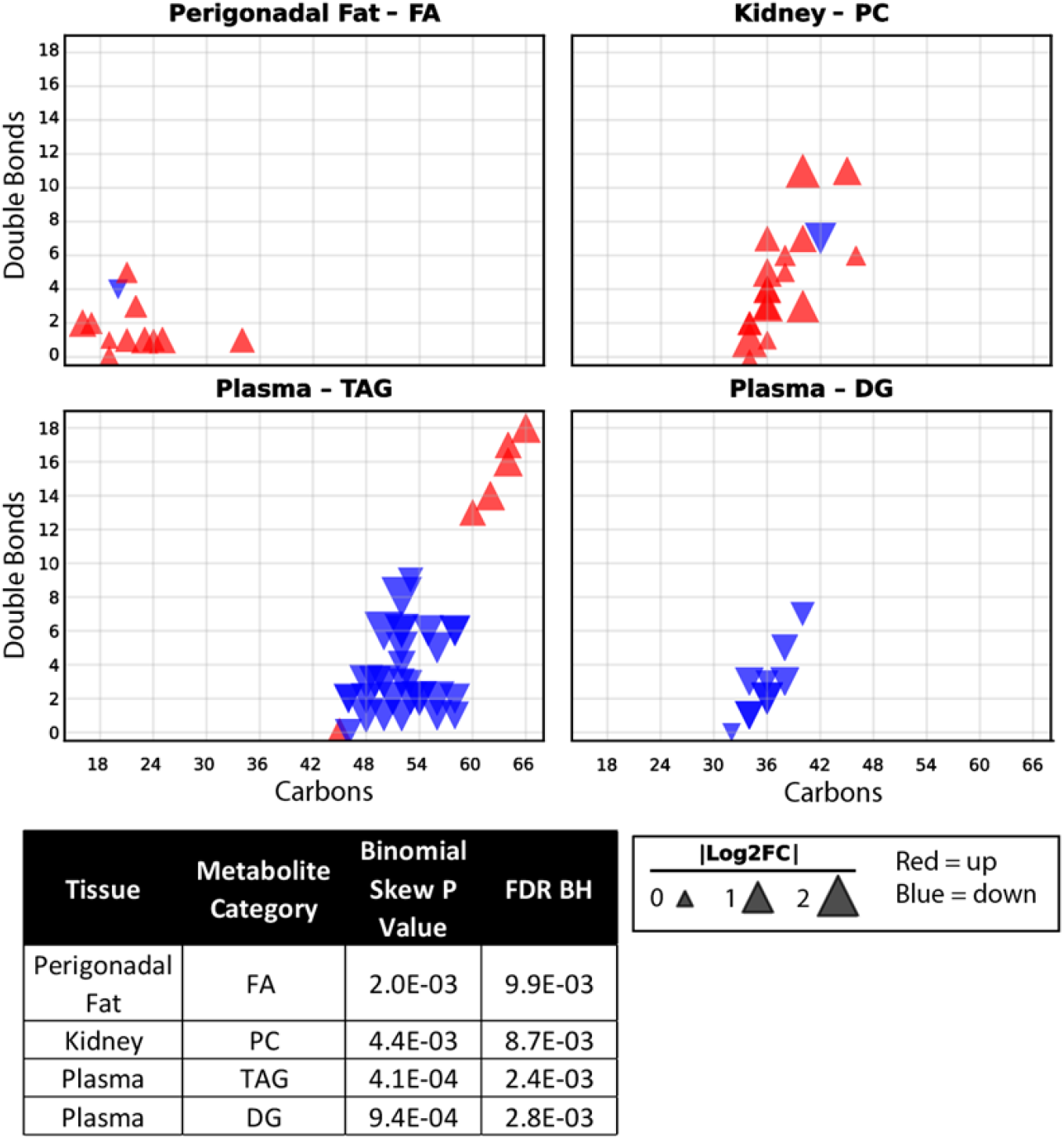
Patterns of changes in over-represented metabolite classes (Male mice) Overrepresented Lipids (males). This plot shows all lipid species that were changed in the same direction (blue = down, red = up) for all male UM-HET3 long-lived mice (Aca, Cana, CR, 17aE2, Rapa), with respect to UM-HET3 male controls. These lipid species were selected by a binomial test for skew, the p-value of these respective lipid classes are shown in the table above the legend for each tissue (the complete table can be viewed in Supplemental Table 2). The triangle size corresponds to the average |Log2FC| for all male long-lived mice, with respect to male UM-HET3 controls. Abbreviations: FA = fatty acid, DG = diacylglyceride, PC = phosphatidylcholine TAG = Triacylglyceride.

**Figure 6.**
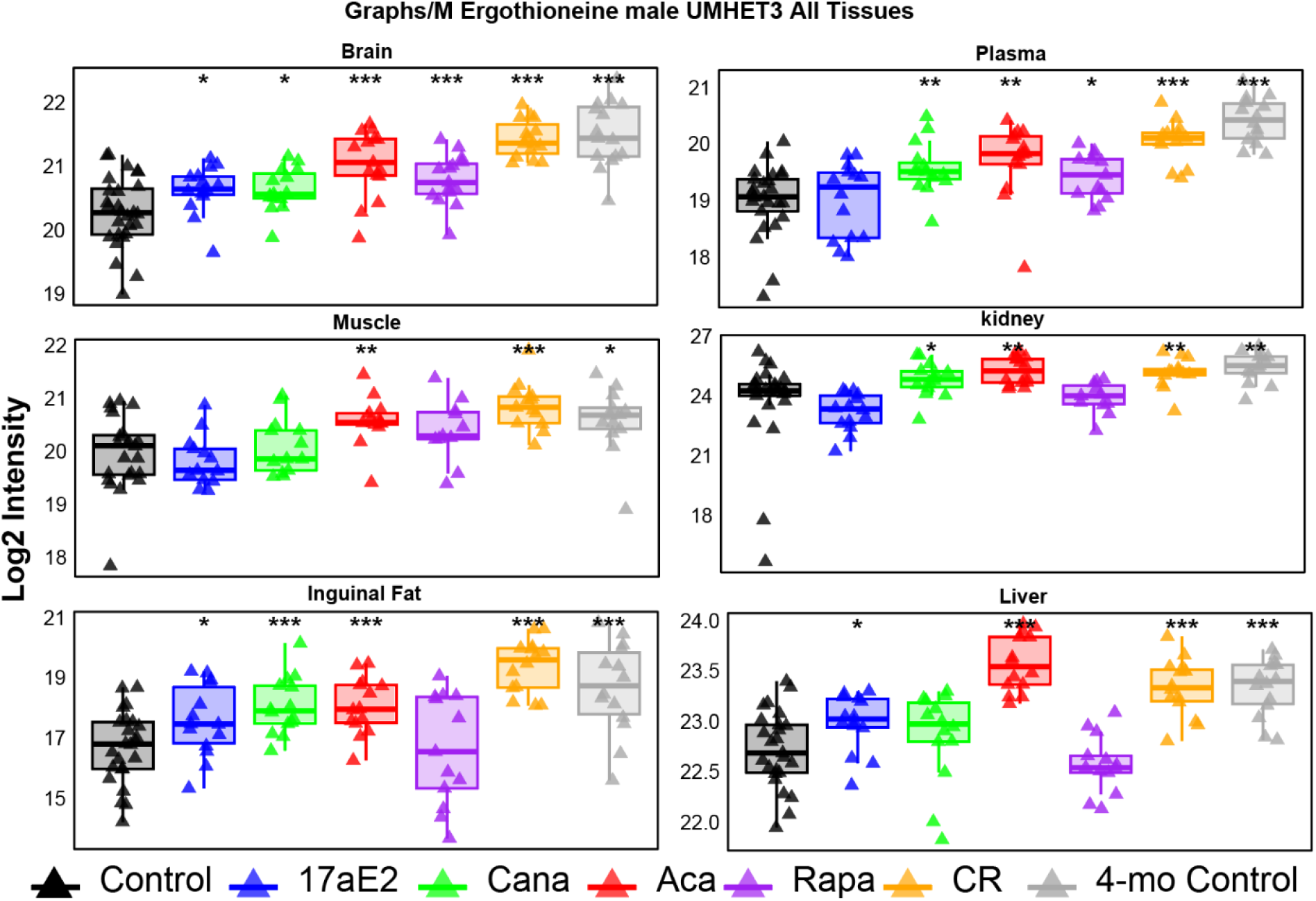
Ergothioneine levels across tissues in male UM-HET3 mice. Log2 intensity of ergothioneine in brain, plasma, muscle, kidney, inguinal fat, and liver. Ergothioneine was not detected in perigonadal fat and is therefore not shown. Male groups: 12-month control (black), caloric restriction (CR), rapamycin (Rapa), acarbose (Aca), canagliflozin (Cana), 17α-estradiol (17aE2), and 4-month control (grey). Boxes show quartiles with overlaid individual data (triangles). Symbols above each box indicate significance for that treatment group versus 12-month control (Welch’s two-tailed two-sample t-test): * = p ≤ 0.05, ** = p ≤ 0.01, *** = p ≤ 0.001.

## Discussion

We report a multi-tissue view of metabolomic effects in male UM-HET3 mice subjected to each of five longevity interventions. Using a feature-stabilized XGBoost pipeline, we show that treatment status, compared to untreated controls, is encoded in metabolite patterns in all seven tissues by 12 months of age in mice first exposed to interventions at 4 months. Most of the features that allow discrimination between treated and control mice are tissue-specific, with only one, ergothioneine, the top features explaining 50% of predictive power in more than two tissues. Several tissues, however, show systematic lipid remodeling after exposure to anti-aging interventions. Models trained on any set of four interventions can discriminate control mice from mice subjected to a fifth intervention not used in the training phase, suggesting that this approach may lead to a method for relatively rapid prioritization of previously untested candidate antiaging drugs, in mice and perhaps in humans. Although the power of the XGBoost approach reflects its ability to use information from metabolites that do not necessarily respond to all antiaging interventions to the same extent or even in the same direction, many of the most informative metabolites in each tested tissue do show directionally consistent changes in response to all five interventions tested. This consistency suggests the hypothesis that these might change in the same direction in response to a wider array of anti-aging interventions, and thus serve as “aging rate indicators”.^14^

XGBoost regression models separate groups of treated from control mice in all seven tissues with acceptably high power, producing significant differences from controls with group sizes of 10 – 15 mice. The data are consistent with other work^14^ showing that longevity interventions cause changes in fat, liver, muscle, brain, plasma, and macrophage subsets by mid-life, long before mortality risks begin to increase perceptibly.^3, 6, 15, 16^

Ergothioneine was elevated by all five interventions in brain and plasma, and by four interventions in muscle. With respect to its influence on predicted lifespan increase, ergothioneine was the most important feature in brain and ranked seventh in plasma and 13^th^ in muscle. Ergothioneine is a histidine-derived thiol antioxidant^17^, and can be synthesized by fungi and some bacteria but not by mammals.^18^ Mammals can obtain ergothioneine from the diet.^19^ Absorption from the intestine is thought to require the high-affinity transporter SLC22A4 (OCTN1).^13^ It is not yet clear if increased tissue levels of ergothioneine convey health benefits, or if elevated tissue ergothioneine is a surrogate marker for increased absorption of other materials that are important to health maintenance. Ergothioneine accumulates to millimolar levels in tissues that express SLC22A4^13^, and it may act as a neuroprotective antioxidant^20^. Low blood ergothioneine levels have been found to correlate with cognitive decline and mortality risks in humans.^21, 22^ Oral ergothioneine has been demonstrated to improve memory function in mice and human (healthy and mildly cognitively impaired)^23^, and prevent neuronal injury induced by amyloid B in mice.^24^ Further work is in progress to determine if slow-aging mice differ from controls in levels of SLC22A4 in gastrointestinal tissue, which might explain higher plasma levels of ergothioneine in these mice.

Cardiolipin 72:8 (constituent fatty acids 18:2_18:2_18:2_18:2), was increased in muscle by each of the five longevity interventions (Figure 4, Panel B), and was ranked #4 among muscle metabolites with respect to influence on lifespan prediction. Cardiolipin is a dimeric phospholipid of the inner mitochondrial membrane and is required for oxidative phosphorylation, cristae structure, and calcium handling in skeletal muscle.^25, 26^ Cardiolipin 72:8 is decreased by age in the hearts of rats^27^, mice^28^, and fish^29^. In human adult and pediatric patients with heart failure, Cardiolipin 72:8 is significantly lower in the left ventricle compared to those without heart failure.^30^ Lower levels of Cardiolipin 72:8 has been hypothesized to be one of the causes of mitochondrial dysfunction in aging, resulting in sarcopenia.^7, 26^ That a linoleate-rich cardiolipin species (72:8, four 18:2 chains) is consistently elevated in muscle of long-lived male mice across interventions suggests that preservation or enhancement of mitochondrial cardiolipin may be part of a shared metabolic phenotype of longevity in this tissue. Whether this reflects better mitochondrial maintenance, reduced oxidative damage to cardiolipin, or altered biosynthesis or remodeling will require further work.

Our earlier work on plasma metabolites^8^ had shown a consistent pattern: elevation of TAGs with long-chain fatty acids accompanied by decline in TAGs with short-chain fatty acid. This pattern is confirmed in our current study, using the same plasma dataset but using 1000 iterations to improve stability of rank-order among predictive metabolites. We have also extended the analysis to other tissues, and found patterns in multiple tissues: lower DGs and higher long-chain polyunsaturated TAGs in plasma, higher fatty acids in PG-fat, and higher PCs in kidney. The shift toward long-chain, polyunsaturated TAGs and DGs and away from short-chain, saturated species in plasma and PG fat may be due to greater flux through elongation and desaturation and may reflect better membrane fluidity and lipid signaling in long-lived mice. Similar polyunsaturation patterns have been seen in long-lived invertebrate models.^31, 32^ The increase of ceramides and glucosylceramides among top muscle features is notable, since ceramide species are tied to insulin resistance and to autophagy and metabolic disease.^33, 34^ Whether these lipid shifts are required for improved health, or are merely surrogate markers of protective states will need additional study.

Several of the top brain metabolites that track with predicted lifespan increase (Figure 3, Panel B) have been linked to brain health or disease in other studies. In addition to ergothioneine, phosphatidylcholines and sphingomyelins in plasma or CSF have been broadly positively associated with age-related cognitive impairment and dementia risk; here, several PC and SM species were among the top brain features that increased with treatment.^35–37^ Notably, PC 40:6, the 4^th^ most important feature identified in the brain and increased in all treated male mice, was found to be depleted in the plasma of individuals who developed dementia and more “AD-like” patterns of brain atrophy.^35, 37^ Higher concentrations of SM 38:0 in the plasma was found to be positively correlated with higher cognition, measured by the Montreal Cognitive Assessment, in a cross-sectional adult human study.^36^

DG 43:8, the other shared multi-tissue feature, was elevated in ING fat and muscle in all interventions except acarbose in both tissues. That exception is interesting: acarbose inhibits intestinal alpha-glucosidases and changes post-prandial glucose and lipid absorption in ways that may differ from the other interventions for some DG species. DG 43:8 is highly unsaturated (8 double bonds over 43 carbons), so it fits the broader pattern of more polyunsaturated lipids in long-lived mice.

Several limitations of the present study merit discussion, and can be addressed by future research. First, the XGBoost regression method successfully distinguished groups of treated from control mice only in males. We were not able to train a successful model using data from females only (not shown), perhaps because only rapamycin and CR lead to strong lifespan benefit in females. Identification of specific sets of metabolites that show sex-specificity in drug effect may help to shed light on the basis for these failures. Second, we did not have access to metabolite profiles from mice treated with compounds known not to extend lifespan. For this reason, it is possible that the model is discriminating drug-treated from untreated mice, regardless of the ability of the drugs used to extend lifespan. Experiments to see if the NIT discriminates interventions that increase lifespan from mice given drugs with no lifespan benefit are now in progress. Third, it will also be valuable to test whether the feature-stabilized XGBoost approach generalizes to metabolomic data from other laboratories and from other mouse stocks. Applying the framework to species, such as dogs, in which breed structure alters body size and expected longevity, will provide a further test of the method.

XGBoost models are interpretable, in the sense that relative feature importance includes a subset of metabolites that reliably distinguish treated from control animals. Unidirectional change suggests testable causal hypotheses for further mechanistic evaluation. Elevated ergothioneine, for example, might contribute to improved health and lifespan in these mice, or might instead reflect some other process, such as an increase in uptake of specific classes of nutrients, that themselves affect health. Disentangling cause from consequence could come from experiments in which ergothioneine was administered to mice to see if it conveys health benefits.

Lastly, our results suggest that combinations of the metabolic features identified here might provide meaningful readouts of anti-aging effects of short-term drug administration in human volunteers. The search for drugs that postpone or decelerate human aging might be facilitated by the use of plasma metabolic profiles to learn which drugs produce changes similar to those seen in slow-aging mice

## Methods

### Mouse Samples

UM-HET3 mice were weaned between 19–21 days of age and group-housed (3 males/cage or 4 females/cage) under conditions established for the Interventions Testing Program.^38^ Caloric restriction was initiated at 4 months of age, with CR animals receiving 80% of the chow intake measured in age- and sex-matched controls, further reduced to 60% at 5 months. CR animals received food once daily between 8–9 am, while all remaining groups were fed ad libitum. Drug-treated UM-HET3 mice began their respective interventions at 4 months of age. Animals were not fasted prior to tissue collection. Euthanasia was performed between 8–11 am at 11–12 months of age via CO2 asphyxiation; loss of consciousness occurred within approximately 10 seconds, after which closed-chest cardiac puncture was performed immediately upon cessation of breathing (typically within 30 seconds). Blood was drawn by closed-chest cardiac puncture and immediately transferred into K2EDTA-coated Vacutainer tubes, which were gently inverted to mix the blood with anticoagulant. Whole blood was then transferred into 1.5 mL microcentrifuge tubes for centrifugation. Plasma was isolated, aliquoted, and stored at −80°C until analysis.

All procedures were reviewed and approved by the University of Michigan Institutional Animal Care and Use Committee (IACUC) and conducted in accordance with institutional guidelines established by the University’s Unit for Laboratory Animal Medicine. All mice were bred and raised at the University of Michigan from founder stock obtained from the Jackson Laboratory.

### Metabolite Profiling of Tissues

Plasma metabolite measurements were obtained using a multi-platform mass spectrometry workflow comprising four liquid chromatography-accurate mass spectrometry (LC-MS) methods and one gas chromatography-mass spectrometry (GC-MS) assay, as described in detail elsewhere.^10^ In brief, 20 μl of plasma was extracted in 1 ml of methanol/water/MTBE, yielding distinct lipophilic and hydrophilic fractions. Both fractions were dried and processed separately for downstream LC-MS and GC-MS analysis.^39^ Polar metabolites were measured by hydrophilic interaction chromatography (HILIC) coupled to an Orbitrap QE HF instrument under both positive and negative electrospray ionization modes.^40^ Lipid profiling was conducted using C18 reversed-phase liquid chromatography (RPLC) on the same Orbitrap platform under positive and negative ionization modes.^41^ Volatile and semi-volatile metabolites were quantified by positive-mode electron ionization GC-MS on a Leco Pegasus IV GC-TOF instrument. LC-MS datasets were processed in MS-DIAL 4.90, with compound identification supported by the NIST20 and MassBank.us spectral libraries. GC-TOF MS data were processed in ChromaTOF 4.0 and annotated against the BinBase database.^42^ Batch normalization was performed using SERRF, a random forest-based machine learning approach calibrated to pooled quality control samples.^43^ Annotated metabolites (Metabolomics Standards Initiative levels 1–3) were retained at signal-to-noise ratios >3; unannotated features were subject to a more stringent threshold of s/n >10 relative to method blank controls for local noise estimation.^44^

### Novel Intervention Test and Model Training

We trained gradient-boosted regression models to predict a median lifespan-increase phenotype from tissue metabolomics and used them in a leave-one-intervention-out “novel intervention test” (NIT) and to estimate feature importance.

Model and target. For each tissue, an XGBoost regression model^11^ was trained to predict median lifespan increase from log_2_-transformed metabolite intensities. Training used male control and five male intervention groups (rapamycin, acarbose, canagliflozin, 17α-estradiol, calorie restriction).

Model hyperparameters were chosen separately per tissue by grid search. We used 10-fold crossvalidation (GridSearchCV; scikit-learn) and minimized negative mean squared error. The search grid was n_estimators [200, 500], max_depth [3, 5, 7], learning_rate [0.05, 0.1, 0.2], subsample [0.6, 0.8, 1.0], and colsample_bytree [0.6, 0.8, 1.0]. The best combination for each tissue was saved and reused for the repeated 10-fold cross-validation (10kCV). A 10-fold cross-validation with multiple iterations was run per tissue using the same training groups and the tissue-specific best hyperparameters. Predictions for control samples from this step were used to define the control reference (median predicted value) for the NIT. Hyperparameters for the Novel Intervention Test were matched using a nested cross-validation scheme to avoid information leakage from the held-out intervention group. For each leave-one-group-out fold, model selection was performed using only the remaining training groups: an inner 10-fold cross-validation (shuffled, fixed seed) was run over a grid of n_estimators values (100, 200, 300, 500, 750, 1000), and the value minimizing the mean inner-CV mean-squared error was selected. The remaining XGBoost hyperparameters were fixed at max_depth = 3, learning_rate = 0.2, subsample = 0.8, and colsample_bytree = 0.6. The model was then refit on all remaining training groups using the selected n_estimators and used to predict the held-out group. Because the inner cross-validation never sees the held-out group, this avoids the leakage that arises when hyperparameters are tuned on the full dataset (which includes the eventual test group), while leaving all other aspects of the NIT unchanged.

For each intervention group in turn, we held out that group and trained an XGBoost model on the remaining groups (including control). The fitted model was applied to the held-out group only. The NIT summary statistic for that intervention was the difference between the median prediction in the held-out group and the median control prediction from the 10kCV step (Tx_Control = median prediction in left-out group − median control prediction). This leave-one-intervention-out procedure was repeated for each intervention group per tissue.

Feature importance was computed from the same XGBoost models using the built-in gain metric (XGBoost’s default for feature_importances_). Gain is the total reduction in squared error (loss) achieved by all splits that use that feature across the ensemble of trees. For each tissue, we ran 1000 fits with fixed seeds(1-1000), recorded each feature’s gain and its normalized value (feature gain / sum of all feature gains) per run, and then median gain and normalized gain across runs. Features were ranked by median gain; we also report the number of features required to explain a a chosen fraction of the total normalized gain (50% for the shared-feature analysis in Table 1; 90% for the enrichment analysis in Supplemental Table 2) for downstream plots and tables.

Software. Analyses used Python 3 with pandas, NumPy, scikit-learn (model selection and metrics), XGBoost (xgboost), and SciPy; R (dplyr, ggplot2, openxlsx, tidyr) was used for figure generation and downstream tables. Random seeds were fixed (e.g. XGBoost random_state, NumPy seed) for reproducibility.

### Category enrichment test

For each tissue (brain, muscle, liver, kidney, GFat, IFat, and plasma), we identified the minimum number of top-ranked features required to explain 90% of the cumulative gain by sorting features by absolute mean gain values and computing cumulative proportions. Metabolites were classified into functional categories (e.g., phospholipids, amino acids, fatty acids) using pattern matching on metabolite names. To test for category enrichment in the top features, we performed one-sided binomial tests comparing the observed proportion of each category in the top N features to its expected proportion in the full dataset, using the full-dataset proportion as the null probability. P-values were calculated using R’s binom.test() function, and categories with p < 0.05 were considered significantly enriched. We also report the Benjamini-Hochberg corrected p-values.

Results were compiled into summary tables showing only significant enrichment and into tissuespecific reports containing all tested categories.

### Tissue phosphatidylcholine comparison

Tissue phosphatidylcholine (PC) levels were compared between male control and pooled male longevity interventions (rapamycin, acarbose, canagliflozin, 17α-estradiol, calorie restriction). For each tissue, metabolite columns classified as PC (phosphatidylcholine) were identified using the same naming rules as in metabolite classification; LysoPC and sphingomyelin were excluded. Log_2_-intensity values were converted to linear scale, and values corresponding to imputed missingness (zeros) were omitted. For each tissue, the sum of linear intensities over all PC species was computed separately for control and for pooled treatment samples. The ratio of total treatment PC to total control PC is reported, along with its log_2_ value, in a supplementary table by tissue.

### Ergothioneine comparison between tissues

Ergothioneine (log2 intensity) was compared across male UM-HET3 groups and tissues. Data were from cleaned log2-transformed metabolomics (zeros set to NA). For each tissue (brain, liver, kidney, perigonadal fat, inguinal fat, muscle, plasma), we restricted to male groups: 12-month control, caloric restriction, rapamycin, acarbose, canagliflozin, 17α-estradiol, and young. For each treatment group within each tissue, we performed a two-tailed two-sample t-test versus 12-month control. Boxplots show the distribution of log2 intensity with overlaid individual values (triangles); significance symbols are placed at the third quartile within each box.

### Feature convergence algorithm

For each run r ∈ {1,…,K}, raw feature importance values were normalized by summing all feature importance, and using that to divide the individual feature importance, as described in the equation below. For a fixed feature, across seeds there are k samples. For a given iteration budget, feature-wise robust summaries were computed; the median importance vector was renormalized across features. Top-n features were defined as the number of features that contain 95% of the feature importance from the run at the largest iteration number. Features were sorted in descending order. The minimum number of renormalized features whose cumulative mass ≥ 95% defines the set T. We required stable feature rankings and stable feature importance. We computed Spearman correlation of the rankings at consecutive budgets for the top T features and defined feature importance shift as the fraction of importance mass reassigned within T. The stopping rule for the number of iterations k (e.g., 10, 100, 1000) was defined as the smallest number of iterations such that stability holds consistently.

For each run r ∈ {1,…,K}, raw feature importance *s*_*f,r*_ were normalized to obtain:

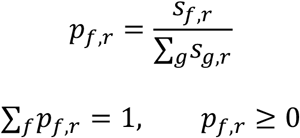

For a fixed feature *f*, across *k* seeds, there are samples: *p*_*f*,1_, *p*_*f*,2_, …, *p*_*f,k*_

For a given iteration budget *k*, feature-wise robust summaries were computed as:

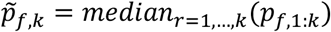

Because 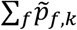 is not guaranteed to equal 1, the median importance vector was renormalized across features:

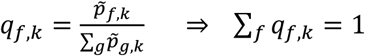

Define top-n features *T* as the number of features that have 95% of the mass from the run *r* at the largest iteration budget *K*.

Sort features by *q*_*f,K*_ descending. Let *π*(1), *π*(2), …be a permutation such that

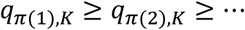

The minimum number of renormalized features *m* whose cumulative mass ≥ 95%:

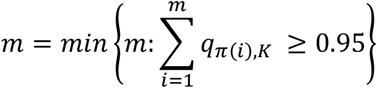

The set T is defined as the set of top features from 1 to *m*:

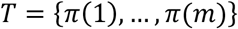

We want stable feature rankings, *π*_*k*_, and stable feature mass. so we compute Spearman correlation between the rankings at consecutive budgets for top T features:

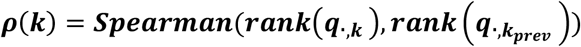

We define feature mass shift as the fraction of importance mass that got reassigned within T. The relative distribution on T is defined as:

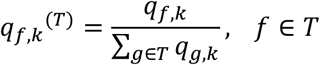

The fraction of importance mass *D*_*T*_ that got reassigned within T is defined as:

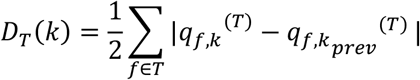

We define the stopping rule for budget k, where *k*_1_ < *k*_2_ < ⋯ < *k*_*M*_(e.g., 10, 100, 1000).

Define 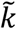 as the smallest budget such that stability holds persistently:

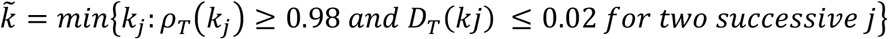

## Data Availability Statement

All data generated in this study are publicly accessible through the ELITE Portal (https://eliteportal.synapse.org), a resource developed to support open-science practices across NIA-funded Exceptional Longevity projects. The portal provides early-cycle access to data, analyses, and tools without publication embargoes on secondary use, facilitating translational research. Access and attribution requirements are outlined at https://eliteportal.synapse.org/DataAccess.

Code used in this manuscript is publicly available at: https://github.com/BrettonB/tissue-metabolomics-aging

## Funding

Research reported in this publication was supported by National Institute on Aging grants U19AG023122 and AG024824, the Career Training in the Biology of Aging T32 - NIH/NIA T32-AG000114, and R35 GM137795.

Research reported in this publication was supported by the National Institute on Aging of the National Institutes of Health under Award Number U19AG023122. The content is solely the responsibility of the authors and does not necessarily represent the official views of the National Institutes of Health.

## Supplemental Figures

**Supplementary table 1.**
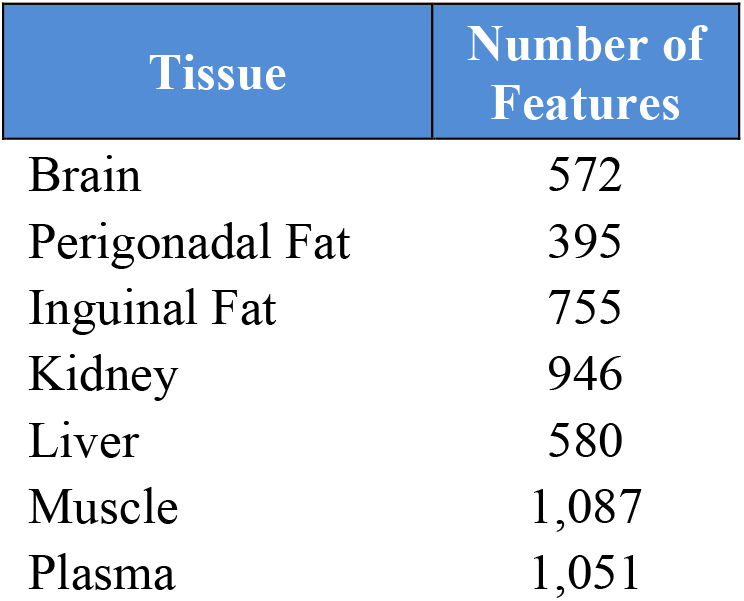
Number of metabolites (features) in each tissue dataset used in this study. Tissues: brain, perigonadal fat, inguinal fat, kidney, liver, muscle, plasma.

**Supplementary Table 2.**
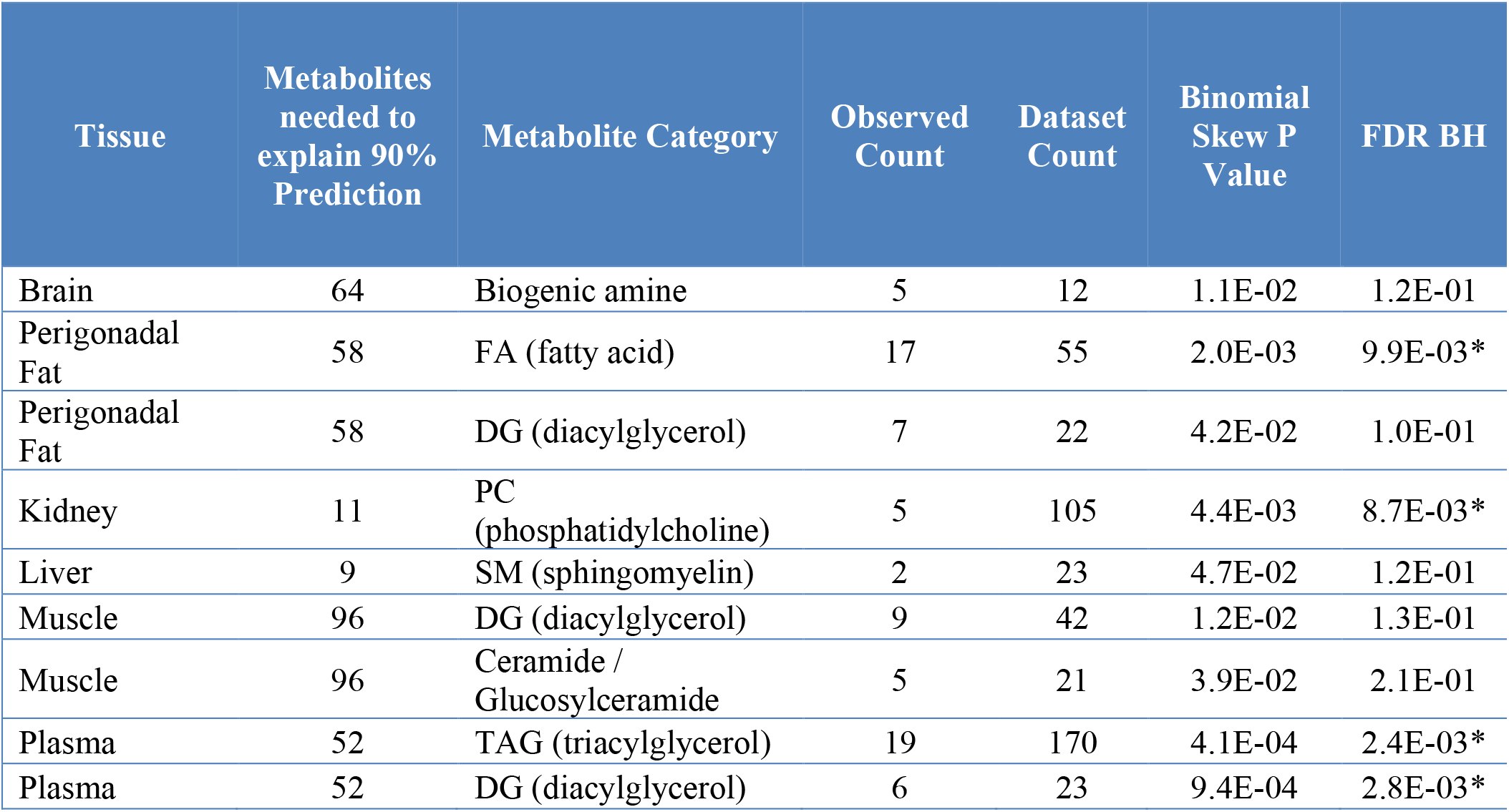
Category over-representation by tissue at 90% feature explainability power for each individual tissue. The “Dataset Count” is the number of times the metabolite category is found in the whole dataset for a particular tissue. The “Observed Count” is the number of times that the metabolite category is observed in the top 90% of predictive features, as given in the “Metabolites needed to explain 90% Prediction column”. The statistical test performed on the data was a binomial test for skew; raw and Benjamini-Hochberg adjusted (“FDR-BH”) p-values are reported.

**Supplementary table 3.**
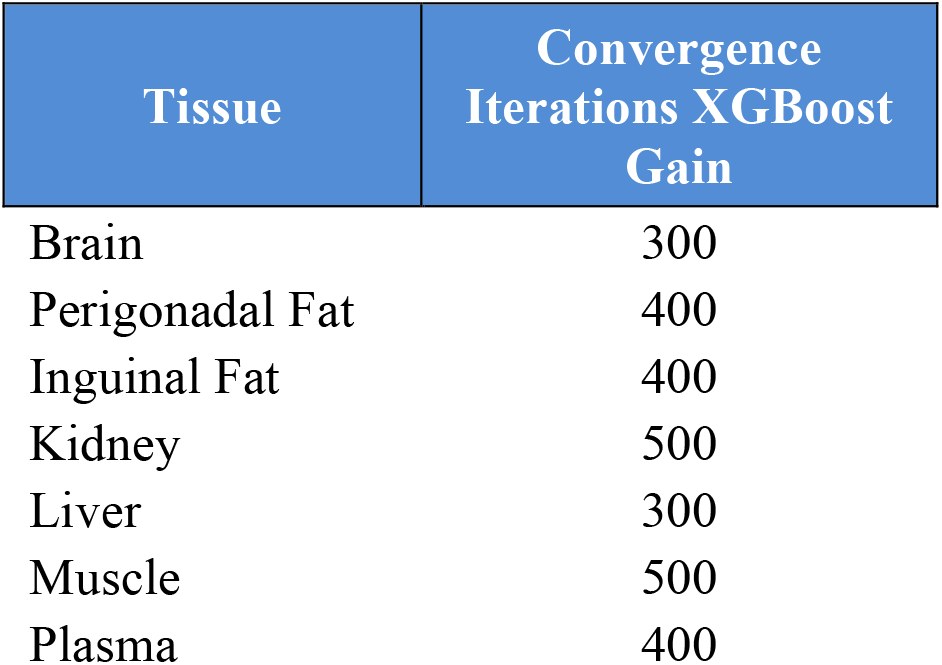
Number of iterations until convergence for the XGBoost algorithm applied to each tissue type. Tissues: brain, perigonadal fat, inguinal fat, kidney, liver, muscle, plasma.

**Supplementary table 4.**
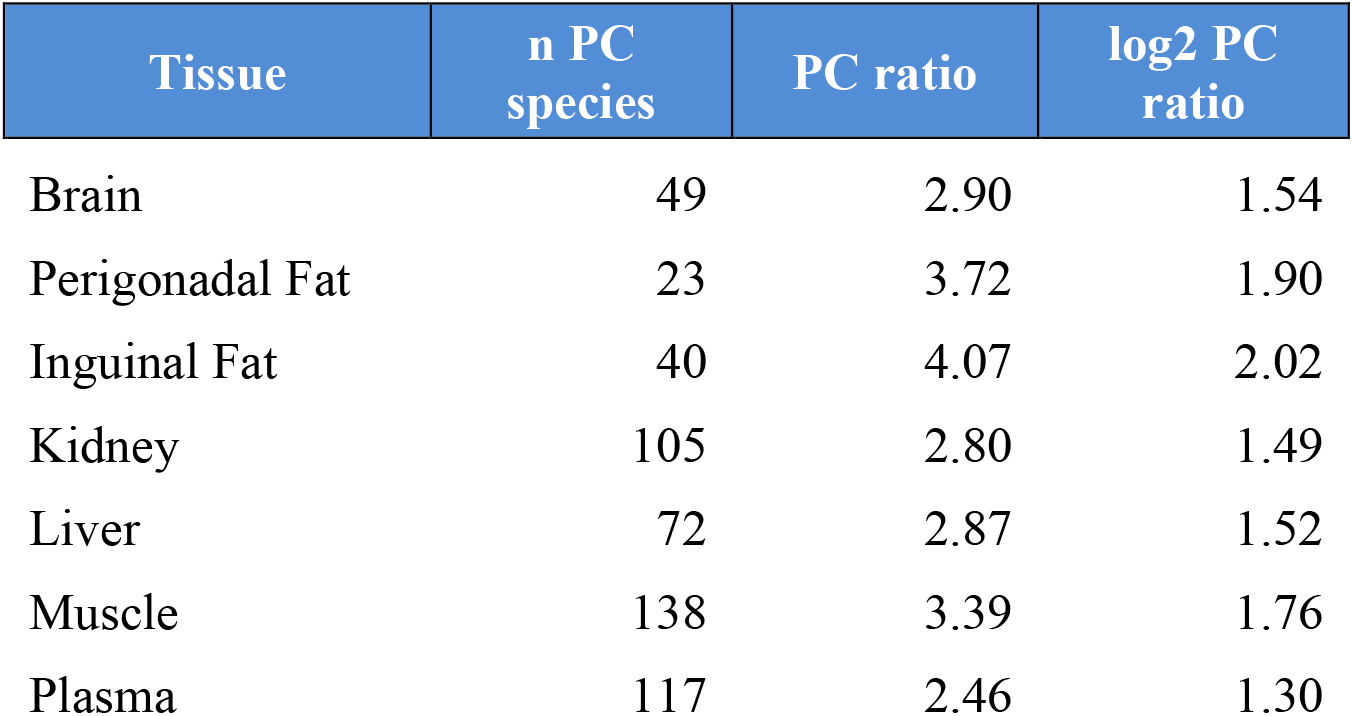
Ratio of total phosphatidylcholine (PC) in pooled male longevity interventions versus control, by tissue. For each tissue, metabolite columns classified as phosphatidylcholine (PC) were identified (LysoPC and sphingomyelin excluded). Linear-scale intensities (log2_intensity) were summed over all PC species and over samples in control or in pooled male treatment groups (rapamycin, acarbose, canagliflozin, 17α-estradiol, calorie restriction). The table reports the ratio of total treatment PC to total control PC and its log_2_ value (log2_PC_ratio). “n PC species” is the number of PC metabolites detected per tissue.

**Supplementary Table 5. Cross-tissue metabolite feature importance rankings from 1,000 XGBoost gain iterations.** Each row represents a unique metabolite detected across the seven tissue panels (brain, gonadal fat, inguinal fat, kidney, liver, muscle, and plasma). Columns display the final rank of each metabolite within each tissue, determined by median XGBoost gain importance averaged over 1,000 iterations with unique random seeds. Lower ranks indicate greater predictive importance for the lifespan extension classification task. “ND” denotes metabolites not detected in a given tissue’s feature matrix. A total of 3,324 unique metabolites are represented, with individual tissue feature counts ranging from 395 (gonadal fat) to 1,087 (muscle).(too big to fit in word document, it is uploaded as its own, separate file.

**Supplementary Table 6.**
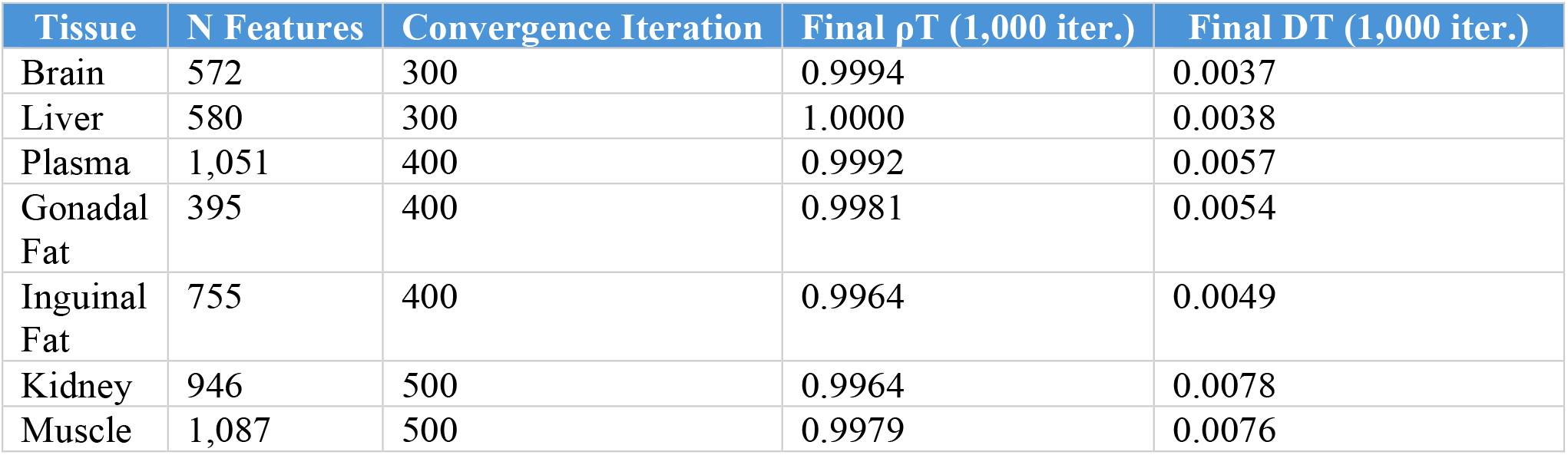
Convergence of XGBoost Gain feature importance rankings across tissues. Convergence iteration indicates the first step at which Spearman rank correlation (ρ_T_) >= 0.98 and total variation distance (D_T_) <= 0.02 were met for two successive iteration steps, evaluated on a fixed top-feature set defined at 1,000 iterations. All tissues achieved near-perfect rank stability (ρ_T_ > 0.99, D_T_ < 0.01) by 1,000 iterations.

## Notes

### Competing Interest Statement

The authors have declared no competing interest.

### Summary of Updates

We updated the formatting of references, added a GitHub link, and changed methods headers.

