## Supplemental Table 5 for "Multi-Tissue Metabolomic Signatures of Five Longevity Interventions Converge on Ergothioneine and Lipid Remodeling in Male UM-HET3 Mice"

| Feature | Plasma Rank | Brain Rank | GFat Rank | IFat Rank | Liver Rank | Muscle Rank | Kidney Rank |
|---|---|---|---|---|---|---|---|
| TG 48:3 \| TG 14:0_16:1_18:2 | 1 | ND | ND | 606 | ND | ND | ND |
| TG O-53:9 \| TG O-19:5_17:2_17:2 | 2 | ND | ND | ND | ND | ND | ND |
| TG 56:9 \| TG 16:1_18:2_22:6 | 3 | ND | ND | 648 | ND | ND | ND |
| N-Methylisoleucine | 4 | ND | ND | ND | ND | ND | ND |
| TG 50:3 \| TG 16:0_16:1_18:2 | 5 | ND | ND | ND | ND | ND | ND |
| TG 64:16 \| TG 20:4_22:6_22:6 | 6 | ND | ND | ND | ND | ND | ND |
| Ergothioneine | 7 | 1 | ND | 377 | 217 | 13 | 940 |
| TG 55:7 \| TG 15:0_18:1_22:6 | 8 | ND | ND | ND | ND | ND | ND |
| TG 62:9 \| TG 22:1_18:2_22:6 | 9 | ND | ND | ND | ND | ND | ND |
| 5-Oxo-1-propyl-2-pyrrolidineacetic acid | 10 | ND | ND | ND | ND | ND | ND |
| TG 58:5 \| TG 22:1_18:2_18:2 | 11 | ND | ND | 670 | ND | ND | ND |
| PC 35:4 | 12 | 318 | ND | ND | 528 | 24 | 552 |
| TG 60:10 b | 13 | ND | ND | ND | ND | ND | ND |
| TG 53:3 \| TG 17:0_17:1_19:2 | 14 | ND | ND | ND | ND | ND | ND |
| N-Acetylaspartylglutamic acid | 15 | 243 | ND | 362 | ND | ND | ND |
| Taurocholic acid | 16 | ND | ND | ND | 248 | 944 | 801 |
| TG 64:17 \| TG 20:5_22:6_22:6 | 17 | ND | ND | ND | ND | ND | ND |
| N-omega-Acetylhistamine | 18 | ND | ND | 345 | 11 | ND | 51 |
| DG 38:5 | 19 | 496 | 53 | ND | 34 | 794 | 337 |
| N-Acetyltyrosine | 20 | ND | ND | ND | ND | ND | ND |
| TG 66:18 \| TG 22:6_22:6_22:6 | 21 | ND | ND | ND | ND | ND | ND |
| PE 36:2 \| PE 18:0_18:2 | 22 | ND | ND | 465 | ND | ND | ND |
| SM 36:1;2O | 23 | ND | ND | ND | ND | ND | ND |
| 3-Hydroxybutyric acid | 24 | ND | ND | ND | ND | ND | ND |
| TG 60:13 \| TG 18:2_20:5_22:6 | 25 | ND | ND | ND | ND | ND | ND |
| TG 62:14 \| TG 18:2_22:6_22:6 | 26 | ND | ND | 698 | ND | 896 | 235 |
| DG 36:2 \| DG 18:1_18:1 | 27 | ND | ND | 37 | ND | ND | ND |
| TG 57:9 \| TG 17:1_18:2_22:6 | 28 | ND | ND | 661 | ND | ND | ND |
| TG 64:14 \| TG 18:2_22:6_24:6 | 29 | ND | ND | ND | ND | ND | ND |
| TG 51:2 \| TG 16:0_17:1_18:1 | 30 | ND | ND | 273 | ND | ND | ND |
| PC O-40:8 | 31 | ND | ND | ND | ND | 552 | ND |
| 2[4-(Trifluoromethoxy)phenyl]cyclopropanecarboxylic acid | 32 | ND | ND | ND | ND | ND | ND |
| Sulfamethoxazole | 33 | ND | ND | ND | ND | ND | ND |
| 3-Hydroxyvaleric acid | 34 | ND | ND | ND | ND | ND | ND |
| PI 36:1 \| PI 18:0_18:1_a | 35 | ND | ND | ND | ND | ND | ND |
| TG 58:8 \| TG 18:1_18:2_22:5 | 36 | ND | ND | ND | ND | ND | ND |
| DG 38:7 \| DG 18:2_20:5 | 37 | ND | ND | ND | ND | 795 | ND |
| DG 40:8 \| DG 18:2_22:6 | 38 | ND | ND | 434 | ND | 801 | ND |
| 3-Pyridinemthanol | 39 | ND | ND | ND | ND | ND | ND |
| DG 36:2 | 40 | 365 | 40 | ND | 541 | 784 | 667 |
| PC 42:10 \| PC 21:5_21:5 | 41 | ND | ND | ND | ND | ND | ND |
| N-Methyllysine | 42 | 32 | ND | ND | ND | ND | 208 |

| | | | | | | | |
|---|---|---|---|---|---|---|---|
| 3-(3-Hydroxyphenyl)propionic acid | 43 | ND | ND | ND | ND | ND | ND |
| DG 40:8 | 44 | ND | ND | ND | ND | ND | 289 |
| 1,5-Anhydro-D-sorbitol | 45 | ND | ND | ND | ND | ND | ND |
| 1-(Piperidin-4-yl)ethan-1-ol | 46 | ND | ND | ND | ND | ND | 32 |
| PC 42:7 | 47 | ND | ND | ND | ND | 36 | 15 |
| CAR 18:0 | 48 | ND | ND | ND | ND | 614 | ND |
| PE P-40:6 \| PE P-18:0_22:6 | 49 | ND | ND | ND | ND | 643 | ND |
| TG 61:13 \| TG 18:2_21:5_22:6 | 50 | ND | ND | ND | ND | ND | ND |
| SM 42:1;2O | 51 | ND | ND | ND | ND | ND | ND |
| LPC 16:1/0:0 | 52 | ND | ND | ND | ND | ND | ND |
| Cer 40:1;2O \| Cer 18:1;2O/22:0 | 53 | ND | ND | ND | ND | ND | ND |
| SE 29:1/18:2 | 54 | ND | ND | ND | ND | ND | ND |
| 5-alpha-Androstan-3-beta-ol-17-one sulfate | 55 | ND | ND | ND | ND | ND | ND |
| O2_PE 36:1 | 56 | ND | ND | ND | ND | ND | ND |
| N-alpha-Acetyl-arginine | 57 | 307 | ND | ND | ND | ND | ND |
| 10.87_603.53 | 58 | ND | ND | ND | ND | ND | ND |
| Deoxycholic acid | 59 | ND | ND | ND | ND | ND | ND |
| TG 60:11 \| TG 18:2_20:3_22:6 | 60 | ND | ND | ND | ND | ND | ND |
| 4-Imidazoleacetic acid | 61 | ND | 233 | 510 | ND | ND | ND |
| PC 40:5 | 62 | 477 | ND | 439 | 517 | 508 | 554 |
| PC 37:5 \| PC 17:0_20:5 | 63 | ND | ND | ND | ND | ND | ND |
| 1-Pentadecanoyl-sn-glycero-3-phosphocholine | 64 | ND | ND | ND | ND | ND | ND |
| gamma-Glutamyltyrosine | 65 | ND | ND | ND | ND | ND | ND |
| PC 36:4 \| PC 16:0_20:4 | 66 | ND | ND | ND | ND | ND | ND |
| N-acetylornithine | 67 | ND | ND | ND | ND | 1078 | ND |
| 1-Heptadecanoyl-sn-glycero-3-phosphocholine | 68 | ND | ND | ND | ND | 92 | 24 |
| LPC 22:1/0:0 | 69 | ND | ND | ND | ND | ND | ND |
| Naringenin-4'-O-.beta.-D-glucuronide | 70 | ND | ND | ND | ND | ND | ND |
| TG O-58:6 \| TG O-20:0_16:0_22:6 | 71 | ND | ND | ND | ND | ND | ND |
| TG 60:8 \| TG 20:0_18:2_22:6 | 72 | ND | ND | ND | ND | ND | ND |
| DG 38:7 | 73 | ND | ND | ND | ND | ND | ND |
| PC 38:7 | 74 | 146 | ND | ND | 513 | 498 | 561 |
| Glutamine | 75 | 457 | 382 | 317 | 461 | ND | 311 |
| (3-Oxo-2,3-dihydro-4H-1,4-benzoxazin-4-yl)acetic acid | 76 | ND | ND | ND | ND | ND | ND |
| (N-(-2-Acetamido))-2-aminoethanesulfonic acid | 77 | ND | ND | ND | ND | ND | ND |
| SM 41:1;2O | 78 | ND | ND | ND | ND | ND | ND |
| (3E)-4-(1-Hydroxy-2,2,6-trimethyl-4-oxocyclohexyl)but-3-en-2-yl hexopyranoside | 79 | ND | ND | ND | ND | ND | ND |
| TG 64:13 \| TG 18:1_22:6_24:6 | 80 | ND | ND | ND | ND | ND | ND |
| 3-Methylglutarylcarnitine | 81 | ND | ND | ND | ND | ND | 224 |
| Lactitol | 82 | 486 | ND | ND | ND | ND | ND |
| Homocitrulline | 83 | ND | ND | ND | ND | ND | ND |
| PI 36:2 | 84 | ND | ND | ND | ND | 668 | ND |
| Nicotinamide | 85 | 560 | 267 | 356 | ND | 420 | ND |

| | | | | | | | |
|---|---|---|---|---|---|---|---|
| TG 60:10 \| TG 18:0_20:4_22:6 | 86 | ND | ND | ND | ND | ND | ND |
| 12R-Hydroxy-5Z,8Z,10E,14Z-eicosatetraenoic acid | 87 | ND | ND | ND | ND | ND | ND |
| 2-Hydroxypalmitic acid | 88 | ND | ND | ND | ND | ND | ND |
| 2,6-Dihydroxybenzoic acid | 89 | ND | ND | ND | ND | ND | ND |
| PI 36:4 \| PI 16:0_20:4_a | 90 | ND | ND | ND | ND | ND | ND |
| FA 24:5 | 91 | ND | ND | 354 | ND | 57 | ND |
| PC O-32:0 | 92 | ND | ND | ND | ND | 525 | ND |
| Prolylalanine | 93 | ND | ND | ND | ND | ND | ND |
| gamma-Glutamylleucine | 94 | ND | ND | ND | ND | ND | 876 |
| Choline | 95 | 376 | 354 | 707 | 309 | 766 | ND |
| SM 36:1;2O \| SM 18:1;2O/18:0 | 96 | ND | ND | ND | ND | ND | ND |
| Ala-Thr | 97 | ND | ND | ND | ND | ND | 310 |
| TG 48:0 \| TG 16:0_16:0_16:0 | 98 | ND | ND | ND | ND | ND | ND |
| Gly-Pro | 99 | ND | 378 | ND | ND | ND | 327 |
| Taurodeoxycholic acid | 100 | ND | ND | ND | ND | ND | ND |
| 1,4-Cyclohexanedicarboxylic acid | 101 | ND | ND | ND | ND | ND | ND |
| SM 36:2;2O | 102 | ND | ND | ND | ND | ND | ND |
| PC 37:6 \| PC 15:0_22:6 | 103 | ND | ND | ND | ND | 274 | ND |
| 3-Hydroxy-3-methylglutaric acid_3-Hydroxy-3-methylglutaric acid | 104 | ND | ND | ND | ND | ND | ND |
| Suberic acid | 105 | ND | ND | ND | ND | ND | ND |
| 1-O-Hexadecyl-2-O-acetyl-sn-glyceryl-3-phosphorylcholine | 106 | ND | ND | ND | ND | ND | ND |
| 2-Hydroxy-4-methylpentanoic acid | 107 | ND | ND | ND | ND | ND | ND |
| gamma-Glutamylglutamine | 108 | ND | ND | ND | ND | ND | ND |
| PE P-42:6 \| PE P-20:0_22:6 | 109 | ND | ND | ND | ND | 382 | ND |
| (2R)-7-Methoxy-3-oxo-3,4-dihydro-2H-1,4-benzoxazin-2-yl .beta.-D-glucopyranoside | 110 | ND | ND | ND | ND | ND | ND |
| 2-Aminonicotinic acid | 111 | ND | 222 | ND | ND | ND | ND |
| N-Acetylglycine | 112 | ND | ND | ND | ND | ND | ND |
| SM 33:1;2O \| SM 17:1;2O/16:0 | 113 | ND | ND | ND | ND | ND | ND |
| 2-Thiobarbituric acid | 114 | ND | ND | ND | ND | ND | ND |
| (4-Methylphenyl)oxidanesulfonic acid | 115 | ND | ND | ND | ND | ND | ND |
| (3-Carboxypropyl)trimethylammonium | 116 | ND | 37 | 35 | ND | ND | ND |
| Methacholine | 117 | 114 | 329 | 88 | ND | 1063 | ND |
| FA 19:0 | 118 | ND | ND | 396 | ND | ND | ND |
| Asn-Val | 119 | ND | ND | ND | ND | ND | ND |
| 1-Stearoyl-2-hydroxy-sn-glycero-3-phosphoethanolamine | 120 | ND | ND | ND | ND | ND | ND |
| 3-Cyclopentene-1-octanoic acid, 2-(3-hydroxy-1-penten-1-yl)-5-oxo- | 121 | ND | ND | ND | ND | ND | ND |
| 2-Dec-9-enylpentanedioic acid | 122 | ND | ND | ND | ND | ND | ND |
| 1-Myristoyl-sn-glycero-3-phosphocholine | 123 | ND | ND | ND | ND | ND | ND |
| 1-Arachidoyl-2-hydroxy-sn-glycero-3-phosphocholine | 124 | ND | ND | ND | ND | ND | ND |
| 1-Methylpiperidine-3-carboxylic acid | 125 | ND | ND | ND | ND | 87 | ND |
| DG 36:4 \| DG 18:2_18:2 | 126 | ND | ND | 161 | ND | ND | ND |
| O2_FA 22:1; (erucic acid) [M-H]- | 127 | ND | ND | ND | ND | ND | ND |
| PC 38:3 \| PC 18:0_20:3 | 128 | ND | ND | ND | ND | ND | ND |

| | | | | | | | |
|---|---|---|---|---|---|---|---|
| (2E)-2-(Propan-2-yl)but-2-enedioic acid | 129 | ND | ND | ND | ND | ND | ND |
| TG 60:14\|TG 18:3_20:5_22:6 | 130 | ND | ND | ND | ND | ND | ND |
| Galactonic acid | 131 | ND | ND | ND | ND | ND | ND |
| (9E,11Z)-8-Hydroxyoctadeca-9,11-dienoic acid | 132 | ND | ND | ND | ND | ND | ND |
| TG O-60:6\|TG O-20:0_18:0_22:6 | 133 | ND | ND | ND | ND | ND | ND |
| Actrarit | 134 | ND | ND | ND | ND | ND | ND |
| Ser-His | 135 | 543 | ND | ND | 64 | ND | 686 |
| FA 22:6;O | 136 | ND | ND | ND | ND | ND | ND |
| Phosphoric acid | 137 | ND | ND | ND | ND | ND | ND |
| LPC 19:0/0:0 | 138 | ND | ND | ND | ND | ND | ND |
| (3-Oxo-2-piperazinyl)acetic acid | 139 | ND | ND | ND | ND | ND | ND |
| beta-Hydroxyisovaleric acid | 140 | ND | ND | ND | ND | ND | ND |
| HexCer 42:2;2O\|HexCer 18:1;2O/24:1 | 141 | ND | ND | 294 | ND | 971 | ND |
| (R)-Butyrylcarnitine | 142 | ND | 162 | ND | 36 | ND | 35 |
| 2-((2R)-2-Hydroxycyclohexyl)acetic acid | 143 | ND | ND | ND | ND | ND | ND |
| 3-Oxo-1,8-octanedicarboxylic acid | 144 | ND | ND | ND | ND | ND | ND |
| Acetyl-threonine | 145 | ND | ND | ND | ND | ND | ND |
| 3-Hydroxybutyrylcarnitine | 146 | 250 | 188 | ND | ND | 186 | 219 |
| 4,6-Dihydroxypyrimidine | 147 | ND | ND | ND | ND | ND | ND |
| Ala-Gln | 148 | ND | ND | ND | 130 | 208 | 849 |
| 1-Methyladenosine | 149 | 176 | ND | ND | 50 | 110 | 53 |
| SM 32:1;2O\|SM 16:1;2O/16:0 | 150 | ND | ND | ND | ND | ND | ND |
| 1-Hexadecyl-sn-glycero-3-phosphocholine | 151 | ND | ND | ND | ND | ND | 57 |
| 1-Oleoyl-sn-glycero-3-phosphocholine | 152 | ND | ND | ND | ND | ND | ND |
| Methyl-beta-galactopyranoside | 153 | ND | ND | ND | ND | ND | ND |
| 1-Hexadecanoyl-2-octadecadienoyl-sn-glycero-3-phosphocholine | 154 | ND | ND | ND | ND | ND | ND |
| (2R)-3-Hydroxyisovaleroylcarnitine | 155 | ND | 129 | ND | 41 | ND | 27 |
| 5-Aminosalicylic acid | 156 | ND | ND | ND | ND | ND | ND |
| (1,5-Dimethyl-1H-pyrazol-3-yl)methylamine | 157 | ND | 82 | 23 | ND | ND | ND |
| Cer 42:1;2O\|Cer 18:1;2O/24:0 | 158 | ND | ND | ND | ND | ND | 281 |
| (2-oxo-2,3-dihydro-1H-indol-3-yl)acetic acid | 159 | ND | ND | ND | ND | ND | ND |
| 4-Hydroxybenzaldehyde | 160 | ND | ND | ND | ND | ND | ND |
| 2-Propanamidoacetic acid | 161 | ND | ND | ND | ND | ND | ND |
| 11-Hydroxyundecanoic acid | 162 | ND | ND | ND | ND | ND | ND |
| 1-(2-Hydroxyethyl)-2,2,6,6-tetramethyl-4-piperidinol | 163 | ND | 140 | ND | ND | ND | ND |
| CE 16:1 | 164 | ND | ND | ND | ND | ND | ND |
| PC 41:6 | 165 | ND | ND | ND | ND | ND | 573 |
| DG 38:3 | 166 | ND | ND | ND | ND | ND | ND |
| LPC 14:0/0:0 | 167 | ND | ND | ND | ND | ND | ND |
| Cholesterol 3-sulfate | 168 | ND | ND | ND | ND | ND | ND |
| (S)-2-Ureidopentanedioic acid | 169 | ND | ND | ND | ND | ND | ND |
| 1-Palmitoyl-2-docosahexaenoyl-sn-glycero-3-phosphocholine | 170 | ND | ND | ND | ND | ND | 72 |
| 3[(2-Methylbenzyl)sulfanyl]-1H-1,2,4-triazol-5-ylamine | 171 | ND | ND | ND | ND | ND | ND |

| | | | | | | | |
|---|---|---|---|---|---|---|---|
| PC 33:2 | 172 | 2 | ND | ND | 551 | 468 | 522 |
| FA 15:0 | 173 | ND | ND | ND | 537 | ND | 68 |
| TG 62:13 \| TG 18:1_22:6_22:6 | 174 | ND | ND | 696 | ND | 895 | ND |
| PE P-36:4 \| PE P-16:0_20:4 | 175 | ND | ND | ND | ND | ND | ND |
| DG 36:5 | 176 | ND | 152 | 29 | 542 | 789 | ND |
| Ile-Leu | 177 | ND | ND | ND | ND | ND | 375 |
| 1-Stearoyl-2-arachidonyl-sn-glycero-3-phosphocholine | 178 | ND | ND | ND | ND | ND | ND |
| Hexuronic acid | 179 | ND | ND | ND | ND | ND | ND |
| 1,2-Diamino-2-methylpropane_1,2-Diamino-2-methylpropane | 180 | ND | ND | ND | ND | ND | ND |
| 2,6-Diaminopimelic acid | 181 | ND | ND | ND | ND | ND | 71 |
| Pentaethylene glycol | 182 | ND | ND | ND | ND | ND | ND |
| PC 35:5 | 183 | ND | ND | ND | ND | ND | ND |
| 1-Methylnicotinamide | 184 | 155 | 100 | 195 | ND | 95 | 54 |
| Equol | 185 | ND | ND | ND | 227 | ND | ND |
| FA 24:0 | 186 | 165 | ND | ND | 27 | ND | 898 |
| 1-Oleoyl-sn-glycero-3-phosphoethanolamine | 187 | ND | ND | ND | ND | 179 | 48 |
| 2-Amino-3-(dimethylamino)propionic acid | 188 | ND | ND | ND | ND | ND | ND |
| 3-Aminopyridine | 189 | ND | 175 | 68 | ND | 56 | ND |
| 4-(1,3-Benzothiazol-2-yl)butanoic acid | 190 | ND | ND | ND | ND | ND | ND |
| ST 28:1;O;S | 191 | ND | ND | ND | ND | ND | ND |
| 1-Palmitoyl-sn-glycero-3-phosphocholine | 192 | ND | ND | ND | ND | ND | ND |
| 2-Hydroxyphenylacetic acid | 193 | ND | ND | ND | ND | ND | ND |
| PI 40:5 \| PI 18:0_22:5 | 194 | ND | ND | 494 | ND | 685 | ND |
| FA 25:0_a | 195 | ND | ND | ND | ND | ND | ND |
| 2-Aminoadipic acid | 196 | ND | ND | ND | ND | ND | ND |
| TG 58:10 \| TG 18:2_18:2_22:6 | 197 | ND | ND | 663 | ND | ND | ND |
| 10-Hydroxydecanoic acid | 198 | ND | ND | ND | ND | ND | ND |
| (+)-Muscarine | 199 | ND | ND | ND | ND | ND | ND |
| Phenylpyruvic acid | 200 | ND | ND | ND | ND | ND | ND |
| PC 34:2;O \| PC 16:0_18:2;O_a | 201 | ND | ND | ND | ND | ND | ND |
| Sarcosine | 202 | ND | ND | ND | ND | 756 | ND |
| PC O-42:7 | 203 | ND | ND | ND | ND | ND | ND |
| 3-Indoleacetic acid | 204 | ND | ND | ND | ND | ND | ND |
| 1,4-Dihydroxy-2-naphthoic acid | 205 | ND | ND | ND | ND | ND | ND |
| PE O-38:6 \| PE O-18:2_20:4 | 206 | ND | ND | 496 | ND | ND | ND |
| 2-Methylglutamic acid | 207 | ND | ND | ND | ND | 255 | ND |
| 2-(3-Methylbenzyl)butanedioic acid | 208 | ND | ND | ND | ND | ND | ND |
| 2-Hydroxybenzaldehyde | 209 | ND | ND | ND | ND | ND | ND |
| 1-Palmitoyl-2-hydroxy-sn-glycero-3-phosphoethanolamine | 210 | ND | ND | ND | ND | 236 | ND |
| Lactobionic acid | 211 | ND | ND | ND | ND | ND | ND |
| Gulono-1,4-lactone | 212 | ND | ND | ND | ND | ND | ND |
| 2-Aminocaprylic acid | 213 | ND | ND | ND | ND | ND | ND |
| Erythronolactone | 214 | ND | ND | ND | ND | ND | ND |

| | | | | | | | |
|---|---|---|---|---|---|---|---|
| 1-Hexadecylpyridinium | 215 | ND | ND | 102 | ND | ND | ND |
| 1-Deoxynojirimycin | 216 | ND | ND | ND | ND | ND | ND |
| 2,3-Dihydroxy-2-methylbutanoic acid | 217 | ND | ND | ND | ND | ND | ND |
| Butyrylcholine | 218 | ND | ND | ND | ND | ND | ND |
| 3-Hydroxy-2-((9Z,12Z)-octadeca-9,12-dienoyloxy)propyl 2-(trimethylazaniumyl)ethyl phosphate | 219 | ND | ND | ND | ND | ND | 8 |
| Xanthurenic acid | 220 | ND | ND | ND | ND | ND | ND |
| Turanose | 221 | ND | ND | ND | ND | ND | ND |
| 1-Methylguanine | 222 | ND | ND | ND | ND | ND | ND |
| 2,2-Dimethyl-2,3-dihydro-4H-1,3-benzoxazin-4-one | 223 | ND | ND | ND | ND | ND | ND |
| Carnitine | 224 | 474 | 331 | 565 | 133 | 231 | 600 |
| 2-Phosphoglyceric acid | 225 | ND | ND | ND | ND | ND | ND |
| 2-Deoxyuridine | 226 | ND | ND | ND | ND | ND | ND |
| 3-Methyl-2-oxovaleric acid | 227 | ND | ND | ND | ND | ND | ND |
| Cytosine | 228 | 320 | ND | 680 | 131 | 190 | 460 |
| Cer 41:1;2O\|Cer 18:1;2O/23:0 | 229 | ND | ND | ND | ND | ND | ND |
| Acetaminophen sulfate | 230 | ND | ND | ND | ND | ND | ND |
| 2-Amino-3-methoxybenzoic acid | 231 | ND | ND | ND | ND | ND | ND |
| PC O-33:2 | 232 | ND | ND | ND | ND | ND | ND |
| Dihydrocapsaicin | 233 | ND | ND | ND | ND | ND | ND |
| 1,3-Dimethylbenzene-1-sulfonic acid | 234 | ND | ND | ND | ND | ND | ND |
| Pantothenic acid | 235 | 187 | 1 | 42 | 423 | 91 | ND |
| PC 35:5\|PC 15:1_20:4 | 236 | ND | ND | ND | ND | ND | ND |
| 3,3-Dimethylacrylic acid | 237 | ND | ND | ND | ND | ND | ND |
| LPC O-20:0 | 238 | ND | ND | ND | ND | ND | ND |
| 1-Acetylimidazole | 239 | 123 | ND | ND | ND | ND | ND |
| Val-Gln | 240 | ND | ND | ND | ND | 958 | ND |
| Alanine | 241 | ND | 353 | 336 | ND | ND | ND |
| O1_PC p-36:4; or PC o-36:5 | 242 | ND | ND | ND | ND | ND | ND |
| PI 38:5\|PI 18:0_20:5 | 243 | ND | ND | 493 | ND | ND | ND |
| PC 33:2\|PC 15:0_18:2 | 244 | ND | ND | ND | ND | ND | ND |
| 2-(4-Morpholinyl)-1-phenylethanol | 245 | ND | ND | ND | ND | ND | ND |
| 4-Hydroxyphenyllactic acid | 246 | ND | ND | ND | ND | ND | ND |
| 1,3-Cyclohexanedicarboxylic acid | 247 | ND | ND | ND | ND | ND | ND |
| Carnosine | 248 | 180 | ND | 158 | ND | 130 | 660 |
| 1-Methyl-L-histidine | 249 | ND | ND | ND | ND | ND | ND |
| N-Methylalanine | 250 | ND | 203 | 367 | 76 | ND | ND |
| 5-Hydroxymethyl-6-methyluracil | 251 | ND | ND | ND | ND | ND | ND |
| N,N-Bis(2-hydroxyethyl)dodecanamide | 252 | 89 | 172 | 179 | ND | ND | ND |
| Undecanedioic acid | 253 | ND | ND | ND | ND | ND | ND |
| FA 16:2 | 254 | ND | ND | 454 | ND | 813 | ND |
| 2-Deoxyribose | 255 | ND | ND | ND | ND | ND | ND |
| 2-dGMP | 256 | 154 | ND | ND | ND | ND | ND |
| Citraconic acid | 257 | ND | ND | ND | ND | ND | ND |

| | | | | | | | |
|---|---|---|---|---|---|---|---|
| TG 50:0 \| TG 16:0_16:0_18:0 | 258 | ND | ND | ND | ND | ND | ND |
| 3'-O-Methylcytidine | 259 | ND | ND | ND | ND | ND | ND |
| 2-Methylbutyryl-L-carnitine | 260 | ND | 187 | ND | ND | ND | ND |
| 2-(9-Decenyl)glutaconic acid | 261 | ND | ND | ND | ND | ND | ND |
| Acetyl-beta-alanine | 262 | ND | ND | ND | ND | ND | ND |
| 2'-O-Methyluridine | 263 | ND | ND | ND | ND | ND | ND |
| 4-Hydroxyhippuric acid | 264 | 48 | ND | ND | ND | ND | ND |
| 2-Methoxy-5-nitrophenol | 265 | ND | ND | ND | ND | ND | ND |
| 5-Hydroxytryptophan | 266 | ND | ND | ND | ND | ND | 239 |
| Abscisic acid | 267 | ND | ND | ND | ND | ND | ND |
| Arabitol | 268 | ND | ND | ND | ND | ND | ND |
| 3-Oxostearic acid | 269 | ND | ND | ND | ND | ND | ND |
| 4-Hydroxy-6-methyl-2-pyrone | 270 | ND | ND | ND | ND | ND | ND |
| FA 18:2;2O | 271 | ND | ND | ND | ND | ND | ND |
| 8-Hydroxyquinoline-2-carbaldehyde | 272 | ND | ND | ND | ND | ND | ND |
| Bradykinin | 273 | ND | ND | ND | ND | ND | ND |
| HexCer 40:1;2O \| HexCer 18:1;2O/22:0 | 274 | ND | ND | 287 | ND | ND | ND |
| 2-Isopropylmalic acid | 275 | ND | ND | ND | ND | ND | ND |
| 3-Hydroxy-3-methyl-2,3-dihydro-1H-indol-2-one | 276 | ND | ND | ND | ND | ND | ND |
| 5-Butyl-1H-pyrazole-3-carboxylic acid | 277 | ND | ND | ND | ND | ND | ND |
| Glucosamine | 278 | ND | ND | ND | ND | ND | ND |
| 2-Aminoisobutyric acid | 279 | ND | ND | ND | ND | ND | ND |
| 3-Hydroxyoctadecanoic acid | 280 | ND | ND | ND | ND | ND | ND |
| LPE 16:0 | 281 | ND | ND | 171 | 365 | 1020 | 401 |
| Betaine aldehyde | 282 | 140 | 218 | 744 | ND | 311 | ND |
| LPC 20:0/0:0 | 283 | ND | ND | ND | ND | ND | ND |
| TG 58:12 \| TG 18:2_18:4_22:6 | 284 | ND | ND | 665 | ND | 865 | ND |
| Cer 34:1;2O \| Cer 18:1;2O/16:0 | 285 | ND | ND | ND | ND | ND | ND |
| Galactosamine-1-phosphate | 286 | ND | ND | ND | ND | ND | ND |
| Indoxyl sulfate | 287 | ND | ND | ND | ND | ND | ND |
| 1,5-Anhydrosorbitol | 288 | ND | ND | ND | ND | ND | ND |
| 1-Palmitoyl-2-azelaoylphosphatidylcholine | 289 | ND | ND | ND | ND | ND | 917 |
| 3-Methylhistidine | 290 | 83 | ND | ND | ND | 213 | 61 |
| 1,4-Butynediol | 291 | ND | ND | ND | ND | ND | ND |
| DG 32:0 | 292 | 293 | 200 | ND | 83 | ND | 455 |
| 3-Propyl-1H-pyrazole-5-carboxylic acid | 293 | ND | ND | ND | ND | ND | ND |
| PC 42:2 | 294 | ND | ND | ND | ND | 519 | ND |
| FA 20:4;O | 295 | ND | ND | ND | ND | ND | ND |
| 2-Hydroxyisobutyric acid | 296 | ND | ND | ND | ND | ND | ND |
| 3-Carboxy-6-methylcoumarin | 297 | ND | ND | ND | ND | ND | ND |
| TG 52:0 \| TG 16:0_18:0_18:0 | 298 | ND | ND | 61 | ND | ND | ND |
| 2-{2-[2-(Methacryloyloxy)ethoxy]ethoxy}ethyl 2-methylacrylate | 299 | ND | 165 | ND | ND | 221 | ND |
| Glutamic acid | 300 | 238 | 334 | 316 | 353 | ND | ND |

| | | | | | | | |
|---|---|---|---|---|---|---|---|
| FA 24:6 | 301 | ND | ND | 261 | ND | 133 | ND |
| SL 32:0;0O | 302 | ND | ND | ND | ND | ND | ND |
| Linoleic acid | 303 | ND | ND | ND | ND | ND | ND |
| 6-Methyl-2-[(2-oxo-2-phenylethyl)sulfanyl]-4(3H)-pyrimidinone | 304 | ND | ND | ND | ND | ND | ND |
| 2'-Deoxycytidine | 305 | 65 | ND | ND | ND | 116 | 916 |
| Lys-Ala | 306 | 68 | ND | ND | ND | ND | 420 |
| 2-(4-Fluorophenyl)acetohydrazide | 307 | ND | ND | ND | ND | ND | ND |
| FA 10:0_b | 308 | ND | ND | ND | ND | ND | ND |
| TG 60:10 \| TG 18:1_20:3_22:6 | 309 | ND | ND | ND | ND | ND | ND |
| 1-Stearoyl-2-hydroxy-sn-glycero-3-phosphocholine | 310 | ND | ND | ND | ND | ND | 102 |
| N,N-Dimethylarginine | 311 | 556 | 371 | 351 | 403 | 265 | 432 |
| Pyroglutamic acid | 312 | ND | 392 | 533 | 449 | ND | 650 |
| Homovanillic acid sulfate | 313 | ND | ND | ND | ND | ND | ND |
| Ala-Ala | 314 | 111 | ND | ND | ND | ND | 864 |
| Ala-Glu | 315 | 317 | ND | ND | ND | ND | 110 |
| 3-Aminoisobutyric acid | 316 | ND | ND | ND | ND | ND | ND |
| Bisphenol A bis(2,3-dihydroxypropyl) ether | 317 | ND | ND | ND | ND | ND | ND |
| erythro-Sphingosine-1-phosphate | 318 | ND | ND | ND | ND | ND | ND |
| 1,2-Dimethylimidazole | 319 | ND | ND | ND | ND | 206 | ND |
| Sorbose | 320 | ND | ND | ND | ND | ND | ND |
| 3-Phenyllactic acid | 321 | ND | ND | ND | ND | ND | ND |
| Glutathione (reduced) | 322 | 348 | ND | ND | 124 | ND | 278 |
| 2-(1H-Pyrazol-1-yl)benzylamine | 323 | ND | ND | ND | ND | ND | ND |
| 3-Oxocholic acid | 324 | ND | ND | ND | ND | ND | ND |
| Tuberonic acid | 325 | ND | ND | ND | ND | ND | ND |
| 3,3-Dimethylglutaric acid | 326 | ND | ND | ND | ND | ND | ND |
| 3-Hydroxydodecanoic acid | 327 | ND | ND | ND | ND | ND | 74 |
| Serotonin | 328 | 444 | ND | ND | ND | ND | ND |
| FA 26:0;O | 329 | ND | ND | ND | ND | ND | ND |
| FA 22:2 | 330 | 512 | ND | ND | 308 | ND | 56 |
| 2-Hydroxyquinoline | 331 | ND | ND | ND | ND | ND | ND |
| TG 46:2 \| TG 12:0_16:0_18:2 | 332 | ND | ND | ND | ND | ND | ND |
| PC O-36:3 | 333 | ND | ND | ND | ND | 535 | ND |
| 2-Oleoyl-1-palmitoyl-sn-glycero-3-phosphocholine | 334 | ND | ND | ND | ND | ND | ND |
| SM 34:1;2O \| SM 18:1;2O/16:0 | 335 | ND | ND | ND | ND | ND | ND |
| DG 36:4 | 336 | 23 | 48 | ND | 348 | 788 | 338 |
| 5-Hydroxyvalproic acid | 337 | ND | ND | ND | ND | ND | ND |
| FA 20:2 | 338 | 466 | ND | ND | 264 | ND | 909 |
| LPC 20:1/0:0 | 339 | ND | ND | ND | ND | ND | ND |
| N-Acetylalanine | 340 | ND | ND | ND | ND | ND | ND |
| 3-Hydroxysebacic acid | 341 | ND | ND | ND | ND | ND | ND |
| LPC 15:0/0:0 | 342 | ND | ND | ND | ND | ND | ND |
| PC 37:5;O \| PC 20:4_17:1;O | 343 | ND | ND | ND | ND | ND | ND |

| | | | | | | | |
|---|---|---|---|---|---|---|---|
| 1,2-dioleoyl-sn-glycero-3-phosphatidylcholine_1,2-dioleoyl-sn-glycero-3-phosphatidylcholine | 344 | ND | ND | ND | ND | ND | ND |
| FA 24:0;O | 345 | ND | ND | ND | ND | ND | ND |
| PC O-40:6 | 346 | ND | ND | ND | ND | 284 | ND |
| 2-(Formylamino)benzoic acid | 347 | ND | ND | ND | ND | ND | ND |
| PC O-38:4 | 348 | ND | ND | ND | ND | 540 | ND |
| 3-Galloylgallocatechin | 349 | ND | ND | ND | ND | ND | ND |
| Cholesterol | 350 | 36 | ND | ND | 210 | 371 | 227 |
| Betaine | 351 | 363 | 291 | 341 | 92 | 77 | 151 |
| 3-Hydroxyanthranilic acid | 352 | ND | ND | ND | ND | ND | ND |
| 4-Fluorobenzene-1,3-dicarboxylic acid | 353 | ND | ND | ND | ND | ND | ND |
| Homovanillic acid | 354 | ND | ND | ND | ND | ND | ND |
| 4-Hydroxy-6-methylnicotinic acid | 355 | ND | ND | ND | ND | ND | ND |
| DG 34:2 | 356 | 361 | 66 | ND | 380 | 346 | 452 |
| PC 36:4;3O\|PC 18:2_18:2;3O | 357 | ND | ND | ND | ND | ND | ND |
| gamma-Glutamyl-alanine | 358 | ND | ND | ND | ND | ND | ND |
| LPC 22:0/0:0 | 359 | ND | ND | ND | ND | ND | ND |
| 7.125_186.1124 | 360 | ND | ND | ND | ND | ND | ND |
| 4-Hydroxybenzoic acid | 361 | ND | ND | ND | ND | ND | ND |
| 6-Hydroxycaproic acid | 362 | ND | ND | ND | ND | ND | ND |
| Isoquinoline-3-carboxylic acid | 363 | ND | ND | ND | ND | ND | ND |
| Pipecolic acid | 364 | 225 | ND | ND | ND | ND | ND |
| N-Methylglutamic acid | 365 | 77 | ND | ND | ND | ND | 447 |
| Spermine | 366 | ND | ND | ND | ND | ND | ND |
| beta-Homoglutamine | 367 | ND | ND | ND | ND | 76 | ND |
| LPC O-18:0 | 368 | ND | ND | ND | ND | ND | ND |
| DG 34:1 | 369 | 434 | 73 | ND | 4 | 19 | 453 |
| Ethylmalonic acid | 370 | ND | ND | ND | ND | ND | ND |
| Hexaethylene glycol | 371 | ND | 189 | 238 | ND | 390 | ND |
| 3,5-Dichlorosalicylic acid | 372 | ND | ND | ND | ND | ND | ND |
| TG O-55:9\|TG O-13:1_21:4_21:4 | 373 | ND | ND | ND | ND | ND | ND |
| 4-Amino-2-methylbenzamide | 374 | ND | ND | ND | ND | ND | ND |
| 8-(3-Octyl-2-oxiranyl)octanoic acid | 375 | ND | ND | ND | ND | ND | ND |
| LPC 23:0/0:0 | 376 | ND | ND | ND | ND | ND | ND |
| Cer 42:0;2O\|Cer 18:0;2O/24:0 | 377 | ND | ND | ND | ND | ND | 266 |
| SM 32:1;2O | 378 | ND | ND | ND | ND | ND | ND |
| Oleoyl ethylamide | 379 | ND | ND | ND | ND | ND | ND |
| 3-Methylcrotonylglycine | 380 | 335 | ND | ND | ND | ND | ND |
| Propionylcarnitine | 381 | 200 | ND | ND | 426 | 411 | 631 |
| Mannitol | 382 | ND | ND | ND | ND | ND | ND |
| Adenine | 383 | 215 | ND | ND | ND | ND | 195 |
| PC 39:6 | 384 | 422 | ND | ND | 190 | 82 | 558 |
| PC O-40:7 | 385 | ND | ND | ND | ND | 349 | ND |
| Guanidinoacetic acid | 386 | ND | ND | ND | ND | ND | 323 |

| | | | | | | | |
|---|---|---|---|---|---|---|---|
| PI 38:3 \| PI 18:0_20:3 | 387 | ND | ND | 491 | ND | ND | ND |
| 2-Piperidinecarboxamide | 388 | ND | ND | 724 | ND | ND | ND |
| 3-Methyladipic acid | 389 | ND | ND | ND | ND | ND | ND |
| N-(2-Furoyl)glycine | 390 | 213 | ND | ND | ND | ND | 430 |
| 4-Hydroxyquinoline | 391 | 236 | ND | ND | 98 | ND | ND |
| O2_FA 24:1; (nervonic acid) | 392 | ND | ND | ND | ND | ND | ND |
| PI 39:4 \| PI 19:0_20:4 | 393 | ND | ND | ND | ND | 41 | ND |
| N-Acetylcytidine | 394 | ND | ND | ND | ND | ND | 262 |
| PE 34:2 | 395 | ND | ND | ND | 493 | 90 | 162 |
| Methyl 4-hydroxycinnamate | 396 | ND | ND | ND | ND | ND | ND |
| 4-Isoxazolepropanoic acid, alpha-amino-2,3-dihydro-5-methyl-3-oxo- | 397 | ND | ND | ND | ND | ND | ND |
| FA 21:5 | 398 | ND | ND | 374 | ND | 901 | ND |
| N-acetyltryptophan | 399 | ND | ND | ND | ND | ND | ND |
| 2-Acetylpyrazine | 400 | ND | ND | ND | ND | 205 | ND |
| Cysteic Acid | 401 | ND | ND | ND | ND | ND | ND |
| SM 40:2;2O \| SM 15:1;2O/25:1 | 402 | ND | ND | ND | ND | ND | ND |
| saccharic acid | 403 | ND | ND | ND | 273 | ND | ND |
| 2(1H)-Pyridinone | 404 | ND | ND | ND | ND | ND | ND |
| P-Coumaric acid | 405 | ND | ND | ND | ND | ND | ND |
| Benzeneethanamine, 3,5-dimethoxy-.alpha.-methyl-4-propoxy- | 406 | ND | ND | ND | ND | ND | ND |
| epsilon-Dimethyl-lysine | 407 | ND | ND | ND | ND | ND | ND |
| Acamprosate | 408 | ND | ND | ND | ND | ND | ND |
| PC O-38:7 | 409 | ND | ND | ND | ND | 545 | ND |
| 3-Hydroxyoctanoic acid | 410 | ND | ND | ND | ND | ND | ND |
| SE 28:1/20:3 | 411 | ND | ND | ND | ND | ND | ND |
| 3'-O-Methylguanosine | 412 | 130 | ND | ND | ND | ND | ND |
| 5-Methylcytidine | 413 | ND | ND | ND | ND | ND | ND |
| TG 60:12 \| TG 18:1_20:5_22:6 | 414 | ND | ND | ND | ND | ND | ND |
| LPC O-16:0 | 415 | ND | ND | ND | ND | 334 | ND |
| N-Glycolylneuraminic acid | 416 | ND | ND | ND | ND | ND | ND |
| 3-Hydroxypropanoic acid | 417 | ND | ND | ND | ND | ND | ND |
| 2,8-Quinolinediol | 418 | ND | ND | ND | ND | ND | ND |
| PE 40:6 \| PE 18:0_22:6 | 419 | ND | ND | 479 | ND | 588 | ND |
| O1_FA 20:4; (arachidonic acid) | 420 | ND | ND | ND | ND | ND | ND |
| Guanidinopropionic acid | 421 | ND | ND | ND | ND | ND | ND |
| LPC 24:0/0:0 | 422 | ND | ND | ND | ND | ND | ND |
| 3-(Trifluoromethyl)cinnamic acid | 423 | ND | ND | ND | ND | ND | ND |
| SM 42:1;2O \| SM 18:1;2O/24:0 | 424 | ND | ND | ND | ND | ND | ND |
| Cystine | 425 | 462 | ND | 679 | 362 | ND | 542 |
| Isoxanthopterin | 426 | ND | ND | ND | ND | ND | ND |
| TG 58:12 \| TG 18:2_20:5_20:5 | 427 | ND | ND | ND | ND | ND | ND |
| 3-Fluoro-5-(methoxycarbonyl)benzoic acid | 428 | ND | ND | ND | ND | ND | ND |
| PC O-41:11 | 429 | ND | ND | ND | ND | ND | ND |

| | | | | | | | |
|---|---|---|---|---|---|---|---|
| Creatine | 430 | 239 | 228 | 199 | 313 | 277 | 545 |
| DG 34:2 \| DG 16:0_18:2 | 431 | ND | ND | 237 | ND | ND | ND |
| PC O-31:7 | 432 | ND | ND | ND | ND | ND | ND |
| N-Acetylneuraminic acid | 433 | 338 | ND | ND | ND | ND | 461 |
| CAR 4:0 | 434 | ND | ND | ND | ND | 682 | ND |
| 4-Hydroxybenzoic acid propyl ester | 435 | ND | ND | ND | ND | ND | ND |
| Citrulline | 436 | 443 | 359 | 677 | 316 | 347 | 546 |
| Palmitoyl sphingomyelin | 437 | ND | ND | ND | ND | ND | ND |
| Benzyldimethylstearylammonium | 438 | 403 | ND | 126 | ND | ND | ND |
| PC 40:4 \| PC 20:0_20:4 | 439 | ND | ND | ND | ND | ND | ND |
| FA 10:0_a | 440 | ND | ND | ND | ND | ND | ND |
| N-Methylproline | 441 | ND | ND | ND | ND | ND | ND |
| Cysteinesulfinic acid | 442 | 515 | ND | ND | ND | ND | ND |
| 5-Methyl-5,6-Dihydrouracil | 443 | ND | ND | ND | ND | ND | ND |
| Glycerophosphocholine | 444 | ND | 279 | 319 | 63 | 308 | 325 |
| Cer 43:1;2O \| Cer 19:1;2O/24:0 | 445 | ND | ND | ND | ND | ND | ND |
| 3,4-Dihydro-3-oxo-2H-(1,4)-benzoxazin-2-ylacetic acid | 446 | ND | ND | ND | ND | ND | ND |
| NAE 14:0 | 447 | ND | ND | ND | ND | ND | ND |
| Docosan-1-amine | 448 | ND | ND | 133 | ND | ND | ND |
| Cysteine-glutathione disulfide | 449 | 425 | ND | ND | ND | ND | ND |
| Glyceraldehyde | 450 | ND | ND | ND | ND | ND | ND |
| O4_FA 20:4; (arachidonic acid) | 451 | ND | ND | ND | ND | ND | ND |
| CAR 12:0 | 452 | ND | ND | ND | ND | 120 | ND |
| PC 38:4 \| PC 18:0_20:4 | 453 | ND | ND | ND | ND | ND | ND |
| SM 36:2;2O \| SM 19:1;2O/17:1 | 454 | ND | ND | ND | ND | ND | ND |
| Phenylacetylglycine | 455 | ND | ND | ND | ND | ND | ND |
| Cinnamic acid | 456 | ND | ND | ND | ND | ND | ND |
| 5(Aminocarbonyl)amino]pentanoic acid | 457 | ND | ND | ND | ND | ND | ND |
| Cholic acid | 458 | ND | ND | ND | ND | ND | ND |
| Asp-Glu | 459 | ND | ND | ND | ND | ND | 232 |
| PC 32:2 | 460 | 96 | ND | 407 | 549 | 465 | 523 |
| Cer 41:0;2O \| Cer 18:0;2O/23:0 | 461 | ND | ND | ND | ND | ND | ND |
| 2-Methyl-4-phenylazophenylamine | 462 | ND | ND | ND | ND | ND | ND |
| SM 36:2;2O \| SM 18:2;2O/18:0 | 463 | ND | ND | ND | ND | ND | ND |
| Dodecanedioic acid | 464 | ND | ND | ND | ND | ND | ND |
| CE 18:3 | 465 | ND | ND | ND | 579 | ND | ND |
| Glyceric acid | 466 | ND | ND | ND | ND | ND | ND |
| Arginine | 467 | 366 | 380 | 338 | ND | 67 | ND |
| FA 18:3 | 468 | 52 | ND | ND | 370 | ND | 923 |
| glycerol-alpha-phosphate | 469 | 174 | ND | ND | 298 | 1012 | 891 |
| TG 55:7 \| TG 16:0_18:2_21:5 | 470 | ND | ND | 635 | ND | 843 | 762 |
| Hypoxanthine | 471 | ND | 317 | 274 | 427 | 983 | 378 |
| Ser-Val | 472 | ND | ND | ND | ND | ND | ND |

| | | | | | | | |
|---|---|---|---|---|---|---|---|
| FA 15:1 | 473 | 160 | ND | ND | 163 | ND | 935 |
| N,N-Dimethylformamide | 474 | 460 | ND | ND | ND | ND | ND |
| 5-Methoxy-3-indoleacetic acid | 475 | ND | ND | ND | ND | ND | ND |
| Caffeic acid | 476 | ND | ND | ND | ND | ND | ND |
| N-Methyl-L-asparagine | 477 | ND | ND | 366 | ND | ND | ND |
| Gly-Val | 478 | ND | ND | ND | ND | 933 | ND |
| PC 32:2\|PC 14:0_18:2 | 479 | ND | ND | ND | ND | ND | ND |
| Hexadecanedioic acid | 480 | ND | ND | ND | ND | ND | ND |
| N-Acetylcysteine | 481 | ND | ND | ND | ND | ND | ND |
| 4-Methylumbelliferone | 482 | ND | ND | ND | ND | ND | ND |
| 7,8-Dihydrobiopterin | 483 | ND | ND | ND | ND | ND | ND |
| Histidine | 484 | 253 | 385 | 307 | ND | ND | 379 |
| Homogentisic acid | 485 | ND | ND | ND | ND | ND | ND |
| 2,3-Dimethoxy-5-methylbenzoquinone | 486 | ND | ND | ND | ND | ND | ND |
| 7-Nitro-2,1,3-benzoxadiazol-4-ylamine | 487 | ND | ND | ND | ND | ND | ND |
| Cysteine S-sulfate | 488 | ND | ND | ND | ND | ND | ND |
| Triethanolamine | 489 | ND | 132 | 259 | ND | ND | ND |
| Aleuritic acid | 490 | ND | ND | ND | ND | ND | ND |
| LPC 18:0 | 491 | 133 | ND | 206 | 237 | 405 | 408 |
| N-Methylhistidine | 492 | 299 | ND | 368 | ND | ND | ND |
| 3-Ureidopropionic acid | 493 | ND | ND | ND | ND | ND | ND |
| Octadecanedioic acid | 494 | ND | ND | ND | ND | ND | 509 |
| 3-Acetoxypyridine | 495 | ND | ND | 72 | ND | ND | ND |
| Theanine | 496 | ND | ND | ND | ND | ND | ND |
| PE 38:6\|PE 16:0_22:6 | 497 | ND | ND | 446 | ND | ND | ND |
| FA 21:0 | 498 | ND | ND | ND | ND | ND | ND |
| Malonyl-L-carnitine | 499 | ND | ND | ND | ND | ND | 414 |
| PC O-34:3 | 500 | ND | ND | ND | ND | 533 | ND |
| 5-Hydroxy-3,4-dihydro-2(1H)-quinolinone | 501 | ND | ND | ND | ND | ND | ND |
| Pro-Leu | 502 | 534 | ND | ND | ND | 714 | ND |
| N-Tigloylglycine | 503 | ND | ND | ND | ND | ND | ND |
| SM 32:1;2O\|SM 18:1;2O/14:0 | 504 | ND | ND | ND | ND | ND | ND |
| CAR 13:0 | 505 | ND | ND | ND | ND | ND | ND |
| Malic acid | 506 | ND | ND | ND | ND | ND | ND |
| FA 14:1 | 507 | 132 | ND | ND | 249 | 245 | 936 |
| N8-Acetylspermidine | 508 | 38 | ND | ND | ND | ND | 440 |
| LPI 18:0 | 509 | ND | ND | 303 | ND | 1044 | ND |
| PE 38:5\|PE 18:1_20:4 | 510 | ND | ND | 121 | ND | ND | ND |
| Adenosine | 511 | ND | ND | ND | 62 | 310 | 146 |
| Abietic acid | 512 | ND | ND | ND | ND | ND | ND |
| Thymidine | 513 | ND | ND | ND | ND | ND | ND |
| SM 40:1;2O\|SM 18:1;2O/22:0 | 514 | ND | ND | ND | ND | ND | ND |
| LPC 20:4/0:0 | 515 | ND | ND | ND | ND | ND | ND |

| | | | | | | | |
|---|---|---|---|---|---|---|---|
| Thiamine | 516 | 566 | ND | 131 | ND | 946 | ND |
| Homoarginine | 517 | ND | ND | ND | ND | ND | ND |
| Pro-Gly | 518 | 539 | ND | ND | ND | 713 | ND |
| Dihydro-4,4-dimethyl-2,3-furandione | 519 | ND | ND | ND | ND | ND | ND |
| Imidazole | 520 | 379 | ND | 252 | ND | ND | ND |
| FA 16:1 | 521 | 167 | ND | ND | 539 | ND | 933 |
| Prolylphenylalanine | 522 | ND | ND | ND | ND | ND | ND |
| FA 18:3;O | 523 | ND | ND | ND | ND | ND | ND |
| Gentisinic acid | 524 | ND | ND | ND | ND | ND | ND |
| P-Toluenesulfonic acid | 525 | ND | ND | ND | ND | ND | ND |
| FA 18:4 | 526 | ND | ND | 178 | ND | 869 | ND |
| 4-Acetamidobutyric acid | 527 | 124 | ND | ND | ND | ND | ND |
| Resveratrol-3-O-sulfate | 528 | ND | ND | ND | ND | ND | ND |
| FA 18:0;O | 529 | ND | ND | ND | ND | ND | ND |
| Cer 40:0;2O \| Cer 18:0;2O/22:0 | 530 | ND | ND | ND | ND | ND | 657 |
| SM 41:2;2O | 531 | ND | ND | ND | ND | ND | ND |
| SM 33:1;2O | 532 | ND | ND | ND | ND | ND | ND |
| FA 28:7 | 533 | ND | ND | 8 | ND | 256 | ND |
| CAR 20:1 | 534 | ND | ND | ND | ND | 269 | ND |
| DG 36:5 \| DG 18:2_18:3 | 535 | ND | ND | ND | ND | ND | ND |
| 2-Hydroxy-5-methoxybenzoic acid | 536 | ND | ND | ND | ND | ND | ND |
| PI 38:5 \| PI 18:1_20:4 | 537 | ND | ND | 77 | ND | ND | ND |
| TG 58:7 \| TG 18:1_18:2_22:4 | 538 | ND | ND | ND | ND | ND | 787 |
| 4-Acetylbutyric acid | 539 | ND | ND | ND | ND | ND | ND |
| Isocitric acid lactone | 540 | ND | ND | ND | ND | ND | ND |
| LPC 18:2/0:0 | 541 | ND | ND | ND | ND | ND | ND |
| 3-Methylcytidine | 542 | 138 | ND | ND | ND | ND | ND |
| N-Acetyllysine | 543 | ND | ND | ND | ND | ND | ND |
| Orotic acid | 544 | ND | ND | ND | ND | ND | ND |
| Phenylalanine, methyl ester | 545 | ND | ND | ND | ND | 711 | ND |
| Guanine | 546 | ND | ND | ND | 318 | ND | 291 |
| 5-Hydroxy-3-indoleacetic acid | 547 | ND | ND | ND | ND | ND | ND |
| FA 18:2;O | 548 | ND | ND | ND | ND | ND | ND |
| DG 40:7 \| DG 18:1_22:6 | 549 | ND | ND | 432 | ND | 800 | ND |
| Cer 34:2;2O \| Cer 18:1;2O/16:1 | 550 | ND | ND | ND | ND | ND | ND |
| Triphenyl phosphate | 551 | ND | ND | ND | ND | ND | ND |
| Glutaric acid | 552 | ND | ND | ND | ND | ND | ND |
| Cyclohexanamine | 553 | ND | ND | ND | ND | ND | ND |
| Docosahexanoic acid | 554 | ND | ND | ND | ND | ND | ND |
| N-Acetylmethionine | 555 | ND | ND | 364 | ND | 1067 | ND |
| Adipic acid | 556 | ND | ND | ND | ND | ND | ND |
| Leu-Leu | 557 | ND | ND | ND | ND | ND | ND |
| Anserine | 558 | 311 | 322 | 337 | 88 | 585 | 113 |

| | | | | | | | |
|---|---|---|---|---|---|---|---|
| N-Acetyl-epsilon-caprolactam | 559 | ND | ND | ND | ND | ND | ND |
| CAR 6:0 | 560 | ND | ND | ND | ND | ND | ND |
| Indole-3-acetonitrile | 561 | ND | ND | ND | ND | ND | ND |
| Pseudouridine | 562 | ND | ND | ND | ND | ND | ND |
| PE 34:1 \| PE 16:0_18:1 | 563 | ND | ND | 462 | ND | 560 | ND |
| N-Acetylhistidine | 564 | 533 | 395 | 363 | ND | ND | ND |
| Ala-Arg | 565 | ND | ND | 335 | 67 | 412 | 822 |
| FA 20:3 | 566 | 519 | ND | ND | 180 | ND | 908 |
| FA 25:0_b | 567 | ND | ND | ND | ND | ND | ND |
| Picolinuric acid | 568 | ND | ND | ND | ND | ND | ND |
| Taurine | 569 | 184 | 379 | 283 | ND | 943 | ND |
| LPC O-18:1 | 570 | ND | ND | ND | ND | 1015 | ND |
| TG 55:8 \| TG 15:0_18:2_22:6 | 571 | ND | ND | ND | ND | 844 | ND |
| PC 38:6 | 572 | 266 | ND | 438 | 305 | 497 | 108 |
| PI 38:4;O \| PI 18:0_20:4;O | 573 | ND | ND | ND | ND | ND | ND |
| TG 62:12 \| TG 18:1_22:5_22:6 | 574 | ND | ND | 695 | ND | 894 | ND |
| Phosphocholine | 575 | 314 | ND | ND | 254 | ND | ND |
| TG 56:8 \| TG 16:0_18:2_22:6 | 576 | ND | ND | 647 | ND | ND | ND |
| Benzoylmalic acid | 577 | ND | ND | ND | ND | ND | ND |
| PC 40:6 \| PC 18:0_22:6 | 578 | ND | ND | ND | ND | ND | ND |
| Cer 42:2;2O \| Cer 18:1;2O/24:1 | 579 | ND | ND | ND | ND | ND | ND |
| LPC 24:1/0:0 | 580 | ND | ND | ND | ND | ND | ND |
| LPE O-18:2 | 581 | ND | ND | 114 | ND | 1039 | ND |
| TG 43:2;1O \| TG 10:0_16:0_17:2;1O | 582 | ND | ND | ND | ND | ND | ND |
| LPC 17:1/0:0 | 583 | ND | ND | ND | ND | ND | ND |
| Icaridin | 584 | ND | ND | ND | ND | ND | ND |
| Diisodecyl phthalate | 585 | ND | ND | ND | ND | ND | ND |
| PC 34:0 | 586 | 309 | ND | 417 | 531 | 384 | 521 |
| PC O-36:1 | 587 | ND | ND | ND | ND | ND | ND |
| Gly-Gly | 588 | ND | ND | ND | ND | ND | ND |
| LPC 20:5/0:0 | 589 | ND | ND | ND | ND | ND | ND |
| O2_PE 38:2 isomer b | 590 | ND | ND | ND | ND | ND | ND |
| FA 23:0 | 591 | ND | ND | ND | ND | ND | ND |
| Gabapentin | 592 | ND | ND | ND | ND | ND | ND |
| Hydroferulic acid | 593 | ND | ND | ND | ND | ND | ND |
| DG 50:1 | 594 | ND | ND | ND | ND | ND | ND |
| PC 39:4 \| PC 19:0_20:4 | 595 | ND | ND | ND | ND | ND | ND |
| O1_FA 20:5; (eicosapentaenoic acid) | 596 | ND | ND | ND | ND | ND | ND |
| N-Acetylmannosamine | 597 | ND | ND | ND | ND | ND | ND |
| Ala-Ile | 598 | 245 | ND | ND | ND | ND | ND |
| PC 36:2 | 599 | 290 | ND | 424 | 529 | 6 | 537 |
| PC O-36:6 | 600 | ND | ND | ND | ND | 177 | ND |
| Palitantin | 601 | ND | ND | ND | ND | ND | ND |

| | | | | | | | |
|---|---|---|---|---|---|---|---|
| 4-Hydroxyvalproic acid | 602 | ND | ND | ND | ND | ND | ND |
| O2_PE 38:2 | 603 | ND | ND | 241 | ND | ND | ND |
| LPC 18:3/0:0 | 604 | ND | ND | ND | ND | ND | ND |
| Cedrin | 605 | ND | ND | ND | ND | ND | ND |
| Ethyl sulfate | 606 | ND | ND | ND | ND | ND | ND |
| Zinterol | 607 | ND | ND | ND | ND | ND | ND |
| 4-Pyridoxic acid | 608 | ND | ND | ND | 26 | ND | ND |
| 5'-S-Methyl-5'-thioadenosine | 609 | ND | ND | ND | ND | ND | ND |
| FA 22:5;O | 610 | ND | ND | ND | ND | ND | ND |
| SM 35:2;3O | 611 | ND | ND | ND | ND | ND | ND |
| SM 42:3;2O | 612 | ND | ND | ND | ND | ND | ND |
| PC O-37:4 | 613 | ND | ND | ND | ND | ND | ND |
| Acetohydroxamic acid | 614 | ND | ND | ND | ND | ND | ND |
| Cer 33:1;2O\|Cer 17:1;2O/16:0 | 615 | ND | ND | ND | ND | ND | ND |
| SE 29:1/20:4 | 616 | ND | ND | ND | ND | ND | ND |
| PI 38:4\|PI 18:0_20:4 | 617 | ND | ND | 492 | ND | ND | ND |
| PI 36:2\|PI 18:0_18:2_a | 618 | ND | ND | ND | ND | ND | ND |
| Putrescine | 619 | ND | 110 | 504 | ND | ND | ND |
| Galactosamine | 620 | 525 | ND | ND | 198 | ND | 944 |
| Hexanoyl-L-carnitine | 621 | 569 | ND | ND | ND | 977 | 365 |
| LPC 22:6/0:0 | 622 | ND | ND | ND | ND | ND | ND |
| TG 47:0\|TG 15:0_16:0_16:0 | 623 | ND | ND | 198 | ND | 243 | ND |
| PC O-36:5\|PC O-16:1_20:4 | 624 | ND | ND | ND | ND | ND | ND |
| FA 20:5 | 625 | 42 | ND | ND | 196 | ND | 905 |
| Acetyl-carnitine | 626 | ND | ND | ND | ND | ND | ND |
| Hexanoylglycine | 627 | ND | ND | ND | ND | ND | ND |
| PC 32:0\|PC 16:0_16:0 | 628 | ND | ND | ND | ND | ND | ND |
| 4.94_805.56 | 629 | ND | ND | ND | ND | ND | ND |
| Epigallocatechin | 630 | 571 | ND | ND | ND | 396 | ND |
| Cer 39:1;2O\|Cer 17:1;2O/22:0 | 631 | ND | ND | ND | ND | ND | 284 |
| Serine | 632 | ND | 387 | 562 | 388 | 759 | 681 |
| LPE 22:6 | 633 | ND | ND | 51 | 351 | 1031 | ND |
| Lauroyl-L-carnitine | 634 | ND | ND | ND | ND | ND | ND |
| Methylacetate | 635 | ND | ND | ND | ND | ND | ND |
| Maleic acid | 636 | ND | ND | ND | ND | ND | ND |
| Normetanephrine | 637 | ND | ND | ND | ND | ND | ND |
| SM 38:2;2O\|SM 18:2;2O/20:0 | 638 | ND | ND | ND | ND | ND | ND |
| PC O-37:8 | 639 | ND | ND | ND | ND | 539 | ND |
| LPC 18:1 | 640 | 136 | ND | 286 | 543 | 996 | 407 |
| PE O-37:4\|PE O-21:3_16:1 | 641 | ND | ND | ND | ND | ND | ND |
| Ketoisovaleric acid | 642 | ND | ND | ND | ND | ND | ND |
| N-Cinnamoylglycine | 643 | ND | ND | ND | ND | ND | ND |
| PI 34:1\|PI 16:0_18:1 | 644 | ND | ND | ND | ND | ND | ND |

| | | | | | | | |
|---|---|---|---|---|---|---|---|
| Xanthosine | 645 | ND | ND | ND | 289 | ND | ND |
| Tetradecanedioic acid | 646 | ND | ND | ND | ND | ND | ND |
| FA 18:1;O | 647 | ND | ND | ND | ND | ND | ND |
| Propionic acid | 648 | ND | ND | ND | ND | ND | ND |
| PC O-38:2 | 649 | ND | ND | ND | ND | ND | ND |
| Malonic acid | 650 | ND | ND | ND | ND | ND | ND |
| phosphoethanolamine | 651 | 197 | ND | ND | 321 | 226 | 857 |
| SM 42:2;2O\|SM 19:1;2O/23:1 | 652 | ND | ND | ND | ND | ND | ND |
| SM 42:3;2O\|SM 14:1;2O/28:2 | 653 | ND | ND | ND | ND | ND | ND |
| Catechol | 654 | ND | ND | ND | ND | ND | ND |
| His-Pro | 655 | 380 | ND | 169 | 387 | 244 | 357 |
| TG 48:2\|TG 14:0_16:0_18:2 | 656 | ND | ND | ND | ND | ND | ND |
| SM 42:2;2O\|SM 18:1;2O/24:1 | 657 | ND | ND | ND | ND | ND | ND |
| Glufosinate | 658 | ND | ND | ND | ND | ND | ND |
| Indole-3-carboxaldehyde | 659 | ND | ND | ND | ND | ND | ND |
| PC O-32:1;1O\|PC O-17:0_15:1;1O | 660 | ND | ND | ND | ND | ND | ND |
| PC 36:0 | 661 | ND | ND | ND | ND | 479 | 539 |
| PE 36:4\|PE 16:0_20:4 | 662 | ND | ND | 15 | ND | ND | ND |
| LPC 16:0/0:0 | 663 | ND | ND | ND | ND | ND | ND |
| TG 46:2\|TG 14:0_16:1_16:1 | 664 | ND | ND | ND | ND | ND | ND |
| Cer 43:1;2O\|Cer 18:1;2O/25:0 | 665 | ND | ND | ND | ND | ND | 609 |
| 3'-O-Methylinosine | 666 | ND | ND | ND | ND | ND | ND |
| Kynurenic acid | 667 | ND | ND | ND | ND | ND | ND |
| 5.10_807.07 | 668 | ND | ND | ND | ND | ND | ND |
| TG O-56:6\|TG O-18:0_16:0_22:6 | 669 | ND | ND | ND | ND | ND | ND |
| 4-Guanidinobutyric acid | 670 | 182 | 20 | 27 | 105 | 266 | ND |
| gamma-Aminobutyric acid | 671 | ND | ND | 239 | ND | ND | ND |
| Pyridoxamine | 672 | 53 | 375 | 505 | 369 | ND | 627 |
| FA 34:1 | 673 | ND | ND | 138 | ND | 32 | ND |
| Corticosterone | 674 | ND | ND | ND | ND | ND | ND |
| DG 50:0 | 675 | ND | ND | ND | ND | ND | ND |
| Ribonic acid gamma-lactone | 676 | ND | ND | ND | ND | ND | ND |
| LPI 20:4 | 677 | ND | ND | 304 | ND | 1049 | ND |
| PE 38:4\|PE 18:0_20:4 | 678 | ND | ND | 468 | ND | ND | ND |
| Psicose | 679 | ND | ND | ND | ND | ND | ND |
| CAR 20:5 | 680 | ND | ND | ND | ND | 335 | ND |
| Thr-His | 681 | ND | ND | ND | ND | ND | 795 |
| LPC 22:5 a | 682 | ND | ND | ND | ND | ND | ND |
| S-Adenosyl-homocysteine | 683 | 10 | ND | 534 | ND | 722 | 648 |
| Tetrabutylammonium | 684 | ND | ND | ND | ND | ND | ND |
| DG 34:3 | 685 | ND | 176 | ND | 23 | 780 | 451 |
| Dexpanthenol | 686 | 47 | ND | 55 | ND | ND | ND |
| Cer 46:0;2O\|Cer 22:0;2O/24:0 | 687 | ND | ND | ND | ND | ND | ND |

| | | | | | | | |
|---|---|---|---|---|---|---|---|
| PI 36:4 \| PI 16:0_20:4_b | 688 | ND | ND | ND | ND | ND | ND |
| TG 58:11 \| TG 16:0_20:5_22:6 | 689 | ND | ND | ND | ND | ND | ND |
| PC O-42:6 | 690 | ND | ND | ND | ND | ND | ND |
| CMPF | 691 | ND | ND | ND | ND | ND | ND |
| Stearamide | 692 | ND | ND | 70 | ND | ND | ND |
| Glu-Thr | 693 | 461 | ND | ND | 127 | ND | 313 |
| FA 20:3;O | 694 | ND | ND | ND | ND | ND | ND |
| Decanoyl-L-carnitine | 695 | ND | ND | ND | ND | 810 | ND |
| TG 52:4;2O \| TG 16:0_18:1_18:3;2O | 696 | ND | ND | ND | ND | ND | ND |
| LPC O-16:1 | 697 | ND | ND | ND | ND | 1013 | ND |
| PC 38:2 | 698 | ND | ND | 193 | 302 | 491 | 564 |
| Uracil | 699 | ND | 350 | 627 | 30 | ND | 815 |
| SM 42:2;2O \| SM 18:2;2O/24:0 | 700 | ND | ND | ND | ND | ND | ND |
| O1_FA 20:5; (eicosapentaenoic acid)_b | 701 | ND | ND | ND | ND | ND | ND |
| LPC 19:1/0:0 | 702 | ND | ND | ND | ND | ND | ND |
| Ethylhexadecyldimethylammonium | 703 | ND | ND | ND | ND | ND | ND |
| N-Isovalerylglycine | 704 | ND | ND | ND | ND | ND | ND |
| Kynurenine | 705 | ND | ND | ND | ND | ND | ND |
| P-Anisic acid | 706 | ND | ND | ND | ND | ND | ND |
| Vitamin C | 707 | ND | ND | ND | ND | ND | ND |
| LPE 18:1 | 708 | ND | ND | 301 | 59 | 1024 | 399 |
| Sebacic acid | 709 | ND | ND | ND | ND | ND | ND |
| N-Formylmethionine | 710 | ND | ND | ND | ND | ND | ND |
| Norleucine | 711 | ND | ND | ND | ND | ND | ND |
| PC 36:2;O \| PC 18:0_18:2;O | 712 | ND | ND | ND | ND | ND | ND |
| N-(4-Hydroxy-3-methylphenyl)acetamide | 713 | ND | ND | ND | ND | ND | ND |
| LPE 18:2 | 714 | 35 | ND | 147 | 573 | 366 | 398 |
| DG 36:3 | 715 | 13 | 149 | ND | 322 | 785 | 339 |
| Thiazolidine-4-carboxylic acid | 716 | 94 | ND | ND | ND | ND | ND |
| Urocanic acid | 717 | ND | ND | 32 | ND | ND | ND |
| N-Isobutyrylglycine | 718 | ND | ND | ND | ND | ND | ND |
| N,N-Dimethyldodecylamine N-oxide | 719 | ND | 288 | 257 | ND | ND | ND |
| PC 35:2 \| PC 17:0_18:2 | 720 | ND | ND | ND | ND | ND | ND |
| FA 18:2 | 721 | 16 | ND | ND | 509 | ND | 926 |
| Cer 43:0;2O \| Cer 20:0;2O/23:0 | 722 | ND | ND | ND | ND | ND | ND |
| N1-(3-Aminopropyl)-N1-methylpropane-1,3-diamine | 723 | ND | ND | ND | ND | ND | ND |
| 4,5,7-Trihydroxyisoflavone | 724 | ND | ND | ND | ND | ND | ND |
| Indolelactic acid | 725 | ND | ND | ND | ND | ND | ND |
| PC O-36:2 | 726 | ND | ND | ND | ND | ND | ND |
| Phenylacetaldehyde | 727 | ND | ND | ND | ND | ND | ND |
| PC O-34:1 | 728 | ND | ND | ND | ND | 530 | ND |
| PC 38:6 \| PC 16:0_22:6 | 729 | ND | ND | ND | ND | ND | ND |
| FA 17:1 | 730 | ND | ND | 146 | ND | 829 | ND |

| | | | | | | | |
|---|---|---|---|---|---|---|---|
| N-Acetyl-D-glucosamine | 731 | 565 | ND | ND | 374 | ND | 423 |
| FA 20:4;2O | 732 | ND | ND | ND | ND | ND | ND |
| LPE 18:0 | 733 | 384 | ND | 328 | 104 | 1023 | 400 |
| 2-Mercaptomethylbenzimidazole | 734 | ND | ND | 124 | ND | ND | ND |
| SM 34:1;3O | 735 | ND | ND | ND | ND | ND | ND |
| Mandelic acid | 736 | ND | ND | ND | ND | ND | ND |
| Acetylcarnitine | 737 | 416 | ND | ND | 134 | 372 | ND |
| TG 60:7 \| TG 18:1_20:1_22:5 | 738 | ND | ND | 720 | ND | ND | ND |
| Pyridoxal phosphate | 739 | ND | ND | ND | ND | ND | ND |
| PC 38:4 \| PC 18:1_20:3 | 740 | ND | ND | ND | ND | ND | ND |
| PC 38:5 \| PC 16:0_22:5 | 741 | ND | ND | 451 | ND | ND | ND |
| PI 36:5 \| PI 16:0_20:5 | 742 | ND | ND | ND | ND | 673 | ND |
| Myristoyl-L-carnitine | 743 | 487 | ND | ND | ND | ND | ND |
| Thr-Gln | 744 | 567 | ND | ND | ND | 949 | 796 |
| CAR 20:4 | 745 | ND | ND | ND | ND | 648 | ND |
| FA 22:6 | 746 | ND | ND | ND | ND | ND | ND |
| gamma-Glutamylmethionine | 747 | 272 | ND | ND | ND | ND | ND |
| LPC 22:5 b | 748 | ND | ND | ND | ND | ND | ND |
| PE 34:2 \| PE 16:0_18:2 | 749 | ND | ND | 60 | ND | ND | ND |
| FA 17:0 | 750 | ND | ND | ND | 335 | ND | 4 |
| Creatinine | 751 | ND | 42 | 678 | ND | ND | 544 |
| PC 36:3 \| PC 18:1_18:2 | 752 | ND | ND | ND | ND | ND | ND |
| Proline | 753 | ND | 388 | 501 | ND | ND | ND |
| PC O-38:4 \| PC O-18:0_20:4 | 754 | ND | ND | ND | ND | ND | ND |
| LPE O-16:1 | 755 | ND | ND | 136 | ND | 1036 | ND |
| Indole-3-carboxylic acid | 756 | ND | ND | ND | ND | ND | ND |
| PC O-32:2 | 757 | ND | ND | ND | ND | ND | ND |
| FA 32:1 | 758 | ND | ND | ND | ND | ND | ND |
| Ribose | 759 | ND | ND | ND | ND | ND | ND |
| DMSO | 760 | 102 | 320 | ND | ND | ND | ND |
| LPC 20:2/0:0 | 761 | ND | ND | ND | ND | ND | ND |
| LPC 20:3/0:0 | 762 | ND | ND | ND | ND | ND | ND |
| PC O-38:6 \| PC O-16:0_22:6 | 763 | ND | ND | ND | ND | ND | ND |
| PC O-34:3 \| PC O-16:1_18:2 | 764 | ND | ND | ND | ND | ND | ND |
| O1_PC p-38:4; or PC o-38:5; B | 765 | ND | ND | ND | ND | ND | ND |
| Allantoin | 766 | ND | ND | ND | ND | ND | ND |
| Proline-hydroxyproline | 767 | ND | ND | ND | ND | ND | 632 |
| Salicylic acid | 768 | ND | ND | ND | ND | ND | ND |
| Palmitoylcarnitine | 769 | ND | ND | ND | ND | 705 | ND |
| N-alpha-methylhistamine | 770 | ND | ND | ND | ND | ND | ND |
| PC O-36:4 \| PC O-16:0_20:4 | 771 | ND | ND | ND | ND | ND | ND |
| TG 45:0 \| TG 14:0_15:0_16:0 | 772 | ND | ND | ND | ND | ND | 773 |
| CAR 14:0 | 773 | ND | ND | ND | ND | 599 | ND |

| | | | | | | | |
|---|---|---|---|---|---|---|---|
| 5.10_807.57 | 774 | ND | ND | ND | ND | ND | ND |
| N-Acetyl-valine | 775 | ND | ND | ND | ND | ND | ND |
| Pyridoxal | 776 | ND | ND | ND | 206 | 721 | 628 |
| N-(1H-Indol-3-ylacetyl)glycine | 777 | ND | ND | ND | ND | ND | 431 |
| Thr-Glu | 778 | ND | ND | ND | ND | ND | ND |
| CAR 14:1 | 779 | ND | ND | ND | ND | 17 | ND |
| Ferulic acid | 780 | ND | ND | ND | ND | ND | ND |
| SM 34:0;2O | 781 | ND | ND | ND | ND | ND | ND |
| PC O-37:5 | 782 | ND | ND | ND | ND | ND | ND |
| Octanoylcarnitine | 783 | ND | ND | ND | ND | 458 | ND |
| FA 18:1 | 784 | 504 | ND | ND | 292 | ND | 694 |
| TG O-55:2\|TG O-19:1_16:0_20:1 | 785 | ND | ND | ND | ND | ND | ND |
| FA 27:0 | 786 | ND | ND | ND | ND | ND | ND |
| PC O-39:10 | 787 | ND | ND | ND | ND | 193 | ND |
| Val-Gly | 788 | ND | ND | ND | ND | ND | ND |
| N-Acetylleucine | 789 | ND | ND | ND | ND | ND | ND |
| FA 36:0 | 790 | ND | ND | ND | ND | ND | ND |
| PC 41:6\|PC 19:0_22:6 | 791 | ND | ND | ND | ND | 514 | ND |
| Cer 40:2;2O\|Cer 18:2;2O/22:0 | 792 | ND | ND | ND | ND | ND | 656 |
| LPC 22:4/0:0 | 793 | ND | ND | ND | ND | ND | ND |
| TMAO | 794 | 9 | 33 | ND | ND | ND | ND |
| PC O-40:4 | 795 | ND | ND | ND | ND | ND | ND |
| Ribulofuranose | 796 | ND | ND | ND | ND | ND | ND |
| TG 58:9\|TG 18:1_18:2_22:6 | 797 | ND | ND | 703 | ND | ND | ND |
| PI 36:3\|PI 18:1_18:2 | 798 | ND | ND | 36 | ND | ND | ND |
| SM 38:1;2O | 799 | ND | ND | ND | ND | ND | ND |
| TG O-60:3\|TG O-22:1_18:1_20:1 | 800 | ND | ND | ND | ND | ND | ND |
| PC 38:5\|PC 18:1_20:4 | 801 | ND | ND | ND | ND | ND | ND |
| delta-Hydroxylysine | 802 | ND | ND | ND | ND | ND | ND |
| ST 24:1;O5 | 803 | ND | ND | ND | ND | ND | ND |
| TG 60:11\|TG 18:1_20:4_22:6 | 804 | ND | ND | ND | ND | ND | ND |
| Daidzein | 805 | ND | ND | ND | ND | ND | 941 |
| Dihydrouracil | 806 | ND | ND | ND | ND | ND | ND |
| FA 20:4 | 807 | ND | ND | ND | ND | ND | 907 |
| PC 39:6\|PC 17:0_22:6 | 808 | ND | ND | ND | ND | 11 | ND |
| FA 22:0 | 809 | 475 | ND | ND | 274 | ND | 902 |
| TG 53:0\|TG 16:0_18:0_19:0 | 810 | ND | ND | ND | ND | ND | ND |
| PI 36:3\|PI 16:0_20:3 | 811 | ND | ND | ND | ND | ND | ND |
| PC 38:6\|PC 18:2_20:4 | 812 | ND | ND | ND | ND | ND | ND |
| PE 40:7\|PE 18:1_22:6 | 813 | ND | ND | 480 | ND | 589 | ND |
| PC O-38:5\|PC O-18:1_20:4 | 814 | ND | ND | ND | ND | ND | ND |
| TG O-62:7\|TG O-22:1_18:0_22:6 | 815 | ND | ND | ND | ND | ND | ND |
| PE O-38:6\|PE O-16:0_22:6 | 816 | ND | ND | ND | ND | ND | ND |

| | | | | | | | |
|---|---|---|---|---|---|---|---|
| Lys-Leu | 817 | 19 | ND | ND | ND | 1058 | ND |
| Indole-3-acetamide | 818 | ND | ND | ND | ND | ND | ND |
| O3_PC 35:2 | 819 | ND | ND | ND | ND | ND | ND |
| Uridine | 820 | 141 | ND | ND | 269 | ND | ND |
| SM 40:2;2O | 821 | ND | ND | ND | ND | ND | ND |
| N-epsilon-Acetyllysine | 822 | 473 | ND | ND | ND | ND | 443 |
| Hippuric acid | 823 | ND | ND | ND | ND | ND | 364 |
| PE O-36:5 \| PE O-16:1_20:4 | 824 | ND | ND | 76 | ND | ND | ND |
| N-(4-Methoxyphenyl)acetamide | 825 | ND | ND | ND | ND | ND | ND |
| CAR 18:2 | 826 | ND | ND | ND | ND | 641 | ND |
| PE P-38:6 \| PE P-16:0_22:6 | 827 | ND | ND | ND | ND | 640 | ND |
| PE 36:1 \| PE 18:0_18:1 | 828 | ND | ND | 464 | ND | 564 | ND |
| FA 19:1 | 829 | ND | ND | 397 | ND | 878 | ND |
| PC O-34:4 | 830 | ND | ND | ND | ND | ND | ND |
| TG 60:9 \| TG 18:1_18:2_24:6 | 831 | ND | ND | ND | ND | ND | ND |
| FA 22:1 | 832 | 464 | ND | ND | 355 | ND | 900 |
| PI 36:3 | 833 | ND | ND | ND | ND | 670 | 619 |
| TG 58:8 \| TG 18:1_18:1_22:6 | 834 | ND | ND | 675 | ND | ND | ND |
| SM 40:2;2O \| SM 18:2;2O/22:0 | 835 | ND | ND | ND | ND | ND | ND |
| PC 40:5 \| PC 18:0_22:5 | 836 | ND | ND | ND | ND | ND | ND |
| CAR 16:2 | 837 | ND | ND | ND | ND | 423 | ND |
| PC 34:3 \| PC 16:1_18:2 | 838 | ND | ND | ND | ND | ND | ND |
| FA 22:5 | 839 | ND | ND | 353 | ND | 175 | ND |
| TG O-51:0 \| TG O-17:0_16:0_18:0 | 840 | ND | ND | ND | ND | ND | ND |
| FA 26:0 | 841 | ND | ND | ND | ND | ND | ND |
| PC 34:1 \| PC 16:0_18:1 | 842 | ND | ND | ND | ND | ND | ND |
| PI 36:1 \| PI 18:0_18:1_b | 843 | ND | ND | ND | ND | ND | ND |
| N-Acetylphenylalanine | 844 | ND | ND | ND | ND | ND | ND |
| Riboflavin | 845 | 22 | ND | ND | 66 | ND | 649 |
| TG 46:0 \| TG 14:0_16:0_16:0 | 846 | ND | ND | 129 | ND | ND | ND |
| ST 27:1;O;S | 847 | ND | ND | ND | ND | ND | ND |
| TG O-48:0 \| TG O-17:0_15:0_16:0 | 848 | ND | ND | 200 | ND | ND | ND |
| SM 34:2;2O \| SM 18:2;2O/16:0 | 849 | ND | ND | ND | ND | ND | ND |
| PC 36:3 \| PC 16:0_20:3 | 850 | ND | ND | ND | ND | ND | ND |
| PC 36:2 \| PC 18:1_18:1 | 851 | ND | ND | ND | ND | ND | ND |
| Salicylic alcohol | 852 | ND | ND | ND | ND | ND | ND |
| PC 32:1 \| PC 16:0_16:1 | 853 | ND | ND | ND | ND | ND | ND |
| m-Toluthioamide | 854 | ND | ND | ND | ND | ND | ND |
| Glycitein | 855 | ND | ND | ND | ND | ND | ND |
| PC 36:1 \| PC 18:0_18:1 | 856 | ND | ND | ND | ND | ND | ND |
| TG O-60:7 \| TG O-20:0_18:1_22:6 | 857 | ND | ND | ND | ND | ND | ND |
| LPE 20:4 | 858 | 159 | ND | 228 | 342 | 1030 | 397 |
| Iminodiacetic acid | 859 | ND | ND | ND | ND | ND | ND |

| | | | | | | | |
|---|---|---|---|---|---|---|---|
| TG 54:0 \| TG 18:0_18:0_18:0 | 860 | ND | ND | ND | ND | ND | ND |
| FA 9:0 | 861 | ND | ND | ND | 10 | ND | ND |
| PI 38:6 \| PI 16:0_22:6 | 862 | ND | ND | ND | ND | 681 | ND |
| Apigenin 7-glucuronide | 863 | ND | ND | ND | ND | ND | ND |
| N-Methylvaline | 864 | ND | ND | ND | ND | ND | ND |
| PC O-38:6 | 865 | ND | ND | ND | ND | 543 | ND |
| PI 40:6 | 866 | ND | ND | ND | ND | 686 | 615 |
| PC O-34:2 | 867 | ND | ND | ND | ND | 532 | ND |
| Daidzein 4'-sulfate | 868 | ND | ND | ND | ND | ND | ND |
| PC 34:2;O \| PC 16:0_18:2;O_b | 869 | ND | ND | ND | ND | ND | ND |
| FA 24:1 | 870 | 516 | ND | ND | 258 | ND | 892 |
| PI 40:6 \| PI 18:0_22:6 | 871 | ND | ND | 526 | ND | ND | ND |
| TG 58:0 \| TG 16:0_18:0_24:0 | 872 | ND | ND | ND | ND | ND | ND |
| TG 60:8 \| TG 18:1_20:1_22:6 | 873 | ND | ND | 722 | ND | 889 | ND |
| TG 46:0 \| TG 14:0_15:0_17:0 | 874 | ND | ND | ND | ND | ND | ND |
| PC 40:8 \| PC 18:2_22:6 | 875 | ND | ND | ND | ND | ND | ND |
| FA 20:1 | 876 | 191 | ND | ND | 359 | ND | 918 |
| DG 36:3 \| DG 18:1_18:2 | 877 | ND | ND | 267 | ND | ND | ND |
| PC 36:5 \| PC 16:0_20:5 | 878 | ND | ND | 425 | ND | ND | ND |
| Pro-Gln | 879 | 117 | ND | ND | 424 | 712 | 638 |
| Tuberonic acid glucoside | 880 | ND | ND | ND | ND | ND | ND |
| TG O-58:4 \| TG O-22:1_18:1_18:2 | 881 | ND | ND | ND | ND | ND | ND |
| TG 58:7 \| TG 18:0_18:1_22:6 | 882 | ND | ND | 672 | ND | ND | ND |
| SM 39:1;2O | 883 | ND | ND | ND | ND | ND | ND |
| PC 34:2 \| PC 16:0_18:2 | 884 | ND | ND | ND | ND | ND | ND |
| PE O-34:3 \| PE O-16:1_18:2 | 885 | ND | ND | 455 | ND | ND | ND |
| PC 37:2 | 886 | ND | ND | 426 | 534 | 37 | 530 |
| SM 42:3;2O \| SM 18:2;2O/24:1 | 887 | ND | ND | ND | ND | ND | ND |
| PI 36:4 | 888 | ND | ND | ND | ND | 671 | ND |
| CAR 16:1 | 889 | ND | ND | ND | ND | 605 | ND |
| PC 40:7 \| PC 18:1_22:6 | 890 | ND | ND | ND | ND | ND | ND |
| PC 35:1 \| PC 17:0_18:1 | 891 | ND | ND | ND | ND | ND | ND |
| FA 28:0 | 892 | ND | ND | ND | ND | ND | ND |
| O2_FA 20:2; (eicosadienoic acid) | 893 | ND | ND | ND | ND | ND | ND |
| PE O-38:5 \| PE O-18:1_20:4 | 894 | ND | ND | 495 | ND | 619 | ND |
| PC O-41:10 | 895 | ND | ND | ND | ND | ND | ND |
| PC 33:1 \| PC 16:0_17:1 | 896 | ND | ND | ND | ND | ND | ND |
| TG 54:5 \| TG 18:1_18:2_18:2 | 897 | ND | ND | ND | ND | ND | ND |
| 10.42_369.35 | 898 | ND | ND | ND | ND | ND | ND |
| TG O-54:0 \| TG O-20:0_16:0_18:0 | 899 | ND | ND | ND | ND | ND | ND |
| TG O-57:0 \| TG O-17:0_18:0_22:0 | 900 | ND | ND | ND | ND | ND | ND |
| PE O-36:2 \| PE O-18:0_18:2 | 901 | ND | ND | ND | ND | ND | ND |
| PE O-42:7 \| PE O-20:1_22:6 | 902 | ND | ND | 471 | ND | ND | ND |

| | | | | | | | |
|---|---|---|---|---|---|---|---|
| PE O-38:7 \| PE O-16:1_22:6 | 903 | ND | ND | 497 | ND | ND | ND |
| N-Carbobenzyloxyleucine | 904 | ND | ND | ND | ND | ND | ND |
| TG O-52:0 \| TG O-18:0_16:0_18:0 | 905 | ND | ND | ND | ND | ND | ND |
| PC O-36:5 | 906 | ND | ND | ND | ND | 536 | ND |
| PC O-36:4 | 907 | ND | ND | ND | ND | 321 | ND |
| PE O-40:7 \| PE O-18:1_22:6 | 908 | ND | ND | 499 | ND | 629 | ND |
| TG 59:11 \| TG 16:0_21:5_22:6 | 909 | ND | ND | ND | ND | ND | ND |
| DG 34:1 \| DG 16:0_18:1 | 910 | ND | ND | 262 | ND | ND | ND |
| methionine | 911 | ND | ND | ND | ND | 1037 | ND |
| TG O-58:3 \| TG O-22:1_18:0_18:2 | 912 | ND | ND | ND | ND | ND | ND |
| O2_FA 20:1; (eicosenoic acid) | 913 | ND | ND | ND | ND | ND | ND |
| TG O-50:3 \| TG O-16:0_16:0_18:3 | 914 | ND | ND | ND | ND | ND | ND |
| PC 36:4 \| PC 18:2_18:2 | 915 | ND | ND | ND | ND | ND | ND |
| TG O-57:13 \| TG O-21:5_18:4_18:4 | 916 | ND | ND | ND | ND | ND | ND |
| PE O-36:3 \| PE O-18:1_18:2 | 917 | ND | ND | ND | ND | ND | ND |
| Formononetine | 918 | ND | ND | ND | ND | ND | ND |
| CAR 18:1 | 919 | ND | ND | ND | ND | 626 | ND |
| TG 52:4;1O \| TG 16:0_18:2_18:2;1O | 920 | ND | ND | ND | ND | ND | ND |
| PE O-40:4 \| PE O-20:0_20:4 | 921 | ND | ND | ND | ND | ND | ND |
| Valine | 922 | 550 | ND | 710 | ND | ND | 809 |
| Methioninesulfoxide | 923 | ND | ND | 350 | ND | 103 | 436 |
| PC 37:2 \| PC 19:0_18:2 | 924 | ND | ND | ND | ND | ND | ND |
| TG O-52:1 \| TG O-18:0_16:0_18:1 | 925 | ND | ND | ND | ND | ND | ND |
| TG 60:9 \| TG 20:1_18:2_22:6 | 926 | ND | ND | 723 | ND | ND | ND |
| TG O-56:1 \| TG O-20:0_18:0_18:1 | 927 | ND | ND | ND | ND | ND | ND |
| TG O-52:2 \| TG O-18:0_16:0_18:2 | 928 | ND | ND | ND | ND | ND | ND |
| Arg-Leu | 929 | 30 | ND | ND | ND | 357 | 238 |
| TG 60:1 \| TG 16:0_26:0_18:1 | 930 | ND | ND | 687 | ND | ND | ND |
| TG 54:9 \| TG 18:2_18:3_18:4 | 931 | ND | ND | ND | ND | ND | ND |
| TG O-50:2 \| TG O-16:0_16:0_18:2 | 932 | ND | ND | ND | ND | ND | ND |
| TG 66:3 \| TG 16:0_32:1_18:2 | 933 | ND | ND | ND | ND | ND | ND |
| TG 49:2 \| TG 15:0_16:0_18:2 | 934 | ND | ND | 25 | ND | ND | ND |
| TG O-59:3 \| TG O-19:1_20:1_20:1 | 935 | ND | ND | ND | ND | ND | ND |
| PE O-40:5 \| PE O-18:0_22:5 | 936 | ND | ND | ND | ND | 624 | ND |
| TG 52:6 \| TG 16:1_18:2_18:3 | 937 | ND | ND | ND | ND | 1087 | ND |
| TG 52:7 \| TG 14:0_18:2_20:5 | 938 | ND | ND | ND | ND | ND | ND |
| TG 52:7 \| TG 18:2_18:2_16:3 | 939 | ND | ND | ND | ND | 151 | 765 |
| TG 54:7 \| TG 18:2_18:2_18:3 | 940 | ND | ND | ND | ND | ND | ND |
| TG 54:3 \| TG 18:1_18:1_18:1 | 941 | ND | ND | ND | ND | ND | ND |
| TG 54:2 \| TG 16:0_18:1_20:1 | 942 | ND | ND | ND | ND | ND | ND |
| TG 54:2 \| TG 18:0_18:1_18:1 | 943 | ND | ND | 618 | ND | ND | ND |
| TG 54:0 \| TG 16:0_18:0_20:0 | 944 | ND | ND | ND | ND | ND | ND |
| TG 53:4 \| TG 17:1_18:1_18:2 | 945 | ND | ND | 613 | ND | ND | ND |

| | | | | | | | |
|---|---|---|---|---|---|---|---|
| TG 53:5 \| TG 17:1_18:2_18:2 | 946 | ND | ND | 614 | ND | ND | ND |
| TG 53:6 \| TG 17:1_18:2_18:3 | 947 | ND | ND | 615 | ND | 826 | ND |
| TG 53:3 \| TG 17:0_18:1_18:2 | 948 | ND | ND | 612 | ND | ND | ND |
| TG 53:2 \| TG 16:0_18:1_19:1 | 949 | ND | ND | 643 | ND | ND | ND |
| TG 53:1 \| TG 16:0_19:0_18:1 | 950 | ND | ND | 642 | ND | ND | ND |
| TG 53:4 \| TG 17:0_18:2_18:2 | 951 | ND | ND | ND | ND | ND | ND |
| TG 52:8 \| TG 18:2_18:2_16:4 | 952 | ND | ND | 641 | ND | ND | ND |
| TG 50:5 \| TG 14:0_18:2_18:3 | 953 | ND | ND | 584 | ND | ND | ND |
| TG 50:5 \| TG 16:0_18:2_16:3 | 954 | ND | ND | ND | ND | ND | ND |
| TG 50:6 \| TG 16:0_18:2_16:4 | 955 | ND | ND | 585 | ND | ND | ND |
| Prostaglandin A3 | 956 | ND | ND | ND | ND | ND | ND |
| Ala-Asn | 957 | ND | ND | ND | ND | ND | 850 |
| Asn-Ala | 958 | ND | ND | ND | ND | ND | ND |
| BA 24:1;O3;T | 959 | ND | ND | ND | ND | ND | ND |
| TG 49:3 \| TG 15:0_16:1_18:2 | 960 | ND | ND | 89 | ND | ND | ND |
| TG 49:0 \| TG 15:0_16:0_18:0 | 961 | ND | ND | ND | ND | ND | ND |
| TG 48:1 \| TG 14:0_16:0_18:1 | 962 | ND | ND | 570 | ND | ND | ND |
| Liquiritigenin | 963 | ND | ND | ND | ND | ND | ND |
| Ala-Leu-Ala-Leu | 964 | ND | ND | ND | ND | ND | ND |
| O1_FA 22:6; (docosahexaenoic acid) | 965 | ND | ND | ND | ND | ND | ND |
| O1_PC 36:4; A | 966 | ND | ND | ND | ND | ND | ND |
| Tri-2-ethylhexyl trimellitate | 967 | ND | ND | ND | ND | ND | ND |
| o-Hydroxyhippuric acid | 968 | ND | ND | ND | ND | ND | ND |
| Tauroursodeoxycholic acid | 969 | ND | ND | ND | 296 | ND | 800 |
| Lys-Ile | 970 | ND | ND | ND | ND | ND | ND |
| SL 33:0;0O | 971 | ND | ND | ND | ND | ND | ND |
| TG 52:4 \| TG 16:0_16:0_20:4 | 972 | ND | ND | ND | ND | ND | ND |
| TG 52:1 \| TG 16:0_18:0_18:1 | 973 | ND | ND | 599 | ND | ND | ND |
| TG 51:5 \| TG 15:0_18:2_18:3 | 974 | ND | ND | 597 | ND | ND | ND |
| TG 52:6 \| TG 16:0_18:2_18:4 | 975 | ND | ND | ND | ND | ND | ND |
| TG 52:6 \| TG 16:0_16:1_20:5 | 976 | ND | ND | ND | ND | ND | ND |
| TG 52:6 \| TG 14:0_16:0_22:6 | 977 | ND | ND | ND | ND | ND | ND |
| TG 52:5 \| TG 16:0_16:0_20:5 | 978 | ND | ND | ND | ND | ND | ND |
| TG 52:4 \| TG 16:0_18:2_18:2 | 979 | ND | ND | ND | ND | ND | ND |
| TG 51:1 \| TG 16:0_17:0_18:1 | 980 | ND | ND | 625 | ND | ND | ND |
| TG 51:3 \| TG 16:0_17:1_18:2 | 981 | ND | ND | ND | ND | ND | ND |
| TG 51:0 \| TG 16:0_17:0_18:0 | 982 | ND | ND | 624 | ND | ND | 766 |
| TG 51:4 \| TG 15:0_18:2_18:2 | 983 | ND | ND | ND | ND | ND | ND |
| TG 50:1 \| TG 16:0_16:0_18:1 | 984 | ND | ND | 278 | ND | ND | ND |
| TG 56:1 \| TG 16:0_22:0_18:1 | 985 | ND | ND | ND | ND | ND | ND |
| PI 37:4 \| PI 17:0_20:4 | 986 | ND | ND | 201 | ND | 674 | ND |
| TG 56:0 \| TG 16:0_18:0_22:0 | 987 | ND | ND | ND | ND | ND | ND |
| PE O-38:4 \| PE O-18:0_20:4 | 988 | ND | ND | ND | ND | 616 | ND |

| | | | | | | | |
|---|---|---|---|---|---|---|---|
| Chrysogine | 989 | ND | ND | 676 | ND | ND | ND |
| TG 59:2 \| TG 16:0_25:0_18:2 | 990 | ND | ND | ND | ND | ND | ND |
| TG 58:6 \| TG 18:0_18:1_22:5 | 991 | ND | ND | 671 | ND | ND | ND |
| TG 58:6 \| TG 18:1_20:1_20:4 | 992 | ND | ND | ND | ND | ND | ND |
| TG 59:1 \| TG 16:0_25:0_18:1 | 993 | ND | ND | 704 | ND | 874 | ND |
| TG 59:4 \| TG 23:0_18:2_18:2 | 994 | ND | ND | 683 | ND | ND | ND |
| Oroxyloside | 995 | ND | ND | ND | ND | ND | 506 |
| TG 58:4 \| TG 22:0_18:2_18:2 | 996 | ND | ND | ND | ND | ND | ND |
| TG 58:3 \| TG 22:0_18:1_18:2 | 997 | ND | ND | 668 | ND | ND | ND |
| TG 58:2 \| TG 22:0_18:1_18:1 | 998 | ND | ND | ND | ND | ND | ND |
| TG 58:2 \| TG 16:0_18:1_24:1 | 999 | ND | ND | 667 | ND | ND | ND |
| TG 57:8 \| TG 18:1_18:2_21:5 | 1000 | ND | ND | 660 | ND | 860 | ND |
| TG 58:1 \| TG 16:0_24:0_18:1 | 1001 | ND | ND | 666 | ND | ND | ND |
| PE O-40:6 \| PE O-18:0_22:6 | 1002 | ND | ND | ND | ND | 627 | ND |
| TG 55:2 \| TG 16:0_21:0_18:2 | 1003 | ND | ND | ND | ND | ND | ND |
| Phe-Leu | 1004 | ND | ND | ND | ND | ND | ND |
| O1_FA 18:3; (linolenic acid) | 1005 | ND | ND | ND | ND | ND | ND |
| Pencolide | 1006 | ND | ND | ND | ND | ND | ND |
| TG 55:3 \| TG 19:0_18:1_18:2 | 1007 | ND | ND | 631 | ND | ND | ND |
| TG 56:2 \| TG 16:0_18:1_22:1 | 1008 | ND | ND | ND | ND | ND | ND |
| TG 56:4 \| TG 18:1_20:1_18:2 | 1009 | ND | ND | 71 | ND | ND | ND |
| TG 56:5 \| TG 16:0_18:1_22:4 | 1010 | ND | ND | ND | ND | ND | ND |
| TG 56:5 \| TG 18:0_18:1_20:4 | 1011 | ND | ND | ND | ND | ND | ND |
| TG 56:5 \| TG 20:1_18:2_18:2 | 1012 | ND | ND | ND | ND | ND | ND |
| Isobutyric acid | 1013 | ND | ND | ND | ND | ND | ND |
| L-Glutarylcarnitine | 1014 | ND | ND | ND | 259 | ND | 390 |
| TG 57:1 \| TG 16:0_23:0_18:1 | 1015 | ND | ND | 651 | ND | ND | ND |
| TG 57:2 \| TG 16:0_23:0_18:2 | 1016 | ND | ND | ND | ND | ND | ND |
| TG 57:3 \| TG 21:0_18:1_18:2 | 1017 | ND | ND | ND | ND | 856 | 791 |
| TG 57:4 \| TG 21:0_18:2_18:2 | 1018 | ND | ND | ND | ND | ND | ND |
| TG 57:8 \| TG 17:1_18:1_22:6 | 1019 | ND | ND | ND | ND | ND | ND |
| TG 55:6 \| TG 16:0_18:1_21:5 | 1020 | ND | ND | 634 | ND | 842 | ND |
| TG 55:5 \| TG 19:1_18:2_18:2 | 1021 | ND | ND | 633 | ND | 841 | 763 |
| TG O-53:6 \| TG O-15:0_19:3_19:3 | 1022 | ND | ND | ND | ND | ND | ND |
| TG O-50:1 \| TG O-16:0_16:0_18:1 | 1023 | ND | ND | ND | ND | ND | ND |
| TG 62:3 \| TG 26:0_18:1_18:2 | 1024 | ND | ND | 701 | ND | ND | ND |
| TG O-52:3 \| TG O-16:0_18:1_18:2 | 1025 | ND | ND | ND | ND | ND | ND |
| TG 62:4 \| TG 20:1_24:1_18:2 | 1026 | ND | ND | ND | ND | ND | ND |
| Carbanilide | 1027 | ND | ND | ND | ND | ND | ND |
| TG 60:5 \| TG 24:1_18:2_18:2 | 1028 | ND | ND | 755 | ND | ND | ND |
| Inosine | 1029 | ND | 309 | 309 | 416 | ND | 373 |
| TG 60:2 \| TG 18:0_24:0_18:2 | 1030 | ND | ND | ND | ND | ND | ND |
| TG 60:3 \| TG 24:0_18:1_18:2 | 1031 | ND | ND | 689 | ND | ND | ND |

| | | | | | | | |
|---|---|---|---|---|---|---|---|
| TG 60:4 \| TG 18:1_24:1_18:2 | 1032 | ND | ND | 690 | ND | ND | ND |
| TG 61:3 \| TG 25:0_18:1_18:2 | 1033 | ND | ND | 693 | ND | 892 | ND |
| PC 38:2 \| PC 20:0_18:2 | 1034 | ND | ND | ND | ND | ND | ND |
| TG O-54:1 \| TG O-20:0_16:0_18:1 | 1035 | ND | ND | ND | ND | ND | ND |
| PC 36:4;O \| PC 16:0_20:4;O | 1036 | ND | ND | ND | ND | ND | ND |
| Gln-Ser | 1037 | ND | ND | ND | ND | 927 | 135 |
| TG O-54:2 \| TG O-20:0_16:0_18:2 | 1038 | ND | ND | ND | ND | ND | ND |
| TG O-54:3 \| TG O-18:0_18:1_18:2 | 1039 | ND | ND | ND | ND | ND | ND |
| TG O-56:2 \| TG O-20:0_18:0_18:2 | 1040 | ND | ND | ND | ND | ND | ND |
| TG O-56:3 \| TG O-20:0_18:1_18:2 | 1041 | ND | ND | ND | ND | ND | ND |
| TG O-56:4 \| TG O-19:2_18:2_19:0 | 1042 | ND | ND | ND | ND | ND | ND |
| Gln-Ala | 1043 | 508 | ND | ND | ND | 159 | 317 |
| TG O-57:2 \| TG O-19:0_18:1_20:1 | 1044 | ND | ND | ND | ND | ND | ND |
| TG O-57:1 \| TG O-19:1_16:0_22:0 | 1045 | ND | ND | ND | ND | ND | ND |
| Glycocholic acid | 1046 | ND | ND | ND | ND | ND | ND |
| Glycyltyrosine | 1047 | ND | ND | ND | ND | ND | 324 |
| TG O-59:2 \| TG O-19:0_18:2_22:0 | 1048 | ND | ND | ND | ND | ND | ND |
| TG O-61:3 \| TG O-19:1_20:1_22:1 | 1049 | ND | ND | ND | ND | ND | ND |
| alpha.-Methylkynurenine | 1050 | ND | ND | ND | ND | ND | ND |
| Xanthine | 1051 | ND | 360 | 737 | 507 | ND | 808 |
| (1R,6S)-Bicyclo[4.1.0]heptane-7-carboxylic acid | ND | ND | 128 | ND | ND | ND | ND |
| (2.beta.,3.alpha.,5.alpha.,16.beta.,17.beta.)-2-(4-Morpholinyl)-16-(1-pyrrolidinyl)androstane-3,17-diol | ND | 120 | ND | ND | ND | ND | ND |
| (2E)-2-Cyano-3-(4-morpholinyl)-2-propenamide | ND | ND | ND | ND | 47 | ND | ND |
| (2E,4E)-Hexa-2,4-dienoic acid | ND | 135 | ND | ND | ND | ND | ND |
| (2S)-pyrrolidine-2-carboxylic acid | ND | ND | 62 | ND | ND | ND | ND |
| (3-Aminophenyl)acetonitrile | ND | ND | ND | 46 | ND | ND | ND |
| (3-Carboxypropyl)trimethylammonium_a | ND | ND | ND | ND | ND | 166 | ND |
| (3-Nitrophenyl)methanamine | ND | ND | 94 | 56 | ND | ND | 39 |
| (5S)-1,2,3,4,5,6,7,8,9,10-Decahydro-5,9-methanobenzo[8]annulen-11-one | ND | ND | ND | ND | ND | ND | 20 |
| (E)-.alpha.-Ionone | ND | ND | 91 | 45 | ND | ND | ND |
| (R)-Pantetheine | ND | ND | ND | ND | 38 | ND | ND |
| (Z)-9,12,13-Trihydroxyoctadec-15-enoic acid | ND | ND | ND | 65 | ND | ND | ND |
| .beta.-Methylphenethylamine | ND | ND | 137 | ND | ND | ND | ND |
| .gamma.-Aminobutyric acid | ND | ND | 106 | ND | ND | ND | ND |
| 1,2,3,3-Tetramethyl-norbornan-2-ol | ND | ND | ND | 137 | ND | ND | ND |
| 1,2,3,3-Tetramethylbicyclo[2.2.1]heptan-2-ol | ND | ND | 168 | ND | ND | ND | ND |
| 1,2,4-benzenetriol | ND | ND | ND | ND | 44 | 194 | ND |
| 1,2-Bis(O-decanoyl)-sn-glyceryl-3-phosphorylcholine | ND | ND | ND | ND | ND | ND | 88 |
| 1,2-Cyclohexanedimethanol, 1,2-dimethanesulfonate, (1R,2R)- | ND | ND | 141 | ND | ND | ND | ND |
| 1,2-Di-(9Z,12Z,15Z-octadecatrienoyl)-sn-glycero-3-phosphocholine | ND | ND | ND | ND | ND | 121 | ND |
| 1,2-Diamino-2-methylpropane | ND | ND | 102 | 81 | ND | 200 | ND |
| 1,2-Diarachidonoyl-sn-glycero-3-phosphocholine | ND | ND | ND | ND | ND | 227 | 81 |
| 1,2-Dilauroyl-sn-glycero-3-phosphatidylcholine | ND | ND | 208 | ND | ND | ND | ND |

| | | | | | | | |
|---|---|---|---|---|---|---|---|
| 1,2-Dilinoleoylglycerol | ND | ND | ND | ND | ND | 134 | ND |
| 1,2-anhydro-myo-inositol | ND | 158 | ND | ND | ND | ND | ND |
| 1,3,5-Benzenetriol | ND | ND | ND | ND | ND | 167 | ND |
| 1,4,7,10,13,16-Hexaoxacyclooctadecane | ND | ND | ND | 105 | ND | ND | ND |
| 1,4,7,10-Tetraazacyclododecane | ND | ND | ND | 101 | ND | ND | ND |
| 1,4-Cyclohexanedione | ND | ND | 163 | 54 | ND | ND | ND |
| 1,5-Pentanediamine | ND | ND | 310 | 727 | ND | ND | ND |
| 1,5-Pentanediamine_a | ND | ND | ND | ND | ND | 219 | ND |
| 1,5-anhydroglucitol | ND | 85 | ND | ND | 22 | 66 | 31 |
| 1-(1Z-Hexadecenyl)-sn-glycero-3-phosphocholine | ND | ND | ND | ND | ND | ND | 46 |
| 1-(2,5-Dimethylphenoxy)-3-(4-morpholinyl)-2-propanol | ND | 66 | ND | ND | ND | ND | ND |
| 1-(2-Aminopropoxy)-2-methoxyethane | ND | ND | ND | 84 | ND | ND | ND |
| 1-(2-Hydroxyethyl)pyrazole | ND | ND | ND | 10 | ND | 62 | ND |
| 1-(4-Piperidinyl)-2-pyrrolidinone | ND | ND | ND | 93 | ND | ND | ND |
| 1-(Methylsulfonyl)-1,4-diazepane | ND | ND | ND | 57 | ND | ND | ND |
| 1-Acetamidocyclopentane-1-carboxylic acid | ND | ND | 179 | ND | ND | ND | ND |
| 1-Cyclobutyl-4-piperidinamine | ND | ND | ND | 33 | ND | ND | ND |
| 1-Cycloheptyl-4-piperidinamine | ND | ND | 196 | 78 | ND | ND | ND |
| 1-Docosahexaenoyl-2-stearoyl-sn-glycero-3-phosphocholine | ND | ND | ND | ND | ND | 343 | ND |
| 1-Ethyl-3-piperidinamine | ND | ND | 101 | 92 | ND | ND | ND |
| 1-Ethyl-N4,N5-dimethyl-1H-imidazole-4,5-dicarboxamide | ND | ND | ND | ND | ND | 187 | ND |
| 1-Hexadecyl-2-(5Z,8Z,11Z,14Z-eicosatetraenoyl)-sn-glycero-3-phosphocholine | ND | ND | ND | ND | ND | ND | 30 |
| 1-Hexadecyl-2-(9Z-octadecenoyl)-sn-glycero-3-phosphocholine | ND | ND | ND | ND | ND | ND | 18 |
| 1-Methyl-1H-indole-3-carboxamide | ND | ND | ND | 108 | ND | ND | ND |
| 1-Methyl-4-piperidinecarboxylic acid | ND | ND | ND | ND | ND | ND | 55 |
| 1-Methylguanosine | ND | ND | ND | ND | 52 | ND | 60 |
| 1-Methylhistamine | ND | ND | ND | ND | ND | 111 | ND |
| 1-Naphthoic acid | ND | ND | 88 | 153 | ND | ND | ND |
| 1-Nitro-3-(4-nitrophenoxy)benzene | ND | ND | ND | ND | ND | ND | 62 |
| 1-Oleoyl-2-myristoyl-sn-glycero-3-phosphocholine | ND | ND | ND | ND | ND | 106 | 10 |
| 1-Palmitoyl-2-arachidonoyl-sn-glycero-3-phosphoserine | ND | ND | ND | ND | ND | ND | 82 |
| 1-Palmitoyl-2-docosahexaenoyl-sn-glycero-3-phosphocholine_b | ND | ND | ND | ND | ND | 146 | ND |
| 1-Palmitoyl-2-glutaryl phosphatidylcholine | ND | ND | ND | ND | ND | ND | 67 |
| 1-Stearoyl-2-arachidonoyl-sn-glycero-3-phospho-(1'-myo-inositol) | ND | ND | ND | ND | ND | 101 | ND |
| 1-Stearoyl-2-arachidonoyl-sn-glycero-3-phosphoserine | ND | ND | ND | ND | ND | ND | 69 |
| 1-Stearoyl-2-docosahexaenoyl-sn-glycerol | ND | 103 | ND | ND | ND | 112 | 75 |
| 1-Stearoyl-2-linoleoyl-sn-glycero-3-phospho-L-serine | ND | ND | ND | ND | ND | ND | 1 |
| 1-Stearoyl-2-myristoyl-sn-glycero-3-phosphocholine | ND | ND | ND | ND | ND | ND | 22 |
| 1-Stearoyl-2-oleoyl-sn-glycero-3-phosphoethanolamine | ND | ND | ND | ND | ND | 230 | ND |
| 1-Stearoylglycerol | ND | ND | ND | ND | 56 | ND | ND |
| 1-kestose | ND | ND | ND | ND | ND | 153 | 25 |
| 1-methylgalactose | ND | ND | ND | ND | ND | 143 | ND |
| 1-monopalmitin | ND | ND | ND | ND | ND | 189 | ND |

| | | | | | | | |
|---|---|---|---|---|---|---|---|
| 1-monostearin | ND | ND | ND | ND | ND | 202 | 28 |
| 1.24_193.12 | ND | 67 | ND | ND | ND | ND | ND |
| 10-Aminodecanoic acid | ND | ND | 41 | 21 | ND | ND | ND |
| 10-Undecynoic acid | ND | ND | 219 | 244 | ND | ND | ND |
| 11,15-Dioxo-9S-hydroxy-5Z-prostenoic acid | ND | ND | ND | ND | 85 | ND | ND |
| 13-cis-Retinol | ND | ND | ND | ND | 32 | ND | ND |
| 15.alpha.-Hydroxyculmorin | ND | ND | 170 | ND | ND | ND | ND |
| 1H-1,2,3-Benzotriazol-1-ylmethanol | ND | ND | ND | 96 | ND | ND | ND |
| 1H-Indole-3-ethanamine, 5-methoxy-N,N-di-2-propen-1-yl- | ND | 31 | ND | ND | ND | ND | ND |
| 2'-Deoxyguanosine | ND | 74 | ND | ND | ND | ND | 112 |
| 2'-Deoxyguanosine 3'-monophosphate | ND | ND | ND | ND | ND | ND | 73 |
| 2'-O-Methylinosine | ND | ND | ND | ND | ND | ND | 172 |
| 2,3,4,5-Tetrahydro-1,4-benzothiazepine | ND | ND | ND | ND | ND | ND | 253 |
| 2,3-dihydroxybutanoic acid | ND | ND | ND | ND | ND | ND | 95 |
| 2,4-Dimethylnicotinic acid | ND | ND | ND | ND | ND | 304 | 83 |
| 2,4-Dimethylthiazole-5-carboxylic acid | ND | ND | 211 | 738 | ND | ND | ND |
| 2,4-hexadienedioic acid | ND | 99 | ND | ND | ND | ND | 104 |
| 2,5-dihydroxypyrazine | ND | 246 | ND | ND | 69 | 78 | 86 |
| 2,6-Di-tert-butyl-4-(4-morpholinylmethyl)phenol | ND | ND | 158 | ND | ND | ND | ND |
| 2-(1H-Benzimidazol-2-ylthio)-N,N-dimethylethanamine | ND | 231 | ND | ND | ND | ND | ND |
| 2-(2-Aminoethanesulfonyl)propane | ND | ND | ND | 141 | ND | ND | ND |
| 2-(2-Anilino-2-oxoethoxy)benzoic acid | ND | ND | ND | ND | ND | ND | 77 |
| 2-(2-Hydroxyphenyl)-1,3-thiazolidine-4-carboxylic acid | ND | ND | ND | 205 | ND | ND | ND |
| 2-(2-Quinolyl)ethanol | ND | ND | 160 | ND | ND | ND | ND |
| 2-(3,8-Dihydroxy-8-(hydroxymethyl)-3-methyl-2-oxodecahydroazulen-5-yl)propan-2-yl hexopyranoside | ND | ND | ND | ND | ND | ND | 84 |
| 2-(5-Oxovaleryl)phosphatidylcholine | ND | ND | ND | ND | ND | ND | 679 |
| 2-Amino-2-deoxy-D-glucopyranose | ND | ND | ND | ND | ND | ND | 59 |
| 2-Amino-3-(2,4-diketo-6,7-dihydro-5H-cyclopenta[d]pyrimidin-1-yl)propionic acid | ND | ND | ND | ND | ND | 165 | ND |
| 2-Amino-5-phenylpentanoic acid | ND | ND | 224 | 260 | ND | ND | ND |
| 2-Amino-N,N-dimethylacetamide | ND | 178 | ND | ND | ND | ND | ND |
| 2-Arachidonoyl-1-palmitoyl-sn-glycero-3-phosphoethanolamine | ND | ND | ND | ND | ND | ND | 16 |
| 2-Arachidonoylglycerol | ND | ND | ND | ND | 46 | ND | 65 |
| 2-Carbamoylpyridine-3-carboxylic acid | ND | 370 | ND | ND | ND | ND | 883 |
| 2-Docosahexaenoyl-1-palmitoyl-sn-glycero-3-phosphoethanolamine | ND | ND | ND | ND | ND | ND | 140 |
| 2-Docosahexaenoyl-1-stearoyl-sn-glycero-3-phosphoethanolamine | ND | ND | ND | ND | ND | 209 | 49 |
| 2-Docosahexaenoyl-1-stearoyl-sn-glycero-3-phosphoserine | ND | ND | ND | ND | ND | 174 | ND |
| 2-Ethoxy-2-oxoethyl 2-hydroxybenzoate | ND | ND | ND | 94 | ND | ND | ND |
| 2-Hydroxy-4-(octyloxyl)benzophenone | ND | ND | ND | 85 | ND | ND | ND |
| 2-LPC 18:2 | ND | ND | ND | ND | 51 | ND | ND |
| 2-Methylbutyryl-carnitine | ND | ND | ND | ND | ND | 199 | ND |
| 2-Methylnicotinic acid | ND | 84 | ND | ND | ND | ND | ND |
| 2-Nitrobenzylamine | ND | ND | ND | ND | 113 | ND | ND |
| 2-Oleoyl-1-stearoyl-sn-glycero-3-phosphoserine | ND | ND | ND | ND | ND | 232 | 100 |

| | | | | | | | |
|---|---|---|---|---|---|---|---|
| 2-Oxo-2,3-dihydro-3-pyridinecarboxylic acid | ND | ND | ND | ND | 61 | ND | ND |
| 2-Palmitoyl-rac-glycerol | ND | ND | ND | ND | ND | ND | 96 |
| 2-Palmitoylglycerol | ND | ND | ND | ND | 81 | ND | ND |
| 2-Pyridone | ND | 62 | ND | ND | ND | ND | ND |
| 2-[(Dimethylamino)methyl]cyclohexanone | ND | ND | ND | ND | ND | 79 | ND |
| 2-[1-(Hydroxymethyl)propyl]-1H-isoindole-1,3(2H)-dione | ND | ND | ND | ND | ND | ND | 826 |
| 2-deoxypentitol | ND | 204 | ND | ND | 65 | ND | 111 |
| 2-deoxytetronic acid | ND | ND | ND | ND | ND | ND | 58 |
| 2-ethylcaproic acid | ND | ND | ND | ND | ND | ND | 41 |
| 2-hydroxybutanoic acid | ND | ND | ND | ND | 82 | ND | 90 |
| 2-hydroxyglutaric acid | ND | 116 | ND | ND | 37 | 85 | 99 |
| 2-hydroxypyrazinyl-2-propenoic acid ethyl ester | ND | ND | ND | ND | ND | 170 | ND |
| 2-hydroxyvaleric acid | ND | ND | ND | ND | ND | ND | 34 |
| 2-ketoglucose dimethylacetal | ND | 181 | ND | ND | 48 | 126 | 101 |
| 2-monopalmitin | ND | ND | ND | ND | ND | 89 | ND |
| 2-phosphoglyceric acid | ND | ND | ND | ND | ND | ND | 79 |
| 2-picolinic acid | ND | ND | ND | ND | ND | 323 | 89 |
| 3,3-Dimethylpyrrolidin-2-one | ND | ND | 343 | ND | ND | ND | ND |
| 3,4,5-Trimethoxydihydrocinnamic acid | ND | ND | ND | 214 | ND | ND | ND |
| 3,4-Difluorobenzenecarboximidamide | ND | ND | 348 | ND | ND | ND | ND |
| 3,4-Dihydroxyphenylalanine | ND | 291 | ND | ND | ND | ND | ND |
| 3,6-anhydro-D-galactose | ND | ND | ND | ND | 54 | 162 | 38 |
| 3,6-anhydro-D-glucose | ND | ND | ND | ND | 58 | ND | ND |
| 3-(2,4-Dimethoxyanilino)-3-oxopropanoic acid | ND | ND | ND | 142 | ND | ND | ND |
| 3-(3-Methoxybenzyl)piperidine | ND | ND | ND | ND | ND | ND | 121 |
| 3-(5-Chloro-1,3-benzothiazol-2-yl)propanoic acid | ND | ND | ND | ND | ND | ND | 144 |
| 3-(6-Amino-9H-purin-9-yl)propanoic acid | ND | ND | 115 | 75 | ND | ND | ND |
| 3-(Carboxymethyl)-1-(beta-D-glucopyranosyl)-1H-indole | ND | ND | ND | 132 | ND | ND | ND |
| 3-Amino-4-methylbenzenesulfonic acid | ND | ND | ND | ND | ND | ND | 167 |
| 3-Aminoadipic acid | ND | ND | ND | ND | ND | 152 | 245 |
| 3-Aminonaphthalene-2-carboxylic acid | ND | ND | ND | ND | ND | ND | 190 |
| 3-Aminopiperidine-2,6-dione | ND | ND | ND | ND | ND | 100 | 128 |
| 3-Aminotyrosine | ND | 207 | ND | 99 | ND | ND | 78 |
| 3-Chloro-2-[(4-methylphenyl)sulfanyl]aniline | ND | ND | 364 | 507 | ND | ND | ND |
| 3-Cyclohexyl-1,1-dimethylurea | ND | ND | 265 | ND | ND | 218 | ND |
| 3-Ethyl-2-piperazinone | ND | ND | ND | ND | ND | ND | 207 |
| 3-Hydroxyhexadecanoylcarnitine | ND | ND | ND | ND | ND | 212 | ND |
| 3-Hydroxyisovaleroylcarnitine | ND | 125 | ND | 203 | ND | 239 | ND |
| 3-Hydroxyoleylcarnitine | ND | ND | ND | ND | ND | 267 | 214 |
| 3-Hydroxypicolinic acid | ND | ND | ND | ND | ND | ND | 779 |
| 3-Hydroxypyridine | ND | ND | ND | 73 | ND | ND | ND |
| 3-Indoleacrylic acid | ND | ND | ND | ND | ND | ND | 177 |
| 3-Methyl-L-histidine | ND | ND | 198 | 191 | ND | ND | ND |

| | | | | | | | |
|---|---|---|---|---|---|---|---|
| 3-aminoisobutyric acid | ND | 152 | ND | ND | 35 | ND | 130 |
| 3-hydroxy-3-methylglutaric acid | ND | 220 | ND | ND | ND | ND | ND |
| 3-hydroxybutyric acid | ND | ND | ND | ND | 99 | 220 | 109 |
| 3-hydroxypropionic acid | ND | 171 | ND | ND | 94 | ND | ND |
| 3-phosphoglycerate | ND | ND | ND | ND | ND | ND | 161 |
| 3.alpha.-Galactobiose | ND | ND | ND | ND | ND | ND | 882 |
| 4',5-Dihydroxy-3',6,7-trimethoxyflavone | ND | ND | ND | ND | ND | 397 | ND |
| 4,8-Dimethylquinolin-2-ol | ND | ND | 368 | ND | ND | ND | ND |
| 4-(1H-Pyrazol-1-yl)butanoic acid | ND | ND | ND | ND | ND | ND | 668 |
| 4-(2,6,6-Trimethyl-3-oxocyclohex-1-en-1-yl)butan-2-yl .beta.-D-glucopyranoside | ND | ND | 365 | ND | ND | ND | ND |
| 4-(Allylamino)benzoic acid | ND | ND | ND | ND | ND | ND | 914 |
| 4-Acetyl-L-phenylalanine | ND | ND | ND | ND | ND | ND | 913 |
| 4-Amino-1-butanol | ND | ND | 99 | 144 | ND | ND | ND |
| 4-Aminobenzoic acid | ND | 46 | 252 | 508 | ND | ND | ND |
| 4-Aminomethylcyclohexanecarboxylic acid | ND | ND | 253 | 509 | ND | 380 | ND |
| 4-Aminomethyltetrahydropyran | ND | ND | ND | ND | ND | 235 | ND |
| 4-Cyano-D-phenylalanine | ND | ND | 283 | ND | ND | ND | ND |
| 4-Hydroxyamphetamine | ND | ND | ND | 219 | ND | ND | ND |
| 4-Hydroxymandelonitrile | ND | 82 | ND | ND | ND | ND | ND |
| 4-Imidazoleacrylic acid | ND | ND | 13 | ND | ND | 215 | ND |
| 4-Methyl-1H-pyrazole | ND | ND | 255 | 342 | ND | ND | ND |
| 4-Methyl-1H-pyrazole_4-Methyl-1H-pyrazole | ND | ND | ND | ND | ND | 127 | ND |
| 4-Methyl-5-thiazoleethanol? | ND | ND | ND | ND | ND | ND | 44 |
| 4-Methylpiperidine-1-carboximidamide | ND | 18 | ND | ND | ND | ND | ND |
| 4-Phenyl-1H-pyrazol-3-ylamine | ND | ND | ND | 253 | ND | ND | ND |
| 4-Piperidinecarboxamide | ND | ND | ND | 116 | ND | ND | ND |
| 4-Trifluoromethylpyrimidine-2-carbaldehyde | ND | ND | ND | 447 | ND | ND | ND |
| 4-aminobutyric acid | ND | 112 | ND | ND | 93 | ND | ND |
| 4-chlorophenylisobutylamine | ND | ND | ND | 448 | ND | ND | ND |
| 4-hydroxybutyric acid | ND | ND | ND | ND | ND | 342 | 156 |
| 5'-Methylthioadenosine | ND | 192 | ND | ND | 138 | ND | 912 |
| 5'-deoxy-5'-methylthioadenosine | ND | ND | ND | ND | ND | 228 | ND |
| 5,6-Dihydro-4H-pyrrolo[3,4-d]thiazole | ND | ND | ND | ND | ND | ND | 911 |
| 5-(6-Methyl-7-oxooctyl)furan-2(5H)-one | ND | ND | ND | 224 | ND | ND | ND |
| 5-(Galactosylhydroxy)-L-lysine | ND | ND | ND | ND | ND | ND | 910 |
| 5-(Hydroxymethyl)-2-furaldehyde | ND | ND | ND | ND | ND | ND | 122 |
| 5-Amino-1-.beta.-D-ribofuranosyl-1H-imidazole-4-carboxamide | ND | 258 | ND | ND | ND | ND | 287 |
| 5-Aminoimidazole-4-carboxamide | ND | ND | ND | ND | 201 | ND | ND |
| 5-Aminoorotic acid | ND | ND | ND | 263 | ND | ND | ND |
| 5-Chloro-1H-indole-3-carbaldehyde | ND | ND | 79 | ND | ND | ND | ND |
| 5-Hydroxymethylcytosine | ND | 121 | ND | ND | ND | ND | ND |
| 5-Methoxy-2-nitrophenylamine | ND | ND | ND | ND | 20 | 248 | 288 |
| 5-Methylisoxazol-3-amine | ND | ND | 223 | ND | ND | ND | ND |

| | | | | | | | |
|---|---|---|---|---|---|---|---|
| 5-Methyluridine | ND | ND | ND | ND | ND | ND | 9 |
| 5-aminovaleric acid | ND | ND | ND | ND | 186 | ND | 221 |
| 6-Biopterin | ND | ND | ND | ND | 117 | ND | 145 |
| 6-Methylnicotinic acid | ND | ND | ND | ND | ND | ND | 200 |
| 6-Phenylthiomorpholin-3-one | ND | ND | 377 | ND | ND | ND | ND |
| 6-deoxyglucitol | ND | ND | ND | ND | ND | ND | 153 |
| 7-Keto-8-aminopelargonic acid | ND | 44 | 70 | ND | ND | ND | ND |
| 7-Methylguanine | ND | 26 | ND | ND | ND | ND | ND |
| 8-Methylcaffeine | ND | ND | 113 | 19 | ND | ND | ND |
| 8-Nitro-7-methoxyisoquinoline | ND | ND | ND | ND | ND | ND | 258 |
| 8-Oxo-2-deoxyadenosine | ND | 374 | 61 | ND | ND | ND | ND |
| 8-Oxononanoic acid | ND | ND | 299 | 248 | ND | ND | ND |
| AC 02:0 | ND | ND | ND | 449 | ND | ND | ND |
| AC 03:0 | ND | ND | ND | 110 | ND | ND | ND |
| AC 06:0 | ND | ND | ND | 118 | ND | ND | ND |
| AC 12:0 | ND | 235 | ND | ND | ND | ND | ND |
| AC 14:0 | ND | ND | ND | ND | ND | ND | 175 |
| AC 14:1 | ND | 20 | ND | ND | ND | ND | ND |
| AC 16:0 | ND | 156 | ND | 172 | ND | ND | 176 |
| AC 16:1 | ND | ND | ND | 266 | ND | ND | 127 |
| AC 18:0 | ND | 217 | ND | 450 | ND | ND | 159 |
| AC 18:1 | ND | 157 | ND | 270 | ND | ND | 97 |
| AC 18:2 | ND | 421 | ND | 148 | ND | ND | 23 |
| AC 2-Methylbutyryl- | ND | ND | ND | 332 | ND | ND | ND |
| AC 20:0 | ND | ND | ND | ND | ND | ND | 174 |
| AC 20:1 | ND | ND | ND | ND | ND | ND | 70 |
| AC 20:2 | ND | ND | ND | ND | ND | ND | 236 |
| AC 20:3 | ND | ND | ND | ND | ND | ND | 134 |
| AC 20:4 | ND | ND | ND | ND | ND | ND | 131 |
| AC 21:0 | ND | ND | ND | ND | ND | ND | 119 |
| AC 21:3 | ND | ND | ND | ND | ND | ND | 166 |
| AC 22:0 | ND | ND | ND | ND | ND | ND | 209 |
| AC 22:1 | ND | ND | ND | ND | ND | ND | 26 |
| AC 22:2 | ND | ND | ND | ND | ND | ND | 103 |
| AC 22:6 | ND | ND | ND | ND | ND | ND | 186 |
| AC 24:0 | ND | ND | ND | ND | ND | ND | 187 |
| AC 24:1 | ND | ND | ND | ND | ND | ND | 40 |
| AC 24:2 | ND | ND | ND | ND | ND | ND | 192 |
| AC 3-Hydroxybutyryl | ND | ND | ND | 333 | ND | ND | ND |
| AC Malonyl- | ND | ND | ND | 334 | ND | ND | ND |
| ALDOPC | ND | ND | ND | ND | 87 | ND | ND |
| AMP | ND | ND | ND | ND | 132 | ND | ND |
| Ac-Asp-Glu | ND | ND | ND | ND | ND | 242 | ND |

| | | | | | | | |
|---|---|---|---|---|---|---|---|
| Acetyl coenzyme A | ND | 40 | ND | ND | ND | ND | ND |
| Acetyl-L-carnitine | ND | ND | ND | ND | ND | ND | 196 |
| Acylcarnitine 18:3 | ND | ND | ND | ND | 119 | ND | ND |
| Adenosine 5'-diphosphoribose | ND | ND | ND | ND | 418 | ND | 263 |
| Adenosine 5'-monophosphate | ND | ND | ND | ND | ND | ND | 867 |
| Adenosine-3-monophosphate | ND | ND | ND | ND | ND | ND | 866 |
| Adenylosuccinic acid | ND | 190 | ND | ND | 84 | ND | 865 |
| Adipoyl-L-carnitine | ND | ND | ND | ND | 71 | ND | 197 |
| Adrenochrome | ND | 323 | ND | ND | ND | ND | ND |
| Ala-Glu.1 | ND | 367 | ND | ND | ND | ND | ND |
| Ala-Ile-Lys | ND | ND | ND | ND | ND | ND | 848 |
| Ala-Leu | ND | ND | ND | ND | ND | 102 | ND |
| Ala-Lys | ND | ND | ND | ND | 112 | 383 | 847 |
| Ala-Phe | ND | ND | ND | ND | ND | 43 | ND |
| Ala-Pro | ND | ND | ND | ND | ND | ND | 846 |
| Ala-Ser | ND | ND | ND | ND | ND | 427 | ND |
| Allopurinol riboside | ND | ND | ND | ND | ND | 157 | 844 |
| Amantadine | ND | ND | ND | ND | ND | ND | 164 |
| Amobarbital | ND | 119 | ND | ND | ND | ND | ND |
| Androstane-3,17-diol | ND | 228 | ND | ND | ND | ND | ND |
| Androsterone | ND | 360 | ND | ND | ND | ND | ND |
| Aniline | ND | ND | 181 | 230 | ND | ND | ND |
| Arachidonoylcarnitine | ND | ND | ND | ND | ND | 268 | ND |
| Arachidonoylglycine | ND | 456 | ND | ND | ND | ND | ND |
| Arg-Asn | ND | ND | ND | ND | ND | ND | 843 |
| Arg-Asp | ND | 506 | ND | ND | ND | ND | 842 |
| Arg-Gln | ND | ND | ND | ND | ND | ND | 841 |
| Arg-Glu | ND | 404 | ND | ND | 79 | 263 | ND |
| Arg-Gly | ND | 410 | ND | ND | ND | 324 | 840 |
| Arg-Met | ND | 33 | ND | ND | ND | ND | ND |
| Arg-Phe | ND | 278 | ND | ND | ND | ND | ND |
| Arg-Pro | ND | ND | ND | ND | ND | ND | 168 |
| Arg-Ser | ND | 483 | ND | ND | ND | ND | ND |
| Arg-Tyr | ND | 143 | ND | ND | ND | ND | ND |
| Arg-Val | ND | 480 | ND | ND | ND | 421 | 213 |
| Argininosuccinic acid | ND | 521 | ND | 339 | 150 | 70 | 834 |
| Asn-Arg | ND | ND | ND | ND | 118 | 391 | 833 |
| Asn-His | ND | ND | ND | ND | ND | 414 | 832 |
| Asn-Lys | ND | ND | ND | ND | ND | 114 | ND |
| Asp-Ala | ND | ND | ND | ND | ND | 314 | ND |
| Asp-Arg | ND | ND | ND | ND | ND | 156 | ND |
| Asp-His | ND | ND | ND | ND | ND | ND | 831 |
| Asparagine | ND | ND | ND | ND | ND | ND | 830 |

| | | | | | | | |
|---|---|---|---|---|---|---|---|
| Aspartic acid | ND | 371 | 370 | ND | ND | ND | 829 |
| Avobenzone | ND | 282 | ND | ND | ND | ND | ND |
| Azelaic acid | ND | ND | ND | 175 | ND | ND | ND |
| Benzyl alcohol | ND | ND | 339 | 340 | ND | ND | ND |
| Benzyldimethyltetradecylammonium | ND | ND | 248 | 90 | ND | ND | ND |
| Benzyltrimethylammonium | ND | ND | ND | 80 | ND | ND | ND |
| Biotin | ND | 490 | ND | ND | ND | ND | ND |
| Bis(2-ethylhexyl) adipate | ND | ND | ND | 139 | ND | ND | ND |
| Butyrylcarnitine | ND | ND | ND | ND | ND | 147 | ND |
| CAR 14:2 | ND | ND | ND | ND | ND | 322 | ND |
| CAR 15:0 | ND | ND | ND | ND | ND | 378 | ND |
| CAR 16:0 | ND | ND | ND | ND | ND | 604 | ND |
| CAR 17:0 | ND | ND | ND | ND | ND | 608 | ND |
| CAR 17:1 | ND | ND | ND | ND | ND | 612 | ND |
| CAR 18:3 | ND | ND | ND | ND | ND | 97 | ND |
| CAR 20:0 | ND | ND | ND | ND | ND | 296 | ND |
| CAR 20:2 | ND | ND | ND | ND | ND | 646 | ND |
| CAR 20:3 | ND | ND | ND | ND | ND | 647 | ND |
| CAR 22:1 | ND | ND | ND | ND | ND | 191 | ND |
| CAR 22:2 | ND | ND | ND | ND | ND | 662 | ND |
| CAR 22:4 | ND | ND | ND | ND | ND | 663 | ND |
| CAR 22:5 | ND | ND | ND | ND | ND | 664 | ND |
| CAR 22:6 | ND | ND | ND | ND | ND | 665 | ND |
| CAR 24:5 | ND | ND | ND | ND | ND | 117 | ND |
| CDP ethanolamine | ND | ND | ND | ND | 225 | ND | ND |
| CE 18:1 | ND | 170 | ND | ND | 45 | ND | 828 |
| CE 18:2 | ND | ND | ND | ND | 222 | ND | 827 |
| CE 20:4 | ND | 389 | ND | ND | ND | ND | 292 |
| CE 20:5 | ND | ND | ND | ND | 354 | ND | 825 |
| CE 22:2 | ND | ND | ND | ND | 299 | ND | 824 |
| CE 22:6 | ND | 328 | ND | ND | 226 | ND | 823 |
| CL 68:2 \| CL 16:0_16:0_18:1_18:1 | ND | ND | ND | 187 | ND | ND | ND |
| CL 68:2 \| CL 16:0_18:1_16:0_18:1 | ND | ND | ND | ND | ND | 693 | ND |
| CL 68:3 \| CL 16:0_18:1_16:0_18:2 | ND | ND | ND | ND | ND | 704 | ND |
| CL 70:4 \| CL 16:0_18:1_18:1_18:2 | ND | ND | ND | ND | ND | 706 | ND |
| CL 70:5 \| CL 16:0_18:2_18:1_18:2 | ND | ND | ND | ND | ND | 709 | ND |
| CL 70:6 \| CL 16:1_18:2_18:1_18:2 | ND | ND | ND | ND | ND | 710 | ND |
| CL 70:7 \| CL 16:1_18:2_18:2_18:2 | ND | ND | ND | ND | ND | 716 | ND |
| CL 72:5 \| CL 18:1_18:1_18:1_18:2 | ND | ND | ND | ND | ND | 717 | ND |
| CL 72:6 \| CL 18:1_18:2_18:1_18:2 | ND | ND | ND | ND | ND | 718 | ND |
| CL 72:7 \| CL 18:1_18:2_18:2_18:2 | ND | ND | ND | 595 | ND | 719 | ND |
| CL 72:8 \| CL 18:2_18:2_18:2_18:2 | ND | ND | ND | 564 | ND | 4 | ND |
| CL 74:11 \| CL 16:1_18:2_18:2_22:6 | ND | ND | ND | ND | ND | 115 | ND |

| | | | | | | | |
|---|---|---|---|---|---|---|---|
| CL 74:7 \| CL 18:1_18:2_18:1_20:3 | ND | ND | ND | ND | ND | 22 | ND |
| CL 74:8 \| CL 16:1_20:3_18:1_20:3 | ND | ND | ND | ND | ND | 732 | ND |
| CL 74:9 \| CL 18:2_18:2_18:2_20:3 | ND | ND | ND | ND | ND | 733 | ND |
| CL 76:10 \| CL 18:1_18:2_18:1_22:6 | ND | ND | ND | ND | ND | 738 | ND |
| CL 76:11 \| CL 18:1_18:2_18:2_22:6 | ND | ND | ND | ND | ND | 742 | ND |
| CL 76:12 \| CL 18:2_18:2_18:2_22:6 | ND | ND | ND | ND | ND | 544 | ND |
| CL 80:15 \| CL 18:1_22:6_18:2_22:6 | ND | ND | ND | ND | ND | 360 | ND |
| CL 80:16 \| CL 18:2_22:6_18:2_22:6 | ND | ND | ND | ND | ND | 744 | ND |
| Carbamic acid, N-(2,2-diphenylacetyl)-, ethyl ester | ND | ND | 237 | ND | ND | ND | ND |
| Castanospermine | ND | 58 | ND | ND | ND | ND | 36 |
| Cer 34:1;3O \| Cer 18:1;2O/16:0;(2OH) | ND | ND | ND | 183 | ND | ND | ND |
| Cer 34:2;2O \| Cer 18:2;2O/16:0 | ND | ND | ND | ND | ND | ND | 659 |
| Cer 34:3;2O \| Cer 18:2;2O/16:1 | ND | ND | ND | ND | ND | ND | 202 |
| Cer 36:2;2O \| Cer 18:2;2O/18:0 | ND | ND | ND | ND | ND | ND | 87 |
| Cer 37:2;2O \| Cer 18:2;2O/19:0 | ND | ND | ND | ND | ND | ND | 199 |
| Cer 38:2;2O \| Cer 18:2;2O/20:0 | ND | ND | ND | ND | ND | ND | 107 |
| Cer 39:2;2O \| Cer 18:2;2O/21:0 | ND | ND | ND | ND | ND | ND | 120 |
| Cer 40:0;3O \| Cer 17:0;2O/23:0;O | ND | ND | ND | ND | ND | ND | 138 |
| Cer 41:2;2O \| Cer 18:2;2O/23:0 | ND | ND | ND | ND | ND | ND | 655 |
| Cer 42:0;3O \| Cer 17:0;2O/25:0;O | ND | ND | ND | ND | ND | ND | 653 |
| Cer 42:0;3O \| Cer 18:0;3O/24:0 | ND | ND | ND | 176 | ND | ND | ND |
| Cer 42:2;2O \| Cer 18:2;2O/24:0 | ND | ND | ND | ND | ND | ND | 610 |
| Cer 42:2;3O \| Cer 18:1;2O/24:1;(2OH) | ND | ND | ND | 134 | ND | ND | ND |
| Cer 42:3;2O \| Cer 18:2;2O/24:1 | ND | ND | ND | ND | ND | ND | 92 |
| Cer 42:4;2O \| Cer 22:3;2O/20:1 | ND | ND | ND | ND | ND | ND | 132 |
| Cer 43:2;2O \| Cer 18:1;2O/25:1 | ND | ND | ND | ND | ND | ND | 608 |
| Cer 44:0;3O \| Cer 18:0;3O/26:0 | ND | ND | ND | 156 | ND | ND | ND |
| Cer 44:1;2O \| Cer 20:1;2O/24:0 | ND | ND | ND | ND | ND | ND | 47 |
| Cer 44:2;2O \| Cer 18:2;2O/26:0 | ND | ND | ND | ND | ND | ND | 42 |
| Cer d32:1 | ND | ND | ND | ND | 122 | ND | 607 |
| Cer d33:1 | ND | ND | ND | 229 | 57 | ND | 606 |
| Cer d34:0 | ND | ND | ND | ND | 86 | ND | 605 |
| Cer d34:0 \| Cer 18:0;2O/16:0 | ND | ND | ND | 566 | ND | 313 | ND |
| Cer d34:1 | ND | 449 | ND | ND | 31 | ND | 604 |
| Cer d34:1 \| Cer 18:1;2O/16:0 | ND | ND | ND | 567 | ND | 135 | ND |
| Cer d34:2 | ND | 241 | 107 | 215 | 142 | ND | 603 |
| Cer d35:1 \| Cer 17:1;2O/18:0 | ND | ND | ND | 568 | ND | 48 | ND |
| Cer d36:0 \| Cer 18:0;2O/18:0 | ND | ND | ND | ND | ND | 746 | ND |
| Cer d36:1 | ND | 494 | ND | ND | 253 | ND | 216 |
| Cer d36:1 \| Cer 18:1;2O/18:0 | ND | ND | ND | 272 | ND | 749 | ND |
| Cer d36:2 \| Cer 18:2;2O/18:0 | ND | ND | ND | 149 | ND | 750 | ND |
| Cer d37:1 \| Cer 18:1;2O/19:0 | ND | ND | ND | ND | ND | 29 | ND |
| Cer d38:0 \| Cer 20:0;2O/18:0 | ND | ND | ND | 210 | ND | 251 | ND |

| | | | | | | | | |
|---|---|---|---|---|---|---|---|---|
| Cer d38:1 | ND | 346 | ND | ND | 77 | ND | | 602 |
| Cer d38:1 \| Cer 18:1;2O/20:0 | ND | ND | ND | 265 | ND | 755 | ND | |
| Cer d38:2 \| Cer 18:2;2O/20:0 | ND | ND | ND | 6 | ND | ND | ND | |
| Cer d39:1 | ND | ND | ND | ND | 151 | ND | | 601 |
| Cer d40:0 | ND | ND | ND | ND | 236 | ND | ND | |
| Cer d40:1 | ND | 297 | ND | ND | 219 | ND | | 448 |
| Cer d40:1 \| Cer 18:1;2O/22:0 | ND | ND | ND | 739 | ND | 764 | ND | |
| Cer d40:2 | ND | ND | ND | ND | 24 | 765 | | 551 |
| Cer d40:2 \| Cer 18:2;2O/22:0 | ND | ND | ND | 208 | ND | ND | ND | |
| Cer d41:0 \| Cer 18:0;2O/23:0 | ND | ND | ND | 170 | ND | ND | ND | |
| Cer d41:1 | ND | 386 | ND | ND | 101 | ND | | 550 |
| Cer d41:1 \| Cer 18:1;2O/23:0 | ND | ND | ND | 524 | ND | 10 | ND | |
| Cer d41:2 \| Cer 18:2;2O/23:0 | ND | ND | ND | 24 | ND | ND | ND | |
| Cer d42:0 | ND | ND | ND | ND | ND | 119 | ND | |
| Cer d42:0 \| Cer 18:0;2O/24:0 | ND | ND | ND | 223 | ND | ND | ND | |
| Cer d42:1 | ND | 572 | ND | ND | 336 | ND | | 549 |
| Cer d42:1 \| Cer 18:1;2O/24:0 | ND | ND | ND | 741 | ND | 5 | ND | |
| Cer d42:2 | ND | 426 | ND | ND | 314 | ND | | 548 |
| Cer d42:2 \| Cer 18:1;2O/24:1 | ND | ND | ND | 277 | ND | 63 | ND | |
| Cer d42:3 \| Cer 18:1;2O/24:2 | ND | ND | ND | ND | ND | 183 | ND | |
| Cer d42:3 \| Cer 18:2;2O/24:1 | ND | ND | ND | 743 | ND | ND | ND | |
| Cer d43:0 \| Cer 17:0;2O/26:0 | ND | ND | ND | ND | ND | 98 | ND | |
| Cer d43:1 | ND | ND | 64 | 168 | ND | ND | | 547 |
| Cer d43:1 \| Cer 17:1;2O/26:0 | ND | ND | ND | ND | ND | 139 | ND | |
| Cer d44:0 \| Cer 18:0;2O/26:0 | ND | ND | ND | 112 | ND | 125 | ND | |
| Cer d44:1 | ND | ND | ND | ND | ND | 279 | ND | |
| Cer d44:1 \| Cer 18:1;2O/26:0 | ND | ND | ND | 197 | ND | ND | ND | |
| Cinnamaldehyde | ND | 230 | ND | ND | ND | ND | ND | |
| CoQ10 | ND | ND | ND | ND | ND | 247 | ND | |
| CoQ8 | ND | ND | ND | ND | ND | 185 | ND | |
| CoQ9 | ND | ND | 59 | ND | ND | 358 | ND | |
| Coniferylaldehyde | ND | 107 | ND | ND | ND | ND | ND | |
| Cortisol | ND | 43 | ND | ND | ND | ND | ND | |
| Creatine phosphate | ND | ND | ND | ND | ND | 315 | ND | |
| Cyclic adenosine diphosphate ribose | ND | 479 | ND | ND | ND | ND | ND | |
| Cys-Gly | ND | ND | ND | ND | ND | ND | | 543 |
| Cystathionine | ND | 265 | ND | ND | ND | ND | | 171 |
| Cytarabine | ND | ND | ND | ND | ND | ND | | 541 |
| Cytidine | ND | 211 | 257 | 207 | ND | ND | ND | |
| Cytidine 2',3'-cyclic monophosphoric acid | ND | 208 | ND | ND | ND | ND | | 225 |
| Cytidine 5'-diphosphate ethanolamine | ND | 353 | ND | ND | ND | ND | ND | |
| Cytidine 5'-diphosphocholine | ND | 442 | ND | ND | 384 | 771 | | 540 |
| Cytidine 5'-monophosphate | ND | ND | ND | ND | 310 | ND | ND | |

| | | | | | | | |
|---|---|---|---|---|---|---|---|
| Cytidine-5'-monophosphate | ND | ND | ND | ND | ND | ND | 332 |
| D-(+)-Trehalose | ND | ND | ND | ND | 2 | ND | 459 |
| D-Fructose 6-phosphate | ND | ND | ND | ND | 120 | ND | 458 |
| D-Glucosaminic acid | ND | ND | ND | ND | ND | ND | 457 |
| D-Glucose 6-phosphate | ND | ND | ND | ND | 128 | ND | 198 |
| D-Leucyl-L-arginine | ND | ND | ND | ND | ND | ND | 456 |
| D-Ribose 5-phosphate | ND | ND | ND | ND | 293 | ND | ND |
| DAG 18:0_18:1 | ND | ND | ND | ND | 193 | ND | ND |
| DAG 18:0_22:6 | ND | ND | ND | ND | 161 | ND | ND |
| DG 30:0 | ND | ND | 166 | ND | ND | ND | ND |
| DG 30:0 \| DG 14:0_16:0 | ND | ND | ND | 681 | ND | 298 | ND |
| DG 30:1 \| DG 14:0_16:1 | ND | ND | ND | 12 | ND | 136 | ND |
| DG 30:2 \| DG 12:0_18:2 | ND | ND | ND | 192 | ND | 428 | ND |
| DG 32:0 \| DG 16:0_16:0 | ND | ND | ND | 209 | ND | 772 | ND |
| DG 32:1 | ND | 423 | 28 | 181 | 16 | 773 | 454 |
| DG 32:2 \| DG 14:0_18:2 | ND | ND | ND | 86 | ND | ND | ND |
| DG 32:2 \| DG 16:1_16:1 | ND | ND | ND | ND | ND | 776 | ND |
| DG 32:3 \| DG 14:1_18:2 | ND | ND | ND | 202 | ND | 413 | ND |
| DG 33:1 \| DG 15:0_18:1 | ND | ND | ND | 276 | ND | 777 | ND |
| DG 33:2 \| DG 15:0_18:2 | ND | ND | ND | 39 | ND | 778 | ND |
| DG 34:0 | ND | 419 | ND | ND | 325 | ND | ND |
| DG 34:0 \| DG 16:0_18:0 | ND | ND | ND | 69 | ND | ND | ND |
| DG 34:3 \| DG 16:0_18:3 | ND | ND | ND | 520 | ND | ND | ND |
| DG 34:3 \| DG 16:1_18:2 | ND | ND | ND | 280 | ND | ND | ND |
| DG 34:4 \| DG 16:1_18:3 | ND | ND | ND | 184 | ND | 781 | ND |
| DG 35:1 \| DG 17:0_18:1 | ND | ND | ND | 185 | ND | 47 | ND |
| DG 35:2 | ND | ND | ND | ND | ND | ND | 450 |
| DG 35:2 \| DG 17:0_18:2 | ND | ND | ND | 522 | ND | ND | ND |
| DG 35:2 \| DG 17:1_18:1 | ND | ND | ND | 128 | ND | 782 | ND |
| DG 35:3 | ND | ND | ND | ND | ND | ND | 449 |
| DG 35:3 \| DG 17:1_18:2 | ND | ND | ND | 194 | ND | 783 | ND |
| DG 36:0 | ND | 337 | ND | ND | ND | ND | 45 |
| DG 36:1 | ND | ND | 75 | ND | 540 | 81 | 191 |
| DG 36:1 \| DG 18:0_18:1 | ND | ND | ND | 243 | ND | ND | ND |
| DG 36:2 \| DG 18:0_18:2 | ND | ND | ND | 16 | ND | ND | ND |
| DG 36:4 \| DG 16:0_20:4 | ND | ND | ND | 48 | ND | ND | ND |
| DG 37:2 | ND | ND | ND | 151 | ND | ND | ND |
| DG 37:3 \| DG 19:1_18:2 | ND | ND | ND | 486 | ND | 790 | ND |
| DG 37:7 | ND | ND | ND | 487 | ND | ND | 80 |
| DG 37:8 | ND | ND | ND | 232 | ND | ND | ND |
| DG 38:1 \| DG 20:0_18:1 | ND | ND | ND | ND | ND | 52 | ND |
| DG 38:2 | ND | ND | ND | ND | ND | ND | 243 |
| DG 38:2 \| DG 18:1_20:1 | ND | ND | ND | 122 | ND | 791 | ND |

| | | | | | | | |
|---|---|---|---|---|---|---|---|
| DG 38:3 \| DG 20:1_18:2 | ND | ND | ND | 245 | ND | 793 | ND |
| DG 38:4 \| DG 18:0_20:4 | ND | ND | ND | 488 | ND | ND | ND |
| DG 38:4 \| DG 18:2_20:2 | ND | ND | ND | ND | ND | 303 | ND |
| DG 38:5 \| DG 16:0_22:5 | ND | ND | ND | 152 | ND | ND | ND |
| DG 38:6 | ND | ND | ND | ND | 306 | ND | 125 |
| DG 38:6 \| DG 16:0_22:6 | ND | ND | ND | 100 | ND | 53 | ND |
| DG 38:7 \| DG 16:1_22:6 | ND | ND | ND | 489 | ND | ND | ND |
| DG 39:7 | ND | ND | 50 | 547 | ND | ND | ND |
| DG 39:8 | ND | ND | 65 | ND | ND | ND | ND |
| DG 39:9 | ND | ND | ND | 516 | ND | ND | ND |
| DG 40:3 \| DG 22:1_18:2 | ND | ND | ND | ND | ND | 797 | ND |
| DG 40:5 | ND | ND | ND | ND | ND | ND | 182 |
| DG 40:5 \| DG 18:0_22:5 | ND | ND | ND | ND | ND | 118 | ND |
| DG 40:6 \| DG 18:1_22:5 | ND | ND | ND | 517 | ND | 798 | ND |
| DG 40:7 | ND | ND | ND | ND | ND | ND | 336 |
| DG 40:7 \| DG 18:2_22:5 | ND | ND | ND | 433 | ND | ND | ND |
| DG 40:9 \| DG 18:3_22:6 | ND | ND | ND | ND | ND | 803 | ND |
| DG 41:10 | ND | ND | ND | 2 | ND | ND | ND |
| DG 41:6 | ND | ND | ND | 435 | ND | ND | ND |
| DG 41:7 | ND | ND | 130 | 59 | ND | 2 | ND |
| DG 41:8 | ND | ND | 259 | 436 | ND | 1 | ND |
| DG 41:9 | ND | ND | 92 | 255 | ND | 804 | ND |
| DG 42:7 | ND | ND | ND | 437 | ND | ND | 184 |
| DG 43:10 | ND | ND | 4 | 127 | ND | 44 | ND |
| DG 43:11 | ND | ND | 35 | 514 | ND | ND | ND |
| DG 43:8 | ND | ND | 274 | 3 | ND | 15 | ND |
| DG 43:9 | ND | ND | 80 | 515 | ND | 805 | ND |
| DG 44:8 | ND | ND | ND | 485 | ND | ND | ND |
| DG 44:9 | ND | ND | 167 | ND | ND | 809 | ND |
| DG 45:12 | ND | ND | ND | 218 | ND | ND | ND |
| DG 45:8 | ND | ND | ND | 174 | ND | ND | ND |
| DG 47:7 | ND | ND | ND | 375 | ND | ND | ND |
| DL-2-Aminocaprylic acid | ND | ND | ND | ND | ND | ND | 335 |
| DL-Arginine | ND | ND | ND | ND | ND | ND | 334 |
| DL-Lanthionine | ND | ND | ND | ND | ND | ND | 91 |
| DL-Tryptophan, methyl ester | ND | ND | ND | ND | ND | ND | 297 |
| DMPE 36:1 | ND | ND | ND | ND | ND | ND | 333 |
| DMPE 40:6 | ND | ND | ND | ND | ND | ND | 942 |
| Decanoyl-carnitine | ND | 81 | ND | ND | ND | ND | ND |
| Diethyl (carboxymethylamino)methylenemalonate | ND | ND | ND | 376 | ND | ND | ND |
| Diethylcarbamazine | ND | 7 | ND | ND | ND | ND | ND |
| Dimethyl sulfoxide | ND | ND | ND | ND | ND | 150 | ND |
| Dimethyllysine | ND | ND | ND | ND | 522 | ND | ND |

| | | | | | | | |
|---|---|---|---|---|---|---|---|
| Dimetilan | ND | ND | ND | ND | ND | 811 | ND |
| Docosahexaenoylglycine | ND | 522 | ND | ND | ND | ND | ND |
| Dodeca-2(E),4(E)-dienoic acid | ND | ND | 207 | ND | ND | ND | ND |
| Dodecanoic acid, 12-[[(cyclohexylamino)carbonyl]amino]- | ND | ND | 341 | ND | ND | ND | ND |
| Ecgonine | ND | ND | ND | ND | 324 | ND | ND |
| Ethanolamine | ND | ND | 325 | 378 | ND | 293 | ND |
| Ethyl 2-acetylpentanoate | ND | ND | ND | 154 | ND | ND | ND |
| Ethyl 4-amino-1-piperidinecarboxylate | ND | 387 | ND | ND | ND | ND | ND |
| Ethyl 4-oxo-3-piperidinecarboxylate | ND | ND | ND | 47 | ND | ND | ND |
| FA 10:0 | ND | 91 | ND | ND | 78 | ND | 37 |
| FA 11:0 | ND | 345 | ND | ND | 202 | ND | 939 |
| FA 12:0 | ND | 88 | ND | ND | 17 | ND | 938 |
| FA 13:0 | ND | 234 | ND | ND | 8 | ND | 163 |
| FA 14:0 | ND | ND | ND | ND | 339 | ND | 937 |
| FA 14:0 (myristic acid) | ND | ND | ND | 452 | ND | ND | ND |
| FA 14:1 (physeteric acid) | ND | ND | ND | 159 | ND | ND | ND |
| FA 15:4 | ND | ND | ND | ND | ND | 288 | ND |
| FA 16:0 | ND | 151 | ND | 275 | 538 | ND | 934 |
| FA 16:1 (palmitoleic acid) | ND | ND | ND | 453 | ND | 812 | ND |
| FA 16:3 | ND | ND | ND | 430 | ND | 817 | ND |
| FA 16:4 | ND | ND | ND | 111 | ND | 818 | ND |
| FA 17:0 (margaric acid) | ND | ND | ND | 431 | ND | ND | ND |
| FA 17:2 | ND | ND | ND | 329 | ND | 836 | ND |
| FA 18:0 | ND | 145 | ND | ND | 319 | ND | 932 |
| FA 18:1 (oleic acid) | ND | ND | ND | ND | ND | 848 | ND |
| FA 18:1; (oleic acid) | ND | ND | ND | 330 | ND | ND | ND |
| FA 18:2 (linoleic acid) | ND | ND | ND | 41 | ND | 866 | ND |
| FA 18:3 (linolenic acid) | ND | ND | ND | 331 | ND | 867 | ND |
| FA 18:5 | ND | ND | ND | ND | ND | 876 | ND |
| FA 19:2 | ND | ND | ND | 398 | ND | 879 | ND |
| FA 20:0 | ND | 316 | ND | ND | 245 | ND | ND |
| FA 20:1 (eicosenoic acid) | ND | ND | ND | 370 | ND | 885 | ND |
| FA 20:2 (eicosadienoic acid) | ND | ND | ND | 371 | ND | 886 | ND |
| FA 20:3 (homo-gamma-linolenic acid) | ND | ND | ND | 372 | ND | 887 | ND |
| FA 20:4 (arachidonic acid) | ND | ND | ND | 189 | ND | 888 | ND |
| FA 20:5 (eicosapentaenoic acid) | ND | ND | ND | 373 | ND | 899 | ND |
| FA 21:1 | ND | ND | ND | 264 | ND | 406 | ND |
| FA 22:1 (erucic acid) | ND | ND | ND | 325 | ND | ND | ND |
| FA 22:2 (docosadienoic acid) | ND | ND | ND | 326 | ND | 903 | ND |
| FA 22:3 | ND | ND | ND | 327 | ND | 907 | ND |
| FA 22:4 | ND | ND | ND | 240 | ND | 912 | ND |
| FA 22:6 (docosahexaenoic acid) | ND | ND | ND | 249 | ND | ND | ND |
| FA 23:1 | ND | ND | ND | 186 | ND | 913 | ND |

| | | | | | | | |
|---|---|---|---|---|---|---|---|
| FA 24:1 (nervonic acid) | ND | ND | ND | 268 | ND | 922 | ND |
| FA 24:2 | ND | ND | ND | 125 | ND | 93 | ND |
| FA 24:4 | ND | ND | ND | 123 | ND | ND | ND |
| FA 25:1 | ND | ND | ND | 355 | ND | ND | ND |
| FA 26:1 | ND | ND | ND | 135 | ND | ND | ND |
| FA 40:6 | ND | ND | ND | 740 | ND | ND | ND |
| FA 44:10 | ND | ND | ND | 7 | ND | ND | ND |
| FAD | ND | 352 | ND | ND | 521 | ND | 671 |
| FAHFA 34:1;O \| FAHFA 16:0/18:1;O | ND | ND | ND | 725 | ND | ND | ND |
| GM3 36:1;2O | ND | ND | ND | ND | ND | 925 | ND |
| Galacto-N-biose | ND | ND | ND | ND | 72 | ND | 945 |
| Gentiobiose | ND | ND | ND | ND | ND | ND | 943 |
| Geranic acid | ND | 39 | ND | ND | ND | ND | ND |
| GlcCer d34:1 | ND | ND | ND | ND | ND | ND | 319 |
| GlcCer d40:1 | ND | 400 | ND | 726 | 107 | ND | 318 |
| GlcCer d41:1 | ND | ND | ND | ND | 165 | ND | ND |
| GlcCer d42:1 | ND | 298 | ND | ND | 241 | ND | 307 |
| GlcCer d42:2 | ND | 415 | ND | 343 | 364 | ND | 277 |
| Gln-Arg | ND | ND | ND | ND | ND | 386 | ND |
| Gln-Gln | ND | ND | ND | ND | ND | 149 | ND |
| Gln-Glu | ND | 527 | ND | ND | ND | 176 | ND |
| Gln-Lys | ND | ND | ND | ND | 74 | 926 | ND |
| Gln-Thr | ND | 500 | ND | ND | 484 | 929 | 316 |
| Glu-Ala | ND | 454 | ND | ND | ND | ND | ND |
| Glu-Asp | ND | 257 | ND | ND | ND | ND | ND |
| Glu-Gln | ND | 385 | ND | ND | ND | ND | 13 |
| Glu-Gln.1 | ND | 210 | ND | ND | ND | ND | ND |
| Glu-Glu-Arg | ND | 175 | ND | ND | ND | ND | ND |
| Glu-Gly-Arg | ND | ND | ND | 344 | ND | ND | 259 |
| Glu-His | ND | 520 | ND | ND | ND | ND | 315 |
| Glu-Phe | ND | ND | ND | ND | ND | ND | 314 |
| Glu-Pro-Arg | ND | ND | ND | ND | ND | 931 | ND |
| Glu-Ser | ND | ND | ND | ND | ND | 327 | ND |
| Glucose-1-phosphate | ND | ND | ND | ND | 218 | ND | 312 |
| Glucose-6-phosphate | ND | ND | ND | 315 | ND | ND | ND |
| Glutarylcarnitine | ND | 78 | ND | ND | ND | ND | ND |
| Glutathione (oxidized) | ND | 570 | ND | ND | 139 | 379 | 928 |
| Glutathionesulfonic acid | ND | 333 | ND | ND | ND | ND | 223 |
| Gly-Arg | ND | 451 | ND | ND | 166 | ND | 331 |
| Gly-Gly-Gly | ND | 437 | ND | ND | ND | ND | 330 |
| Gly-His | ND | ND | ND | ND | 392 | ND | 329 |
| Gly-Lys | ND | ND | ND | ND | 211 | 932 | 328 |
| Gly-Pro-Lys | ND | ND | ND | 318 | ND | ND | ND |

| | | | | | | | | |
|---|---|---|---|---|---|---|---|---|
| Gly-val | ND | ND | ND | ND | 39 | ND | | 326 |
| Glycerol 1-myristate | ND | 355 | ND | ND | ND | ND | ND | |
| Glycine | ND | ND | 381 | 320 | ND | ND | ND | |
| Goralatide | ND | 409 | ND | ND | ND | ND | ND | |
| Guanidinosuccinic acid | ND | ND | ND | ND | 229 | 934 | | 240 |
| Guanosine | ND | 357 | ND | ND | 279 | ND | | 322 |
| Guanosine 5'-monophosphate | ND | 285 | ND | ND | 173 | ND | | 321 |
| HMBA | ND | 468 | ND | ND | ND | ND | ND | |
| Heptadecasphing-4-enine | ND | ND | 262 | ND | ND | 399 | ND | |
| Heptadecasphinganine | ND | 8 | ND | ND | ND | ND | ND | |
| Heptanedioic acid, 1-(2-cyclopentylidenehydrazide) | ND | ND | 297 | ND | ND | ND | ND | |
| Hex2Cer 40:1 | ND | ND | ND | ND | ND | ND | | 320 |
| Hex2Cer 40:2 | ND | ND | ND | ND | ND | ND | | 210 |
| Hex2Cer 41:1 | ND | ND | ND | ND | ND | ND | | 354 |
| Hex2Cer 41:2 | ND | ND | ND | ND | ND | ND | | 353 |
| Hex2Cer 42:1 | ND | ND | ND | ND | ND | ND | | 352 |
| Hex2Cer 42:3 | ND | ND | ND | ND | ND | ND | | 351 |
| Hex2Cer 43:2 | ND | ND | ND | ND | ND | ND | | 303 |
| Hex3Cer 34:1 | ND | ND | ND | ND | ND | ND | | 350 |
| Hex3Cer 34:2 | ND | ND | ND | ND | ND | ND | | 349 |
| Hex3Cer 38:1 | ND | ND | ND | ND | ND | ND | | 348 |
| Hex3Cer 40:1 | ND | ND | ND | ND | ND | ND | | 347 |
| Hex3Cer 40:2 | ND | ND | ND | ND | ND | ND | | 346 |
| Hex3Cer 41:1 | ND | ND | ND | ND | ND | ND | | 345 |
| Hex3Cer 41:2 | ND | ND | ND | ND | ND | ND | | 344 |
| Hex3Cer 42:1 | ND | ND | ND | ND | ND | ND | | 343 |
| Hex3Cer 42:2 | ND | ND | ND | ND | ND | ND | | 342 |
| Hex3Cer 42:3 | ND | ND | ND | ND | ND | ND | | 341 |
| HexCer 34:2 | ND | ND | ND | ND | ND | ND | | 340 |
| HexCer 36:1;2O | ND | ND | ND | 321 | ND | ND | ND | |
| HexCer 36:1;2O\|HexCer 18:1;2O/18:0 | ND | ND | ND | ND | ND | 936 | ND | |
| HexCer 38:1 | ND | ND | ND | ND | ND | ND | | 188 |
| HexCer 38:1;2O | ND | ND | ND | 322 | ND | ND | ND | |
| HexCer 38:1;2O\|HexCer 18:1;2O/20:0 | ND | ND | ND | ND | ND | 938 | ND | |
| HexCer 38:1;3O | ND | ND | ND | ND | ND | 939 | ND | |
| HexCer 38:1;3O\|HexCer 16:1;2O/22:0;O | ND | ND | ND | 323 | ND | ND | ND | |
| HexCer 38:1;3O\|HexCer 18:1;2O/20:0;O | ND | ND | ND | ND | ND | 940 | ND | |
| HexCer 40:0 | ND | ND | ND | ND | ND | ND | | 66 |
| HexCer 40:0;2O | ND | ND | ND | 324 | ND | 941 | ND | |
| HexCer 40:0;2O\|HexCer 18:0;2O/22:0 | ND | ND | ND | ND | ND | 942 | ND | |
| HexCer 40:0;3O | ND | ND | ND | ND | ND | 947 | ND | |
| HexCer 40:1;3O | ND | ND | ND | ND | ND | 953 | ND | |
| HexCer 40:1;3O\|HexCer 16:1;2O/24:0;O | ND | ND | ND | 288 | ND | ND | ND | |

| | | | | | | | |
|---|---|---|---|---|---|---|---|
| HexCer 40:1;3O\|HexCer 18:1;2O/22:0;O | ND | ND | ND | ND | ND | 959 | ND |
| HexCer 40:2;2O\|HexCer 18:1;2O/22:1 | ND | ND | ND | ND | ND | 960 | ND |
| HexCer 41:1;2O | ND | ND | ND | 289 | ND | ND | ND |
| HexCer 41:1;2O\|HexCer 18:1;2O/23:0 | ND | ND | ND | ND | ND | 962 | ND |
| HexCer 41:1;3O | ND | ND | ND | ND | ND | 963 | ND |
| HexCer 41:1;3O\|HexCer 16:1;2O/25:0;O | ND | ND | ND | 290 | ND | ND | ND |
| HexCer 41:1;3O\|HexCer 18:1;2O/23:0;O | ND | ND | ND | ND | ND | 965 | ND |
| HexCer 41:2 | ND | ND | ND | ND | ND | ND | 137 |
| HexCer 41:3 | ND | ND | ND | ND | ND | ND | 366 |
| HexCer 42:0 | ND | ND | ND | ND | ND | ND | 275 |
| HexCer 42:0;2O | ND | ND | ND | 291 | ND | ND | ND |
| HexCer 42:0;2O\|HexCer 18:0;2O/24:0 | ND | ND | ND | ND | ND | 966 | ND |
| HexCer 42:0;3O | ND | ND | ND | ND | ND | 967 | ND |
| HexCer 42:1;2O\|HexCer 18:1;2O/24:0 | ND | ND | ND | 292 | ND | 968 | ND |
| HexCer 42:1;3O | ND | ND | ND | ND | ND | 969 | ND |
| HexCer 42:1;3O\|HexCer 16:1;2O/26:0;O | ND | ND | ND | 293 | ND | ND | ND |
| HexCer 42:1;3O\|HexCer 18:1;2O/24:0;O | ND | ND | ND | ND | ND | 970 | ND |
| HexCer 42:2;3O | ND | ND | ND | ND | ND | 973 | ND |
| HexCer 42:2;3O\|HexCer 16:1;2O/26:1;O | ND | ND | ND | 295 | ND | ND | ND |
| HexCer 42:2;3O\|HexCer 18:1;2O/24:1;O | ND | ND | ND | ND | ND | 974 | ND |
| HexCer 42:3;2O | ND | ND | ND | 296 | ND | 975 | ND |
| HexCer 42:3;2O\|HexCer 18:1;2O/24:2 | ND | ND | ND | ND | ND | 976 | ND |
| Hexadecyltrimethylammonium | ND | ND | 272 | 297 | ND | ND | ND |
| Hexamethylcyclotrisiloxane | ND | 268 | ND | ND | ND | ND | ND |
| His-Ala | ND | 179 | ND | 298 | ND | 978 | 363 |
| His-Asn | ND | ND | ND | ND | ND | ND | 362 |
| His-Asp | ND | 432 | ND | ND | ND | ND | 361 |
| His-Gln | ND | 511 | ND | ND | 70 | 979 | 360 |
| His-Glu | ND | 531 | ND | ND | ND | ND | 359 |
| His-Gly | ND | ND | ND | ND | ND | 330 | 358 |
| His-Leu | ND | 129 | ND | ND | ND | ND | ND |
| His-Met | ND | ND | ND | ND | ND | 216 | ND |
| His-Ser | ND | 493 | ND | ND | ND | ND | 356 |
| His-Thr | ND | ND | ND | ND | 257 | ND | 355 |
| His-Tyr | ND | 5 | ND | ND | ND | 377 | ND |
| His-Val | ND | 484 | ND | 299 | ND | 980 | 380 |
| Histamine | ND | 34 | 367 | 300 | ND | 69 | ND |
| Hydroxylysine | ND | ND | ND | ND | ND | ND | 254 |
| Ile-Arg | ND | 394 | ND | 308 | ND | ND | ND |
| Ile-Glu | ND | ND | ND | ND | ND | 984 | ND |
| Ile-Gly-Lys | ND | ND | ND | ND | ND | ND | 377 |
| Ile-His | ND | ND | ND | ND | 413 | 49 | 376 |
| Ile-Ile | ND | ND | ND | ND | 414 | 26 | ND |

| | | | | | | | |
|---|---|---|---|---|---|---|---|
| Ile-Met | ND | ND | ND | ND | ND | 38 | ND |
| Ile-Pro | ND | ND | ND | ND | ND | ND | 374 |
| Ile-Pro-Arg | ND | 498 | ND | ND | ND | ND | ND |
| Ile-Pro-Ile | ND | ND | ND | ND | ND | 54 | ND |
| Ile-Ser | ND | ND | ND | ND | ND | 986 | ND |
| Indarubicin | ND | ND | ND | 91 | ND | ND | ND |
| Inosine 5'-monophosphate | ND | ND | ND | ND | ND | 987 | ND |
| Inosine-5'-monophosphate | ND | 101 | ND | ND | 19 | ND | 372 |
| Isoleucine | ND | ND | 383 | 310 | ND | ND | ND |
| Isoproturon | ND | ND | 266 | ND | ND | ND | ND |
| Isouron | ND | 71 | ND | ND | ND | ND | ND |
| Isovaleryl-L-carnitine | ND | ND | ND | ND | 285 | ND | 43 |
| Ketamine | ND | ND | ND | 34 | ND | ND | ND |
| L-.beta.-Homoglutamine | ND | ND | ND | ND | ND | ND | 371 |
| L-.beta.-Homolysine | ND | ND | ND | ND | ND | ND | 370 |
| L-.beta.-Homomethionine | ND | ND | ND | ND | ND | ND | 369 |
| L-.gamma.-Glutamyl-L-glutamic acid | ND | ND | ND | ND | ND | ND | 368 |
| L-Arginine, methyl ester | ND | ND | ND | ND | ND | ND | 367 |
| L-Cysteine S-sulfate | ND | ND | ND | ND | ND | ND | 392 |
| L-Cysteine-glutathione disulfide | ND | ND | ND | ND | 323 | ND | 272 |
| L-Glutamic acid, dimethyl ester | ND | ND | ND | ND | ND | ND | 391 |
| L-Homocitrulline | ND | ND | ND | ND | ND | ND | 389 |
| L-Leucine, methyl ester | ND | ND | 242 | ND | ND | ND | ND |
| L-Phenylalanine, methyl ester | ND | ND | ND | 311 | ND | ND | ND |
| L-Proline | ND | ND | ND | 312 | ND | ND | ND |
| L-Propionylcarnitine | ND | 330 | ND | ND | ND | ND | 388 |
| L-Saccharopine | ND | ND | ND | ND | 284 | ND | 387 |
| L-Tryptophan | ND | ND | ND | 313 | ND | ND | 386 |
| L-Valinamide | ND | ND | 361 | 177 | ND | ND | ND |
| L-gamma-Glutamyl-L-glutamic acid | ND | ND | ND | ND | 280 | ND | ND |
| LPC 17:1 | ND | ND | ND | ND | 382 | ND | 385 |
| LPC 20:0 | ND | ND | ND | ND | 553 | ND | 384 |
| LPC 20:1 | ND | ND | ND | ND | 294 | ND | 383 |
| LPC 20:2 | ND | ND | ND | ND | ND | ND | 382 |
| LPC 22:6 | ND | ND | ND | ND | 554 | ND | ND |
| LPC 14:0 | ND | 507 | ND | ND | 288 | 990 | 294 |
| LPC 15:0 | ND | ND | ND | ND | 555 | 348 | 148 |
| LPC 16:0 | ND | 244 | ND | 314 | 557 | 991 | 381 |
| LPC 16:1 | ND | 284 | ND | 285 | 343 | 993 | 193 |
| LPC 17:0 | ND | ND | ND | ND | 373 | 994 | ND |
| LPC 17:1 | ND | ND | ND | ND | ND | 995 | ND |
| LPC 18:2 | ND | 362 | ND | 220 | 544 | 998 | 406 |
| LPC 18:3 | ND | ND | ND | ND | 400 | 1000 | 405 |

| | | | | | | | |
|---|---|---|---|---|---|---|---|
| LPC 19:0 | ND | ND | ND | ND | ND | 86 | ND |
| LPC 20:0 | ND | 446 | ND | ND | ND | 271 | ND |
| LPC 20:1 | ND | 413 | ND | ND | ND | ND | ND |
| LPC 20:2 | ND | 334 | ND | ND | 290 | ND | ND |
| LPC 20:3 | ND | 186 | ND | ND | 417 | 1001 | 404 |
| LPC 20:4 | ND | 260 | ND | 106 | 571 | 1005 | ND |
| LPC 20:4.1 | ND | ND | ND | ND | ND | 1007 | ND |
| LPC 20:5 | ND | 216 | ND | ND | ND | 270 | 403 |
| LPC 22:4 | ND | 144 | ND | ND | 572 | ND | ND |
| LPC 22:5 | ND | ND | ND | ND | 224 | 401 | 402 |
| LPC 22:5.1 | ND | ND | ND | ND | ND | 389 | ND |
| LPC 22:6 | ND | 312 | ND | 246 | ND | 1008 | ND |
| LPC 22:6.1 | ND | ND | ND | ND | ND | 1011 | ND |
| LPC 24:0 | ND | ND | ND | ND | ND | 355 | ND |
| LPE 16:0.1 | ND | ND | ND | ND | 315 | 1021 | ND |
| LPE 22:5 | ND | ND | ND | ND | ND | 65 | ND |
| LPE 22:6.1 | ND | ND | ND | ND | ND | 1034 | ND |
| LPE O-18:1 | ND | ND | ND | 302 | ND | 1038 | ND |
| LPG 16:0 | ND | ND | ND | ND | ND | 1042 | ND |
| LPS 18:0 | ND | ND | ND | 305 | ND | 1051 | ND |
| LPS 22:6 | ND | ND | ND | ND | ND | 1052 | ND |
| Lactosylceramide d42:2 | ND | ND | ND | ND | ND | ND | 396 |
| Lactoylglutathione | ND | ND | ND | ND | ND | 1053 | ND |
| Leu-Ala | ND | ND | ND | ND | ND | ND | 395 |
| Leu-Arg | ND | ND | ND | ND | 141 | 1054 | 394 |
| Leu-Lys | ND | ND | ND | ND | 103 | 1056 | 393 |
| Leu-Pro | ND | ND | ND | ND | ND | ND | 421 |
| Leu-Pro-Arg | ND | 568 | ND | ND | ND | ND | ND |
| Leu-Ser | ND | ND | ND | ND | ND | 214 | ND |
| Leu-Val | ND | 283 | ND | ND | ND | ND | ND |
| Leucine | ND | ND | 384 | 306 | 574 | ND | ND |
| Linoleoylcarnitine | ND | ND | ND | ND | 109 | ND | ND |
| Lumichrome | ND | ND | ND | ND | ND | ND | 29 |
| Lys-Asn | ND | 532 | ND | ND | ND | ND | 419 |
| Lys-Gln | ND | ND | ND | ND | ND | ND | 418 |
| Lys-Glu | ND | 564 | ND | ND | ND | 1057 | 417 |
| Lys-Gly | ND | 529 | ND | ND | ND | ND | ND |
| Lys-Phe | ND | 109 | ND | ND | ND | ND | ND |
| Lys-Pro | ND | ND | ND | ND | ND | 297 | 416 |
| Lys-Thr | ND | 359 | ND | ND | ND | ND | ND |
| Lys-Tyr | ND | 108 | ND | ND | ND | ND | ND |
| Lys-Val | ND | 27 | ND | 347 | ND | 1059 | ND |
| Lysine | ND | 3 | 393 | 348 | 169 | 113 | 415 |

| | | | | | | | | |
|---|---|---|---|---|---|---|---|---|
| Malonyl-L-carnitine_Malonyl-L-carnitine | ND | ND | ND | ND | 189 | ND | ND | |
| Malonyl-carnitine | ND | ND | ND | ND | ND | 1060 | ND | |
| Maltose | ND | ND | ND | ND | 6 | ND | | 21 |
| Maltotetraose | ND | ND | ND | ND | ND | ND | | 413 |
| Maltotriose | ND | ND | ND | ND | 15 | ND | | 412 |
| Mannosamine | ND | ND | ND | ND | 136 | ND | ND | |
| Melamine | ND | ND | ND | ND | ND | 1061 | ND | |
| Memantine | ND | ND | ND | ND | ND | ND | | 411 |
| Met-Arg | ND | ND | ND | ND | ND | 1062 | ND | |
| Met-Phe | ND | ND | ND | ND | ND | 359 | ND | |
| Metanephrine | ND | ND | ND | ND | ND | ND | | 295 |
| Metformin | ND | ND | 201 | 281 | ND | ND | ND | |
| Methacholine cation | ND | ND | ND | ND | ND | ND | | 410 |
| Methadone | ND | 306 | ND | ND | ND | ND | ND | |
| Methionine | ND | 562 | 394 | 349 | 402 | ND | | 409 |
| Methioninesulfoxide A | ND | ND | 316 | ND | ND | ND | ND | |
| Methyl 1-piperazinecarboxylate | ND | 526 | ND | ND | ND | ND | ND | |
| Methyl DL-pyroglutamate | ND | ND | ND | ND | ND | ND | | 435 |
| Methyl jasmonate | ND | 276 | ND | 20 | ND | ND | ND | |
| Methyl-histidine_b | ND | ND | ND | ND | ND | 336 | ND | |
| Methylecgonine | ND | ND | ND | ND | ND | ND | | 434 |
| Methylpropionic acid | ND | ND | 142 | 173 | ND | ND | ND | |
| Metoprolol acid | ND | 513 | ND | ND | ND | ND | ND | |
| Miglitol | ND | ND | 120 | ND | ND | ND | ND | |
| Muramic acid | ND | ND | ND | ND | ND | ND | | 433 |
| Muscarine | ND | ND | ND | ND | ND | 131 | ND | |
| Myricetin | ND | ND | ND | ND | 146 | ND | ND | |
| Myristamidopropyl dimethylamine | ND | ND | ND | ND | 250 | ND | ND | |
| Myristoyl ethanolamide | ND | 356 | ND | ND | ND | ND | ND | |
| Myristoyl-carnitine | ND | ND | ND | ND | ND | 1064 | ND | |
| N,N'-Diacetylcystine | ND | 561 | ND | ND | ND | ND | ND | |
| N,N-Dibutyl-N'-(3-chloro-2-methylphenyl)urea | ND | ND | ND | 143 | ND | ND | ND | |
| N,N-Diethyl-2-aminoethanol | ND | 15 | ND | ND | ND | ND | ND | |
| N-(1-Amino-3,3-dimethyl-1-oxobutan-2-yl)-1-pentyl-1H-indole-3-carboxamide | ND | ND | ND | 62 | ND | ND | ND | |
| N-(15Z-Tetracosenoyl)-1-.beta.-galactosylsphing-4-enine | ND | 427 | ND | ND | ND | ND | ND | |
| N-(2'-Oxobi(cyclohex)yl)acetamide | ND | ND | 44 | ND | ND | ND | ND | |
| N-(2-Chlorobenzyl)-2-propanamine | ND | ND | ND | ND | 137 | ND | ND | |
| N-(3-(Aminomethyl)benzyl)acetamidine | ND | ND | ND | 165 | ND | ND | ND | |
| N-(3-Methoxypropyl)-9H-purin-6-amine | ND | ND | ND | 352 | ND | ND | ND | |
| N-(5-Amino-1,3,4-thiadiazol-2-yl)methanesulfonamide | ND | ND | ND | 213 | ND | ND | ND | |
| N-(Octadecanoyl)sphing-4-enine-1-phosphocholine | ND | ND | ND | ND | ND | 1065 | ND | |
| N-(Piperidin-4-yl)methanesulfonamide | ND | ND | 215 | ND | ND | ND | ND | |
| N-(tert-Butyl)-4-phenylbutanamide | ND | ND | 192 | ND | ND | ND | ND | |

| | | | | | | | |
|---|---|---|---|---|---|---|---|
| N-.alpha.-(tert-Butoxycarbonyl)-L-histidine | ND | ND | 390 | 360 | ND | ND | ND |
| N-.alpha.-Acetyl-L-arginine | ND | ND | ND | ND | ND | ND | 429 |
| N-.alpha.-Acetyl-L-ornithine | ND | ND | 43 | 361 | ND | ND | 428 |
| N-3-Hydroxydecanoyl-L-homoserine lactone | ND | ND | ND | ND | ND | ND | 427 |
| N-Acetyl-D-galactosamine | ND | ND | ND | ND | ND | ND | 426 |
| N-Acetyl-D-galactosamine-6-phosphate | ND | ND | ND | ND | ND | ND | 425 |
| N-Acetyl-D-galactosaminitol | ND | ND | ND | ND | ND | ND | 424 |
| N-Acetyl-D-glucosamine 6-phosphate | ND | ND | ND | ND | 200 | ND | ND |
| N-Acetyl-D-lactosamine | ND | ND | ND | ND | ND | ND | 422 |
| N-Acetyl-L-Prolinamide | ND | ND | ND | 221 | ND | ND | ND |
| N-Acetyl-L-carnosine | ND | 322 | ND | ND | ND | 1066 | ND |
| N-Acetyl-lactosamine | ND | ND | ND | ND | ND | 210 | ND |
| N-Acetyl-leucine | ND | ND | 161 | ND | ND | ND | ND |
| N-Acetyl-methionine | ND | 528 | ND | ND | ND | ND | ND |
| N-Acetylhistamine | ND | ND | ND | ND | ND | 129 | ND |
| N-Acetylneuraminic acid, 2,3-dehydro-2-deoxy- | ND | 105 | ND | ND | ND | ND | ND |
| N-Arachidonoyl-gamma-aminobutyric acid | ND | 499 | ND | ND | ND | ND | ND |
| N-Arachidonoyltaurine | ND | 514 | ND | ND | ND | ND | ND |
| N-Benzyl-N,N-dimethyl-1-hexadecanaminium | ND | ND | ND | 279 | ND | ND | ND |
| N-Carboxyethyl-.gamma.-aminobutyric acid | ND | 41 | ND | ND | ND | ND | ND |
| N-Cyclohexyl-N'-[2-(1H-imidazol-4-yl)ethyl]urea | ND | ND | ND | 365 | ND | ND | ND |
| N-Cyclohexylcyclohexanecarboxamide | ND | 163 | ND | ND | ND | ND | ND |
| N-Decyl-N,N-dimethyl-1-decanaminium | ND | ND | ND | 155 | ND | ND | ND |
| N-Desmethylvenlafaxine | ND | ND | 241 | ND | ND | ND | ND |
| N-Formylnornicotine | ND | ND | ND | ND | ND | 1068 | ND |
| N-Hydroxy-3,4-methylenedioxyamphetamine | ND | 351 | ND | ND | ND | ND | ND |
| N-Isobutyl-3-methylbutanamide | ND | ND | ND | ND | ND | 402 | ND |
| N-Methyl-asparagine | ND | ND | 227 | ND | ND | ND | ND |
| N-Methyl-histidine | ND | ND | ND | ND | ND | 1069 | ND |
| N-Methyl-serine | ND | ND | ND | ND | ND | 1070 | ND |
| N-Methylleucine | ND | ND | ND | ND | ND | ND | 7 |
| N-Methylphenylalanine | ND | ND | ND | ND | ND | ND | 446 |
| N-Nervonoyl-D-erythro-sphingosylphosphorylcholine | ND | ND | ND | ND | ND | 387 | ND |
| N-Nitroimidazolidin-2-imine | ND | ND | ND | ND | ND | ND | 445 |
| N-[2-(4-Methoxyphenyl)ethyl]-3-methyl-2-butenamide | ND | ND | 358 | 369 | ND | ND | ND |
| N-[3-(1H-Imidazol-1-yl)propyl]-2-(4-methylphenyl)acetamide | ND | ND | ND | 250 | ND | ND | ND |
| N-acetyl-D-galactosamine | ND | ND | ND | ND | 75 | ND | ND |
| N-acetyl-D-mannosamine | ND | ND | ND | ND | 175 | ND | ND |
| N-acetylaspartic acid | ND | 126 | ND | ND | ND | 1071 | ND |
| N-acetylmannosamine | ND | ND | ND | ND | ND | ND | 444 |
| N-acetylmethionine | ND | ND | ND | ND | ND | 1077 | ND |
| N-alpha-Acetyl-L-arginine | ND | ND | ND | ND | 33 | ND | ND |
| N-epsilon-dimethyl-lysine | ND | ND | ND | ND | ND | 1082 | ND |

| | | | | | | | |
|---|---|---|---|---|---|---|---|
| N-n-Butylpropionamide | ND | 541 | ND | ND | ND | ND | ND |
| N.epsilon.-Dimethyl-L-lysine | ND | ND | ND | 346 | ND | ND | 442 |
| N1,N12-Diethylspermine | ND | ND | ND | 115 | ND | ND | ND |
| N2-(1-Oxo-4-phenylbutyl)-L-glutamine | ND | 14 | ND | ND | ND | ND | ND |
| N2-Methylguanosine | ND | 553 | ND | ND | ND | ND | 441 |
| NAE 16:0 | ND | ND | ND | ND | ND | 429 | ND |
| NAE 18:0 | ND | ND | ND | 382 | ND | 430 | ND |
| NAE 18:1 | ND | ND | ND | 269 | ND | 431 | ND |
| NAE 18:3 | ND | ND | ND | ND | ND | 345 | ND |
| NAE 20:4 | ND | ND | ND | ND | ND | 432 | ND |
| NAE 22:5 | ND | ND | ND | 233 | ND | ND | ND |
| NAE 23:1 | ND | ND | ND | ND | ND | 300 | ND |
| NG,NG-Dimethyl-L-arginine | ND | ND | ND | 383 | ND | ND | ND |
| Nicotinamide adenine dinucleotide | ND | ND | ND | ND | ND | ND | 439 |
| Nicotinamide riboside | ND | ND | ND | ND | ND | 433 | ND |
| Nicotinamide riboside cation | ND | ND | ND | ND | 102 | ND | 438 |
| Norcarane-7-carboxylic acid | ND | ND | ND | 357 | ND | ND | ND |
| Norvaline | ND | ND | 363 | ND | ND | ND | ND |
| Nudifloramide | ND | 203 | ND | ND | ND | ND | ND |
| O-Desmethyl-cis-tramadol | ND | ND | 148 | ND | ND | ND | ND |
| O-Desmethylvenlafaxine | ND | ND | 373 | ND | ND | ND | ND |
| O1_CE 18:3 | ND | ND | ND | ND | 242 | ND | ND |
| O1_Cer d34:1 | ND | ND | ND | ND | 106 | ND | 437 |
| O1_Cer d36:1 | ND | 397 | ND | ND | ND | ND | ND |
| O1_Cer d38:1 | ND | 219 | ND | ND | ND | ND | ND |
| O1_Cer d40:1 | ND | 221 | ND | ND | 135 | ND | 169 |
| O1_Cer d41:1 | ND | ND | ND | ND | ND | ND | 475 |
| O1_Cer d42:1 | ND | 267 | ND | ND | 60 | ND | ND |
| O1_Cer d42:2 | ND | 492 | ND | 358 | 243 | ND | 474 |
| O1_Ceramide d34:1 | ND | ND | 139 | ND | ND | ND | 473 |
| O1_Ceramide d40:1 | ND | ND | 205 | ND | ND | ND | ND |
| O1_Ceramide d42:2 | ND | ND | 243 | ND | ND | ND | ND |
| O1_Ceramide d44:1 | ND | ND | ND | 216 | ND | ND | ND |
| O1_FA 14:0 | ND | ND | ND | 167 | ND | ND | ND |
| O1_FA 14:1 | ND | ND | ND | ND | ND | ND | 472 |
| O1_FA 15:1 | ND | ND | ND | ND | 3 | ND | 471 |
| O1_FA 15:4 | ND | ND | ND | ND | ND | 33 | ND |
| O1_FA 16:0 | ND | ND | ND | ND | ND | 409 | ND |
| O1_FA 16:1 | ND | ND | ND | ND | 145 | ND | 215 |
| O1_FA 16:2 | ND | ND | ND | 13 | ND | 434 | ND |
| O1_FA 18:2 | ND | ND | ND | 359 | 331 | 435 | 470 |
| O1_FA 18:3 | ND | ND | ND | 97 | 179 | 436 | 469 |
| O1_FA 18:4 | ND | ND | ND | ND | ND | 437 | ND |

| | | | | | | | |
|---|---|---|---|---|---|---|---|
| O1_FA 20:3 | ND | ND | ND | 150 | 287 | ND | ND |
| O1_FA 20:4 | ND | 45 | ND | 63 | ND | 438 | 468 |
| O1_FA 20:5 | ND | ND | ND | 9 | ND | ND | 467 |
| O1_FA 22:6 | ND | ND | ND | 387 | ND | 439 | ND |
| O1_GlcCer d34:1 | ND | ND | ND | ND | ND | ND | 466 |
| O1_GlcCer d40:1 | ND | ND | ND | 388 | ND | ND | 255 |
| O1_GlcCer d41:1 | ND | ND | ND | ND | 216 | ND | ND |
| O1_GlcCer d42:1 | ND | ND | ND | ND | 575 | ND | ND |
| O1_GlcCer d42:2 | ND | 229 | ND | ND | 372 | 440 | ND |
| O1_LPC 20:1 | ND | ND | ND | ND | ND | ND | 465 |
| O1_LPC 18:1 | ND | ND | ND | ND | 361 | ND | ND |
| O1_LPC 18:2 | ND | ND | ND | ND | 204 | 318 | ND |
| O1_LPC 20:2 | ND | ND | ND | ND | 311 | ND | ND |
| O1_LPC 20:3 | ND | ND | ND | ND | 266 | ND | ND |
| O1_PC 34:2 | ND | 224 | ND | ND | 267 | ND | 464 |
| O1_PC 34:3 | ND | ND | ND | ND | 567 | ND | 463 |
| O1_PC 36:3 | ND | ND | ND | ND | 366 | ND | ND |
| O1_PC 36:4 | ND | 372 | ND | ND | 261 | ND | 462 |
| O1_PC 36:5 | ND | 57 | ND | ND | 330 | 441 | ND |
| O1_PC 36:6 | ND | ND | ND | ND | ND | ND | 489 |
| O1_PC 38:4 | ND | ND | ND | ND | 235 | ND | ND |
| O1_PC 38:6 | ND | ND | ND | ND | 129 | 442 | ND |
| O1_PC 38:7 | ND | ND | ND | ND | ND | 419 | 488 |
| O1_PC 40:5 | ND | ND | ND | ND | 181 | ND | ND |
| O1_PC 40:6 | ND | ND | ND | ND | 160 | ND | ND |
| O1_PC 40:7 | ND | ND | ND | ND | 568 | ND | 283 |
| O1_PE 38:4 | ND | ND | ND | ND | ND | 443 | ND |
| O1_PE 38:4 B | ND | ND | ND | 389 | ND | ND | ND |
| O1_PE 38:6 | ND | ND | ND | ND | 385 | ND | 487 |
| O1_SHexCer 40:1 | ND | ND | ND | 390 | ND | ND | ND |
| O1_SHexCer 40:1;2O | ND | ND | ND | ND | ND | 40 | ND |
| O1_SHexCer 42:1 | ND | ND | ND | 391 | ND | ND | ND |
| O1_SHexCer 42:1;2O | ND | ND | ND | ND | ND | 283 | ND |
| O1_SHexCer 42:2;2O | ND | ND | ND | ND | ND | 444 | ND |
| O1_SM d34:1 | ND | ND | ND | 119 | ND | 128 | ND |
| O1_SM d40:1 | ND | ND | ND | ND | ND | ND | 486 |
| O1_SM d42:1 | ND | ND | ND | ND | ND | ND | 5 |
| O1_TAG 42:1 | ND | ND | ND | ND | 91 | ND | ND |
| O1_TAG 44:1 | ND | ND | ND | ND | 256 | ND | ND |
| O1_TAG 48:4 | ND | ND | ND | 66 | ND | ND | ND |
| O1_TAG 49:1 | ND | ND | ND | ND | ND | ND | 485 |
| O1_TAG 49:2 | ND | ND | ND | ND | 255 | ND | 484 |
| O1_TAG 49:3 | ND | ND | ND | ND | 334 | ND | ND |

| | | | | | | | |
|---|---|---|---|---|---|---|---|
| O1_TAG 50:1 | ND | ND | ND | ND | 569 | ND | 483 |
| O1_TAG 50:2 | ND | ND | ND | ND | 570 | ND | 482 |
| O1_TAG 51:1 | ND | ND | ND | ND | ND | ND | 481 |
| O1_TAG 51:2 | ND | ND | ND | ND | ND | ND | 480 |
| O1_TAG 51:3 | ND | ND | ND | ND | ND | ND | 479 |
| O1_TAG 52:2 | ND | ND | ND | ND | 182 | ND | 478 |
| O1_TAG 52:3 | ND | ND | ND | ND | 559 | ND | 477 |
| O1_TAG 52:4 | ND | ND | ND | 392 | ND | ND | 476 |
| O1_TAG 52:5 | ND | ND | ND | 393 | ND | ND | 503 |
| O1_TAG 53:1 | ND | ND | ND | ND | ND | ND | 502 |
| O1_TAG 53:3 | ND | ND | ND | ND | ND | ND | 501 |
| O1_TAG 53:4 | ND | ND | ND | ND | ND | ND | 500 |
| O1_TAG 53:5 | ND | ND | ND | ND | ND | ND | 499 |
| O1_TAG 54:3 | ND | ND | ND | ND | 271 | ND | ND |
| O1_TAG 54:4 | ND | ND | ND | 394 | 561 | ND | 498 |
| O1_TAG 54:5 | ND | ND | ND | 395 | ND | ND | 497 |
| O1_TAG 55:1 | ND | ND | ND | ND | ND | ND | 496 |
| O1_TAG 55:2 | ND | ND | ND | ND | 252 | ND | ND |
| O1_TAG 55:3 | ND | ND | ND | ND | ND | ND | 495 |
| O1_TAG 56:7 | ND | ND | ND | ND | 562 | ND | ND |
| O1_TAG 57:2 | ND | ND | ND | ND | 563 | ND | ND |
| O1_TG 41:1 | ND | ND | ND | 236 | ND | ND | ND |
| O1_TG 43:2 | ND | ND | ND | 379 | ND | ND | ND |
| O1_TG 43:3 | ND | ND | ND | 217 | ND | ND | ND |
| O1_TG 45:4\|TG 10:0_18:2_17:2;1O | ND | ND | ND | ND | ND | 445 | ND |
| O1_TG 45:5 | ND | ND | ND | 380 | ND | ND | ND |
| O1_TG 50:2 | ND | ND | ND | ND | ND | 72 | ND |
| O1_TG 50:3 | ND | ND | ND | ND | ND | 446 | ND |
| O1_TG 50:4 | ND | ND | ND | 381 | ND | 447 | ND |
| O1_TG 52:3 | ND | ND | ND | 409 | ND | ND | ND |
| O1_TG 52:3\|TG 16:0_18:1_18:2;1O | ND | ND | ND | ND | ND | 180 | ND |
| O1_TG 52:4 | ND | ND | ND | 410 | ND | ND | ND |
| O1_TG 52:4\|TG 16:0_18:1_18:3;1O | ND | ND | ND | ND | ND | 448 | ND |
| O1_TG 52:5 | ND | ND | ND | 411 | ND | ND | ND |
| O1_TG 52:5\|TG 16:0_18:2_18:3;1O | ND | ND | ND | ND | ND | 449 | ND |
| O1_TG 52:6 | ND | ND | ND | 412 | ND | ND | ND |
| O1_TG 52:6\|TG 16:1_18:2_18:3;1O | ND | ND | ND | ND | ND | 137 | ND |
| O1_TG 54:4 | ND | ND | ND | 413 | ND | ND | ND |
| O1_TG 54:4\|TG 18:1_18:2_18:1;1O | ND | ND | ND | ND | ND | 237 | ND |
| O1_TG 54:5 | ND | ND | ND | 11 | ND | ND | ND |
| O1_TG 54:5\|TG 18:1_18:2_18:2;1O | ND | ND | ND | ND | ND | 374 | ND |
| O1_TG 54:6 | ND | ND | ND | 414 | ND | ND | ND |
| O1_TG 54:6\|TG 18:1_18:2_18:3;1O | ND | ND | ND | ND | ND | 450 | ND |

| | | | | | | | | |
|---|---|---|---|---|---|---|---|---|
| O1_free fatty acid 16:0 | ND | ND | 3 | ND | ND | ND | ND | |
| O1_free fatty acid 16:2 | ND | ND | 15 | ND | ND | ND | ND | |
| O1_free fatty acid 18:0 | ND | ND | 2 | ND | ND | ND | ND | |
| O1_free fatty acid 18:2 | ND | ND | 68 | ND | ND | ND | ND | |
| O1_free fatty acid 18:3 | ND | ND | 300 | ND | ND | ND | ND | |
| O1_free fatty acid 20:4 | ND | ND | 25 | ND | ND | ND | ND | |
| O1_free fatty acid 20:5 | ND | ND | 286 | ND | ND | ND | ND | |
| O1_free fatty acid 22:6 | ND | ND | 22 | ND | ND | ND | ND | |
| O1_phosphatidylethanolamine 38:4 | ND | ND | 231 | ND | ND | ND | ND | |
| O1_sphingomyelin d34:1 | ND | ND | 11 | ND | ND | ND | ND | |
| O1_triacylglyceride 41:1 | ND | ND | 30 | ND | ND | ND | ND | |
| O1_triacylglyceride 50:4 | ND | ND | 151 | ND | ND | ND | ND | |
| O1_triacylglyceride 52:3 | ND | ND | 77 | ND | ND | ND | ND | |
| O1_triacylglyceride 52:4 | ND | ND | 323 | ND | ND | ND | ND | |
| O1_triacylglyceride 52:5 | ND | ND | 302 | ND | ND | ND | ND | |
| O1_triacylglyceride 54:5 | ND | ND | 123 | ND | ND | ND | ND | |
| O1_triacylglyceride 54:6 | ND | ND | 84 | ND | ND | ND | ND | |
| O2_Cer d40:1 | ND | ND | ND | 44 | ND | ND | ND | |
| O2_Cer d42:1 | ND | ND | ND | ND | ND | ND | | 305 |
| O2_Cer d42:2 | ND | ND | ND | 234 | ND | ND | ND | |
| O2_Ceramide d38:1 | ND | ND | 281 | ND | ND | ND | ND | |
| O2_Ceramide d40:1 | ND | ND | 159 | ND | ND | ND | ND | |
| O2_Ceramide d42:2 | ND | ND | 276 | ND | ND | ND | ND | |
| O2_FA 16:1 | ND | ND | ND | ND | 247 | ND | ND | |
| O2_FA 18:0 | ND | ND | ND | 17 | ND | ND | ND | |
| O2_FA 18:1 | ND | ND | ND | 384 | ND | ND | ND | |
| O2_FA 18:2 | ND | ND | ND | 109 | ND | 451 | | 494 |
| O2_FA 18:3 | ND | ND | ND | ND | 340 | 452 | ND | |
| O2_FA 18:3 A | ND | ND | ND | 53 | ND | ND | ND | |
| O2_FA 20:5 | ND | ND | ND | ND | 116 | ND | | 493 |
| O2_FA 22:5 | ND | ND | ND | 385 | ND | ND | ND | |
| O2_FA 22:6 | ND | ND | ND | 386 | 238 | 453 | ND | |
| O2_GlcCer d34:1 | ND | ND | ND | ND | ND | ND | | 492 |
| O2_LPC 20:2 | ND | ND | ND | ND | ND | 257 |  | 491 |
| O2_LPC 18:3 | ND | ND | ND | ND | 234 | ND | ND | |
| O2_PC 34:1 | ND | ND | ND | ND | ND | ND | | 300 |
| O2_PC 35:3 | ND | ND | ND | ND | ND | ND | | 490 |
| O2_PC 35:4 | ND | ND | ND | ND | 9 | ND | ND | |
| O2_PC 36:2 | ND | ND | ND | ND | ND | ND | | 515 |
| O2_PC 36:4 | ND | ND | ND | ND | 564 | 454 | | 211 |
| O2_PC 38:4 | ND | ND | ND | ND | 223 | ND | | 514 |
| O2_PC 38:6 | ND | ND | ND | ND | 397 | 455 | ND | |
| O2_PC 40:3 | ND | ND | ND | ND | ND | ND | | 17 |

| | | | | | | | |
|---|---|---|---|---|---|---|---|
| O2_PC 40:6 | ND | ND | ND | ND | 207 | ND | 513 |
| O2_PC 40:7 | ND | ND | ND | ND | 251 | ND | 512 |
| O2_PC 40:8 | ND | ND | ND | ND | ND | ND | 511 |
| O2_PC p-32:1/PC o-32:2 | ND | ND | ND | ND | 565 | ND | ND |
| O2_PC p-34:1/PC o-34:2 | ND | ND | ND | ND | 178 | ND | ND |
| O2_PC p34:1/PC o34:2 | ND | 408 | ND | ND | ND | ND | ND |
| O2_PE 36:4 | ND | ND | ND | ND | 566 | ND | ND |
| O2_PE 38:4 | ND | ND | ND | ND | 303 | ND | ND |
| O2_PE 38:6 | ND | ND | ND | ND | 556 | ND | ND |
| O2_PE 40:6 | ND | ND | ND | ND | 398 | ND | ND |
| O2_PE o-40:5 | ND | ND | ND | 428 | ND | ND | ND |
| O2_PE o-40:6 | ND | ND | ND | 429 | ND | ND | ND |
| O2_PE o-40:8 | ND | ND | ND | 103 | ND | ND | ND |
| O2_SM d34:1 | ND | ND | ND | ND | ND | 456 | ND |
| O2_TAG 54:2 | ND | ND | ND | ND | ND | ND | 510 |
| O2_TG 52:3 | ND | ND | ND | 399 | ND | ND | ND |
| O2_TG 52:5 | ND | ND | ND | 400 | ND | ND | ND |
| O2_TG 54:6 | ND | ND | ND | 82 | ND | ND | ND |
| O2_free fatty acid 18:0 | ND | ND | 17 | ND | ND | ND | ND |
| O2_free fatty acid 18:1 | ND | ND | 321 | ND | ND | ND | ND |
| O2_free fatty acid 18:2 | ND | ND | 295 | ND | ND | ND | ND |
| O2_free fatty acid 18:3 A | ND | ND | 67 | ND | ND | ND | ND |
| O2_free fatty acid 18:3 B | ND | ND | 336 | ND | ND | ND | ND |
| O2_free fatty acid 20:4 | ND | ND | 249 | ND | ND | ND | ND |
| O2_free fatty acid 22:6 | ND | ND | 263 | ND | ND | ND | ND |
| O2_phosphatidylcholine 38:5 | ND | ND | 280 | ND | ND | ND | ND |
| O2_phosphatidylethanolamine 38:2 | ND | ND | 72 | ND | ND | ND | ND |
| O2_triacylglyceride 50:3 | ND | ND | 210 | ND | ND | ND | ND |
| O2_triacylglyceride 52:3 | ND | ND | 229 | ND | ND | ND | ND |
| O2_triacylglyceride 52:4 | ND | ND | 202 | ND | ND | ND | ND |
| O2_triacylglyceride 54:4 | ND | ND | 303 | ND | ND | ND | ND |
| O2_triacylglyceride 54:5 | ND | ND | 32 | ND | ND | ND | ND |
| O2_triacylglyceride 54:6 | ND | ND | 6 | ND | ND | ND | ND |
| O3_FA 18:1 | ND | ND | ND | 212 | ND | ND | ND |
| O3_FA 18:3 | ND | ND | ND | ND | 195 | ND | 64 |
| O3_FA 20:5 | ND | ND | ND | ND | 199 | ND | ND |
| O3_LPC 20:2 | ND | ND | ND | ND | 270 | 457 | ND |
| O3_LPC 20:1 | ND | ND | ND | ND | 545 | ND | ND |
| O3_LPC 20:5 | ND | ND | ND | ND | 123 | ND | ND |
| O3_PE o-40:7 | ND | ND | ND | 401 | ND | ND | ND |
| O3_TAG 50:4 | ND | ND | ND | ND | ND | ND | 304 |
| O3_free fatty acid 18:1 | ND | ND | 87 | ND | ND | ND | ND |
| O4_FA 20:4 | ND | 270 | ND | ND | ND | ND | ND |

| | | | | | | | |
|---|---|---|---|---|---|---|---|
| O4_PC 38:6 | ND | ND | ND | ND | 341 | ND | ND |
| Oleamide | ND | ND | ND | 79 | ND | ND | ND |
| Oleoyl-L-carnitine | ND | ND | ND | ND | 246 | ND | 508 |
| Oleoyl-carnitine | ND | ND | ND | ND | ND | 392 | ND |
| Ornithine | ND | 262 | 277 | 402 | 191 | 229 | 507 |
| PC 18:1_14:0 | ND | ND | ND | ND | 356 | ND | ND |
| PC 25:0 | ND | ND | ND | ND | 220 | ND | ND |
| PC 26:0 | ND | 332 | ND | ND | ND | ND | ND |
| PC 28:0 | ND | ND | ND | ND | 157 | ND | 505 |
| PC 30:0 | ND | 59 | ND | 403 | 13 | 459 | 504 |
| PC 30:1 \| PC 14:0_16:1 | ND | ND | ND | ND | ND | 460 | ND |
| PC 31:0 | ND | 354 | ND | 404 | 21 | 461 | 525 |
| PC 31:0 \| PC 15:0_16:0 | ND | ND | ND | ND | ND | 462 | ND |
| PC 31:1 | ND | 347 | ND | ND | 390 | 463 | ND |
| PC 32:0 | ND | 263 | ND | 405 | 546 | 464 | 205 |
| PC 32:1 | ND | 438 | ND | 406 | 550 | 225 | 524 |
| PC 32:3 | ND | ND | ND | ND | ND | 282 | 155 |
| PC 33:0 | ND | 164 | ND | ND | 307 | 9 | 11 |
| PC 33:1 | ND | 455 | ND | 408 | 552 | 466 | ND |
| PC 33:1.1 | ND | ND | ND | ND | ND | 467 | ND |
| PC 33:3 | ND | ND | ND | ND | ND | 369 | ND |
| PC 34:1 | ND | 465 | ND | 418 | 209 | 469 | 520 |
| PC 34:2 | ND | 98 | ND | 419 | 532 | 470 | 3 |
| PC 34:3 | ND | 431 | ND | 420 | 525 | 471 | 183 |
| PC 34:3.1 | ND | ND | ND | ND | ND | 472 | ND |
| PC 34:4 | ND | 439 | ND | ND | 393 | 473 | 519 |
| PC 35:0 | ND | ND | ND | ND | ND | 68 | 33 |
| PC 35:1 | ND | 377 | ND | 421 | 214 | 474 | 518 |
| PC 35:2 | ND | 495 | ND | 28 | 526 | 475 | 517 |
| PC 35:2.1 | ND | ND | ND | ND | ND | 476 | ND |
| PC 35:3 | ND | ND | ND | 422 | 527 | 477 | 516 |
| PC 35:4.1 | ND | ND | ND | ND | ND | 478 | ND |
| PC 36:1 | ND | 113 | ND | 423 | ND | 480 | 538 |
| PC 36:3 | ND | 50 | ND | 258 | 530 | 481 | 536 |
| PC 36:3.1 | ND | ND | ND | ND | ND | 482 | 535 |
| PC 36:4 | ND | 6 | ND | 5 | 547 | 483 | 534 |
| PC 36:5 | ND | 259 | ND | 226 | 548 | 295 | 533 |
| PC 36:5.1 | ND | ND | ND | ND | ND | 484 | ND |
| PC 36:6 | ND | 189 | ND | ND | 533 | 398 | 532 |
| PC 36:6 \| PC 14:0_22:6 | ND | ND | ND | ND | ND | 485 | ND |
| PC 36:7 | ND | ND | ND | ND | ND | ND | 531 |
| PC 37:1 | ND | ND | ND | ND | ND | 486 | ND |
| PC 37:3 | ND | ND | ND | ND | 53 | 294 | 529 |

| | | | | | | | |
|---|---|---|---|---|---|---|---|
| PC 37:4 | ND | 150 | ND | ND | 535 | 487 | 528 |
| PC 37:5 | ND | 95 | ND | ND | 125 | 488 | 527 |
| PC 37:6 | ND | 476 | ND | ND | 536 | 489 | 6 |
| PC 37:7 | ND | ND | ND | ND | ND | ND | 526 |
| PC 38:1 | ND | 340 | ND | ND | 510 | 490 | 273 |
| PC 38:3 | ND | 441 | ND | 427 | 511 | 492 | 563 |
| PC 38:3.1 | ND | ND | ND | ND | ND | 493 | ND |
| PC 38:4 | ND | 502 | ND | 40 | 395 | 494 | 63 |
| PC 38:5 | ND | 445 | ND | ND | 512 | 495 | 562 |
| PC 38:5.1 | ND | ND | ND | ND | ND | 496 | ND |
| PC 38:7 \| PC 16:1_22:6 | ND | ND | ND | ND | ND | 499 | ND |
| PC 38:8 | ND | ND | ND | ND | ND | 500 | 560 |
| PC 38:8 \| PC 16:2_22:6 | ND | ND | ND | ND | ND | 501 | ND |
| PC 39:4 | ND | 64 | ND | ND | 514 | 395 | 559 |
| PC 39:5 \| PC 17:0_22:5 | ND | ND | ND | ND | ND | 502 | ND |
| PC 39:7 | ND | ND | ND | ND | ND | ND | 557 |
| PC 39:7 \| PC 17:1_22:6 | ND | ND | ND | ND | ND | 503 | ND |
| PC 39:8 | ND | ND | ND | ND | ND | 60 | ND |
| PC 40:0 | ND | ND | ND | ND | ND | ND | 556 |
| PC 40:11 | ND | ND | ND | ND | ND | ND | 2 |
| PC 40:1 \| PC 22:0_18:1 | ND | ND | ND | ND | ND | 504 | ND |
| PC 40:2 | ND | ND | ND | ND | ND | 505 | ND |
| PC 40:4 | ND | ND | ND | ND | 518 | 506 | 555 |
| PC 40:4.1 | ND | ND | ND | ND | ND | 507 | ND |
| PC 40:5.1 | ND | ND | ND | ND | ND | 509 | ND |
| PC 40:6 | ND | 4 | ND | 440 | 519 | 50 | 553 |
| PC 40:6.1 | ND | ND | ND | ND | ND | 510 | ND |
| PC 40:7 | ND | 305 | ND | 441 | 520 | 511 | 576 |
| PC 40:8 | ND | 375 | ND | 120 | 523 | 14 | 575 |
| PC 40:8.1 | ND | ND | ND | ND | ND | 512 | ND |
| PC 40:9 | ND | ND | ND | ND | ND | ND | 574 |
| PC 40:9 \| PC 18:3_22:6 | ND | ND | ND | ND | ND | 240 | ND |
| PC 41:1 | ND | ND | ND | ND | ND | 513 | ND |
| PC 41:7 | ND | ND | ND | ND | ND | 515 | 572 |
| PC 42:10 | ND | 177 | ND | ND | 350 | 516 | 571 |
| PC 42:10 \| PC 20:4_22:6 | ND | ND | ND | ND | ND | 416 | ND |
| PC 42:11 | ND | ND | ND | ND | ND | 517 | ND |
| PC 42:1 \| PC 24:0_18:1 | ND | ND | ND | 442 | ND | 518 | ND |
| PC 42:5 | ND | ND | ND | ND | 239 | ND | ND |
| PC 42:6 | ND | 398 | ND | ND | 524 | 520 | 302 |
| PC 42:8 | ND | ND | ND | ND | ND | 23 | 570 |
| PC 42:9 | ND | ND | ND | ND | ND | 521 | 569 |
| PC 44:10 | ND | ND | ND | ND | ND | 132 | ND |

| | | | | | | | |
|---|---|---|---|---|---|---|---|
| PC 44:11 | ND | ND | ND | ND | ND | ND | 568 |
| PC 44:11 \| PC 22:5_22:6 | ND | ND | ND | ND | ND | 522 | ND |
| PC 44:12 | ND | ND | ND | ND | ND | ND | 567 |
| PC 44:12 \| PC 22:6_22:6 | ND | ND | ND | ND | ND | 523 | ND |
| PC 44:6 | ND | ND | ND | ND | ND | ND | 566 |
| PC 44:8 | ND | ND | ND | ND | ND | ND | 203 |
| PC 45:11 | ND | ND | ND | ND | ND | ND | 565 |
| PC 46:12 | ND | ND | ND | ND | ND | ND | 250 |
| PC 46:6 | ND | ND | ND | ND | ND | ND | 590 |
| PC 46:7 | ND | ND | ND | ND | ND | ND | 115 |
| PC 60:11 | ND | ND | ND | ND | ND | ND | 589 |
| PC 62:12 | ND | ND | ND | ND | ND | ND | 588 |
| PC 64:12 | ND | ND | ND | ND | ND | ND | 587 |
| PC 64:13 | ND | ND | ND | ND | ND | ND | 586 |
| PC 64:15 | ND | ND | ND | ND | ND | ND | 585 |
| PC 64:17 | ND | ND | ND | ND | ND | ND | 584 |
| PC 66:17 | ND | ND | ND | ND | ND | ND | 583 |
| PC O-30:0 | ND | ND | ND | ND | ND | 524 | ND |
| PC O-32:1 | ND | ND | ND | ND | ND | 526 | ND |
| PC O-32:1.1 | ND | ND | ND | ND | ND | 527 | ND |
| PC O-33:6 | ND | ND | ND | ND | ND | 528 | ND |
| PC O-34:0 | ND | ND | ND | ND | ND | 529 | ND |
| PC O-34:1.1 | ND | ND | ND | ND | ND | 531 | ND |
| PC O-34:2.1 | ND | ND | ND | ND | ND | 195 | ND |
| PC O-34:3.1 | ND | ND | ND | ND | ND | 534 | ND |
| PC O-36:5.1 | ND | ND | ND | ND | ND | 537 | ND |
| PC O-36:7 | ND | ND | ND | ND | ND | 538 | ND |
| PC O-37:9 | ND | ND | ND | ND | ND | 168 | ND |
| PC O-38:5 | ND | ND | ND | ND | ND | 541 | ND |
| PC O-38:5.1 | ND | ND | ND | ND | ND | 542 | ND |
| PC O-38:6.1 | ND | ND | ND | ND | ND | 952 | ND |
| PC O-38:7 \| PC O-16:1_22:6 | ND | ND | ND | ND | ND | 546 | ND |
| PC O-39:7 | ND | ND | ND | ND | ND | 548 | ND |
| PC O-40:5 | ND | ND | ND | ND | ND | 549 | ND |
| PC O-40:8 \| PC O-18:2_22:6 | ND | ND | ND | ND | ND | 84 | ND |
| PC O-40:9 | ND | ND | ND | ND | ND | 554 | ND |
| PC o-32:0 | ND | ND | ND | 443 | ND | ND | ND |
| PC o-32:1 | ND | ND | ND | 58 | ND | ND | ND |
| PC o-34:1 | ND | ND | ND | 444 | ND | ND | ND |
| PC o-34:2 | ND | ND | ND | 38 | ND | ND | ND |
| PC o-36:4 | ND | ND | ND | 188 | ND | ND | ND |
| PC o-36:5 | ND | ND | ND | 204 | ND | ND | ND |
| PC o-37:2 \| PC o-19:0_18:2 | ND | ND | ND | 415 | ND | ND | ND |

| | | | | | | | |
|---|---|---|---|---|---|---|---|
| PC o-38:5 | ND | ND | ND | 416 | ND | ND | ND |
| PC o-38:6 | ND | ND | ND | 461 | ND | ND | ND |
| PC p-36:1/PC o-36:2 | ND | ND | ND | ND | 500 | ND | ND |
| PC p-42:4/PC o-42:5 | ND | ND | ND | ND | 367 | ND | ND |
| PC p38:2 | ND | 331 | ND | ND | ND | ND | ND |
| PC p42:3 | ND | 368 | ND | ND | ND | ND | ND |
| PE 32:0 \| PE 16:0_16:0 | ND | ND | ND | ND | ND | 172 | ND |
| PE 32:1 | ND | ND | ND | ND | ND | ND | 582 |
| PE 32:1 \| PE 16:0_16:1 | ND | ND | ND | ND | ND | 25 | ND |
| PE 34:0 | ND | ND | ND | ND | ND | 557 | ND |
| PE 34:0.1 | ND | ND | ND | ND | ND | 558 | ND |
| PE 34:1 | ND | ND | ND | ND | ND | 559 | ND |
| PE 34:3 | ND | ND | ND | ND | ND | ND | 581 |
| PE 34:3 \| PE 16:1_18:2 | ND | ND | ND | 463 | ND | 376 | ND |
| PE 35:1 | ND | ND | ND | ND | ND | ND | 580 |
| PE 35:2 | ND | ND | ND | ND | ND | ND | 579 |
| PE 35:2 \| PE 17:0_18:2 | ND | ND | ND | 160 | ND | ND | ND |
| PE 36:1 | ND | 206 | ND | ND | 494 | 563 | 578 |
| PE 36:2 | ND | 63 | ND | ND | 495 | 8 | 577 |
| PE 36:2.1 | ND | ND | ND | ND | ND | 16 | ND |
| PE 36:3 | ND | 86 | ND | ND | 344 | 567 | 106 |
| PE 36:3 \| PE 18:1_18:2 | ND | ND | ND | 466 | ND | 568 | ND |
| PE 36:4 | ND | 193 | ND | ND | 496 | 569 | 599 |
| PE 36:4.1 | ND | ND | ND | ND | ND | 570 | ND |
| PE 36:5 \| PE 16:0_20:5 | ND | ND | ND | 256 | ND | 571 | ND |
| PE 36:5 \| PE 16:1_20:4 | ND | ND | ND | 271 | ND | ND | ND |
| PE 36:6 | ND | ND | ND | ND | ND | 171 | ND |
| PE 36:6 \| PE 14:0_22:6 | ND | ND | ND | ND | ND | 573 | ND |
| PE 37:4 | ND | ND | ND | ND | ND | ND | 158 |
| PE 37:4 \| PE 17:0_20:4 | ND | ND | ND | ND | ND | 574 | ND |
| PE 38:1 | ND | ND | ND | ND | ND | ND | 598 |
| PE 38:2 | ND | ND | ND | ND | ND | ND | 114 |
| PE 38:3 \| PE 18:0_20:3 | ND | ND | ND | 467 | ND | 575 | ND |
| PE 38:4 | ND | 399 | ND | ND | 497 | 576 | 160 |
| PE 38:5 | ND | ND | ND | ND | ND | ND | 597 |
| PE 38:5 \| PE 16:0_22:5 | ND | ND | ND | ND | ND | 577 | ND |
| PE 38:5 \| PE 18:0_20:5 | ND | ND | ND | 445 | ND | ND | ND |
| PE 38:6 | ND | 247 | ND | ND | 277 | 578 | 596 |
| PE 38:6.1 | ND | ND | ND | ND | ND | 579 | ND |
| PE 38:7 | ND | ND | ND | ND | ND | ND | 595 |
| PE 38:7 \| PE 16:1_22:6 | ND | ND | ND | 476 | ND | 580 | ND |
| PE 39:4 \| PE 19:0_20:4 | ND | ND | ND | ND | ND | 163 | ND |
| PE 39:6 | ND | ND | ND | ND | ND | ND | 594 |

| | | | | | | | | |
|---|---|---|---|---|---|---|---|---|
| PE 39:6 \| PE 17:0_22:6 | ND | ND | ND | ND | ND | 582 | ND | |
| PE 39:7 | ND | ND | ND | ND | ND | 583 | ND | |
| PE 39:7.1 | ND | ND | ND | ND | ND | 584 | ND | |
| PE 40:4 | ND | ND | ND | ND | ND | 262 | | 116 |
| PE 40:4 \| PE 18:0_22:4 | ND | ND | ND | 477 | ND | ND | ND | |
| PE 40:5 | ND | ND | ND | ND | ND | 586 | | 593 |
| PE 40:5 \| PE 18:0_22:5 | ND | ND | ND | 478 | ND | 587 | ND | |
| PE 40:6 | ND | 161 | ND | ND | 498 | ND | | 592 |
| PE 40:7 | ND | ND | ND | ND | ND | ND | | 591 |
| PE 40:8 | ND | ND | ND | ND | ND | ND | | 290 |
| PE 40:8 \| PE 18:2_22:6 | ND | ND | ND | 481 | ND | 590 | ND | |
| PE 40:9 | ND | ND | ND | ND | ND | 591 | | 141 |
| PE 41:6 | ND | ND | ND | ND | ND | 592 | ND | |
| PE 42:10 | ND | ND | ND | ND | ND | 593 | | 624 |
| PE 42:10 \| PE 20:4_22:6 | ND | ND | ND | ND | ND | 594 | ND | |
| PE 42:11 | ND | ND | ND | ND | ND | 595 | ND | |
| PE 42:1 \| PE 24:0_18:1 | ND | ND | ND | 482 | ND | 596 | ND | |
| PE 42:6 | ND | ND | ND | ND | ND | 597 | | 623 |
| PE 42:6 \| PE 20:0_22:6 | ND | ND | ND | ND | ND | 598 | ND | |
| PE 42:7 \| PE 20:1_22:6 | ND | ND | ND | ND | ND | 108 | ND | |
| PE 42:8 | ND | ND | ND | ND | ND | ND | | 149 |
| PE 42:8 \| PE 20:2_22:6 | ND | ND | ND | ND | ND | 600 | ND | |
| PE 42:9 \| PE 20:3_22:6 | ND | ND | ND | ND | ND | 601 | ND | |
| PE 44:10 \| PE 22:4_22:6 | ND | ND | ND | ND | ND | 602 | ND | |
| PE 44:11 | ND | ND | ND | ND | ND | ND | | 622 |
| PE 44:11 \| PE 22:5_22:6 | ND | ND | ND | ND | ND | 603 | ND | |
| PE 44:12 \| PE 22:6_22:6 | ND | ND | ND | ND | ND | 292 | ND | |
| PE 44:7 | ND | ND | ND | ND | ND | ND | | 256 |
| PE 44:9 \| PE 22:3_22:6 | ND | ND | ND | ND | ND | 309 | ND | |
| PE O-32:2 \| PE O-16:1_16:1 | ND | ND | ND | 483 | ND | ND | ND | |
| PE O-34:1 \| PE O-18:1_16:0 | ND | ND | ND | ND | ND | 606 | ND | |
| PE O-34:2 | ND | ND | ND | ND | ND | 607 | ND | |
| PE O-34:2 \| PE O-16:1_18:1 | ND | ND | ND | 484 | ND | ND | ND | |
| PE O-34:3 | ND | ND | ND | ND | ND | 272 | ND | |
| PE O-35:2 \| PE O-17:1_18:1 | ND | ND | ND | 456 | ND | ND | ND | |
| PE O-36:2 | ND | ND | ND | ND | ND | 609 | ND | |
| PE O-36:2 \| PE O-18:1_18:1 | ND | ND | ND | 457 | ND | ND | ND | |
| PE O-36:3 \| PE O-18:2_18:1 | ND | ND | ND | 458 | ND | 610 | ND | |
| PE O-36:4 \| PE O-18:3_18:1 | ND | ND | ND | 459 | ND | 611 | ND | |
| PE O-36:5 | ND | ND | ND | ND | ND | 250 | ND | |
| PE O-36:6 | ND | ND | ND | ND | ND | 613 | ND | |
| PE O-36:6 \| PE O-16:1_20:5 | ND | ND | ND | 460 | ND | ND | ND | |
| PE O-37:7 \| PE O-15:1_22:6 | ND | ND | ND | ND | ND | 354 | ND | |

| | | | | | | | |
|---|---|---|---|---|---|---|---|
| PE O-38:2 \| PE O-20:1_18:1 | ND | ND | ND | ND | ND | 615 | ND |
| PE O-38:5 \| PE O-16:0_22:5 | ND | ND | ND | ND | ND | 617 | ND |
| PE O-38:5 \| PE O-16:1_22:4 | ND | ND | ND | 30 | ND | 618 | ND |
| PE O-38:6 | ND | ND | ND | ND | ND | 620 | ND |
| PE O-38:7 | ND | ND | ND | ND | ND | 621 | ND |
| PE O-38:7 \| PE O-18:3_20:4 | ND | ND | ND | 26 | ND | ND | ND |
| PE O-38:8 \| PE O-16:2_22:6 | ND | ND | ND | ND | ND | 622 | ND |
| PE O-39:7 \| PE O-17:1_22:6 | ND | ND | ND | 145 | ND | 623 | ND |
| PE O-40:5 \| PE O-18:1_22:4 | ND | ND | ND | ND | ND | 625 | ND |
| PE O-40:5 \| PE O-20:1_20:4 | ND | ND | ND | ND | ND | 31 | ND |
| PE O-40:6 \| PE O-18:1_22:5 | ND | ND | ND | 498 | ND | 628 | ND |
| PE O-40:7 \| PE O-18:2_22:5 | ND | ND | ND | 469 | ND | 630 | ND |
| PE O-40:8 | ND | ND | ND | ND | ND | 631 | ND |
| PE O-40:8 \| PE O-18:2_22:6 | ND | ND | ND | 470 | ND | ND | ND |
| PE O-40:9 \| PE O-18:3_22:6 | ND | ND | ND | 4 | ND | ND | ND |
| PE O-41:7 \| PE O-19:1_22:6 | ND | ND | ND | ND | ND | 632 | ND |
| PE P-34:1 | ND | ND | ND | 472 | ND | 633 | ND |
| PE P-34:2 \| PE P-16:0_18:2 | ND | ND | ND | ND | ND | 634 | ND |
| PE P-35:1 \| PE P-17:0_18:1 | ND | ND | ND | ND | ND | 635 | ND |
| PE P-36:1 | ND | ND | ND | 473 | ND | 636 | ND |
| PE P-36:2 | ND | ND | ND | ND | ND | 637 | ND |
| PE P-36:4 | ND | ND | ND | ND | ND | 638 | ND |
| PE P-38:4 | ND | ND | ND | 474 | ND | ND | ND |
| PE P-38:5 | ND | ND | ND | ND | ND | 639 | ND |
| PE P-40:4 | ND | ND | ND | ND | ND | 351 | ND |
| PE P-40:5 | ND | ND | ND | ND | ND | 642 | ND |
| PE P-40:7 \| PE P-18:1_22:6 | ND | ND | ND | ND | ND | 644 | ND |
| PE P-40:8 \| PE P-18:2_22:6 | ND | ND | ND | ND | ND | 645 | ND |
| PE o-38:6 \| PE O-18:1_20:5 | ND | ND | ND | 475 | ND | ND | ND |
| PE p-34:1/PE o-34:2 | ND | ND | ND | ND | 213 | ND | ND |
| PE p-38:5/PE o-38:6 | ND | ND | ND | ND | 502 | ND | ND |
| PE p-40:4/PE o-40:5 | ND | ND | ND | ND | 501 | ND | ND |
| PE p-40:5/PE o-40:6 | ND | ND | ND | ND | 391 | ND | ND |
| PE p38:3 | ND | 73 | ND | ND | ND | ND | ND |
| PE p40:7 | ND | 275 | ND | ND | ND | ND | ND |
| PG 32:0 \| PG 16:0_16:0 | ND | ND | ND | ND | ND | 289 | ND |
| PG 32:1 \| PG 16:0_16:1 | ND | ND | ND | ND | ND | 223 | ND |
| PG 34:1 \| PG 16:0_18:1 | ND | ND | ND | 506 | ND | 649 | ND |
| PG 34:2 \| PG 16:0_18:2 | ND | ND | ND | 511 | ND | 650 | ND |
| PG 35:1 \| PG 16:0_19:1 | ND | ND | ND | ND | ND | 651 | ND |
| PG 36:1 \| PG 18:0_18:1 | ND | ND | ND | ND | ND | 652 | ND |
| PG 36:2 \| PG 18:0_18:2 | ND | ND | ND | 512 | ND | 653 | ND |
| PG 36:3 \| PG 16:0_20:3 | ND | ND | ND | ND | ND | 654 | ND |

| | | | | | | | |
|---|---|---|---|---|---|---|---|
| PG 36:3 \| PG 18:1_18:2 | ND | ND | ND | 104 | ND | ND | ND |
| PG 36:4 \| PG 16:0_20:4 | ND | ND | ND | ND | ND | 655 | ND |
| PG 36:4 \| PG 18:2_18:2 | ND | ND | ND | 67 | ND | ND | ND |
| PG 38:4 \| PG 18:0_20:4 | ND | ND | ND | ND | ND | 656 | ND |
| PG 38:5 \| PG 16:0_22:5 | ND | ND | ND | ND | ND | 657 | ND |
| PG 38:6 \| PG 16:0_22:6 | ND | ND | ND | ND | ND | 658 | ND |
| PG 40:6 \| PG 18:0_22:6 | ND | ND | ND | ND | ND | 659 | ND |
| PG 40:7 \| PG 18:1_22:6 | ND | ND | ND | ND | ND | 660 | ND |
| PG 40:8 \| PG 18:2_22:6 | ND | ND | ND | ND | ND | 661 | ND |
| PG 44:12 \| PG 22:6_22:6 | ND | ND | ND | ND | ND | 64 | ND |
| PGPC | ND | 261 | ND | ND | 504 | ND | 620 |
| PI 18:0_20:4 | ND | ND | ND | ND | 508 | ND | ND |
| PI 32:0 \| PI 16:0_16:0 | ND | ND | ND | ND | ND | 381 | ND |
| PI 32:1 | ND | ND | ND | ND | ND | 285 | ND |
| PI 34:0 \| PI 16:0_18:0 | ND | ND | ND | ND | ND | 319 | ND |
| PI 34:2 | ND | ND | ND | ND | ND | 666 | ND |
| PI 34:2 \| PI 16:0_18:2 | ND | ND | ND | 513 | ND | ND | ND |
| PI 36:1 | ND | ND | ND | ND | ND | 667 | ND |
| PI 36:1 \| PI 18:0_18:1 | ND | ND | ND | 14 | ND | ND | ND |
| PI 36:2.1 | ND | ND | ND | ND | ND | 669 | ND |
| PI 36:2 \| PI 18:0_18:2 | ND | ND | ND | 164 | ND | ND | ND |
| PI 36:4.1 | ND | ND | ND | ND | ND | 672 | ND |
| PI 36:4 \| PI 16:0_20:4 | ND | ND | ND | 490 | ND | ND | ND |
| PI 38:3 | ND | ND | ND | ND | ND | 675 | 618 |
| PI 38:4 | ND | ND | ND | ND | ND | 676 | 617 |
| PI 38:4.1 | ND | ND | ND | ND | ND | 677 | ND |
| PI 38:5 | ND | ND | ND | ND | ND | 678 | 308 |
| PI 38:5.1 | ND | ND | ND | ND | ND | 679 | ND |
| PI 38:6 | ND | ND | ND | ND | ND | 680 | 178 |
| PI 39:6 \| PI 17:0_22:6 | ND | ND | ND | ND | ND | 683 | ND |
| PI 40:4 \| PI 18:0_22:4 | ND | ND | ND | ND | ND | 684 | ND |
| PI 40:6.1 | ND | ND | ND | ND | ND | 687 | ND |
| PI 40:7 | ND | ND | ND | ND | ND | ND | 269 |
| PI 40:7 \| PI 18:1_22:6 | ND | ND | ND | ND | ND | 688 | ND |
| PI 40:8 \| PI 18:2_22:6 | ND | ND | ND | ND | ND | 689 | ND |
| POVPC | ND | 344 | ND | ND | 422 | ND | 268 |
| PS 18:0_18:2 | ND | ND | ND | ND | 377 | ND | ND |
| PS 36:1 \| PS 18:0_18:1 | ND | ND | ND | 527 | ND | 690 | ND |
| PS 36:2 \| PS 18:0_18:2 | ND | ND | ND | 528 | ND | ND | ND |
| PS 36:2 \| PS 18:1_18:1 | ND | ND | ND | ND | ND | 691 | ND |
| PS 38:4 \| PS 18:0_20:4 | ND | ND | ND | 529 | ND | 692 | ND |
| PS 38:5 \| PS 18:1_20:4 | ND | ND | ND | 530 | ND | ND | ND |
| PS 38:6 | ND | ND | ND | ND | ND | 144 | ND |

| | | | | | | | |
|---|---|---|---|---|---|---|---|
| PS 38:6 \| PS 16:0_22:6 | ND | ND | ND | ND | ND | 694 | ND |
| PS 39:6 \| PS 17:0_22:6 | ND | ND | ND | ND | ND | 695 | ND |
| PS 40:1 \| PS 22:0_18:1 | ND | ND | ND | ND | ND | 696 | ND |
| PS 40:5 \| PS 18:0_22:5 | ND | ND | ND | 531 | ND | 697 | ND |
| PS 40:6 \| PS 18:0_22:6 | ND | ND | ND | 196 | ND | 698 | ND |
| PS 40:7 \| PS 18:1_22:6 | ND | ND | ND | ND | ND | 699 | ND |
| PS 40:8 \| PS 18:2_22:6 | ND | ND | ND | ND | ND | 700 | ND |
| PS 42:1 \| PS 24:0_18:1 | ND | ND | ND | ND | ND | 701 | ND |
| PS 44:11 | ND | ND | ND | ND | ND | 702 | ND |
| PS 44:11 \| PS 22:5_22:6 | ND | ND | ND | ND | ND | 703 | ND |
| PS 44:12 \| PS 22:6_22:6 | ND | ND | ND | ND | ND | 178 | ND |
| Palmitoleoyl ethanolamide | ND | 453 | ND | ND | ND | ND | ND |
| Palmitoyleicosapentaenoyl phosphatidylcholine | ND | ND | ND | ND | ND | 258 | ND |
| Pantetheine | ND | ND | ND | ND | ND | 707 | ND |
| Pantethine | ND | ND | ND | ND | ND | 708 | ND |
| Phe-Ala | ND | ND | ND | ND | ND | ND | 612 |
| Phe-Arg | ND | ND | ND | ND | ND | 30 | 611 |
| Phe-Lys | ND | ND | ND | ND | ND | ND | 640 |
| Phenacyltriphenylphosphonium | ND | ND | ND | 231 | ND | ND | ND |
| Phenethylamine | ND | 25 | ND | ND | ND | ND | ND |
| Phenylacetaldehyde B | ND | 118 | ND | ND | ND | ND | ND |
| Phenylalanine | ND | ND | 389 | 500 | ND | ND | ND |
| Phenylalanine methyl ester | ND | ND | ND | ND | 371 | ND | ND |
| Phomalone | ND | ND | ND | 64 | ND | ND | ND |
| Phosphotyrosine | ND | 428 | ND | ND | ND | ND | ND |
| Piperine | ND | 51 | ND | ND | ND | ND | ND |
| Pregabalin | ND | 21 | ND | ND | ND | ND | ND |
| Pro-Ala | ND | 546 | ND | ND | ND | ND | ND |
| Pro-Arg | ND | 540 | ND | ND | ND | ND | ND |
| Pro-Asn | ND | 545 | ND | ND | ND | ND | 639 |
| Pro-Asp | ND | 547 | ND | ND | ND | ND | ND |
| Pro-Glu | ND | 544 | ND | ND | ND | ND | ND |
| Pro-Leu-Lys | ND | ND | ND | ND | ND | 715 | ND |
| Pro-Lys | ND | 535 | ND | ND | ND | ND | 637 |
| Pro-Pro | ND | 509 | ND | ND | ND | ND | 636 |
| Pro-Pro_b | ND | ND | ND | ND | ND | 301 | ND |
| Pro-Ser | ND | 537 | ND | ND | ND | 302 | 635 |
| Pro-Ser-Arg | ND | ND | ND | ND | 42 | ND | 634 |
| Pro-Thr | ND | 536 | ND | ND | 425 | 326 | 633 |
| Propamocarb | ND | ND | ND | 502 | ND | ND | ND |
| Propanoic acid, 3-[[[2-[(aminoiminomethyl)amino]-4-thiazolyl]methyl]thio]- | ND | ND | 372 | 503 | ND | ND | ND |
| Prophosphatidylinositolonylcarnitine | ND | ND | 311 | ND | ND | ND | ND |
| Protoporphyrin IX | ND | ND | ND | ND | 389 | ND | ND |

| | | | | | | | |
|---|---|---|---|---|---|---|---|
| Pterine | ND | ND | ND | ND | 328 | ND | 630 |
| Purine | ND | 538 | ND | ND | 415 | ND | 629 |
| Pyridine | ND | ND | 306 | ND | ND | 720 | ND |
| Pyridoxine | ND | ND | ND | ND | ND | ND | 626 |
| PyroGlu-Asn-Lys | ND | ND | ND | ND | ND | ND | 625 |
| PyroGlu-Glu-Lys | ND | ND | ND | ND | ND | ND | 661 |
| PyroGlu-Gly-Arg | ND | ND | ND | ND | ND | ND | 652 |
| PyroGlu-Gly-Lys | ND | ND | ND | ND | ND | ND | 651 |
| S-Adenosyl-methionine | ND | 524 | ND | ND | 148 | 723 | 265 |
| SHexCer 36:1;2O | ND | ND | ND | 535 | ND | 724 | ND |
| SHexCer 38:1;2O | ND | ND | ND | 536 | ND | 725 | ND |
| SHexCer 40:1;2O | ND | ND | ND | 537 | ND | 726 | ND |
| SHexCer 41:1;2O | ND | ND | ND | 538 | ND | 727 | ND |
| SHexCer 42:0;2O | ND | ND | ND | 539 | ND | 728 | ND |
| SHexCer 42:1;2O | ND | ND | ND | 540 | ND | 729 | ND |
| SHexCer 42:2;2O | ND | ND | ND | 541 | ND | 730 | ND |
| SHexCer 42:3;2O | ND | ND | ND | 542 | ND | 731 | ND |
| SM d32:1 | ND | 503 | ND | ND | 28 | 352 | ND |
| SM d32:2 | ND | ND | ND | ND | ND | ND | 173 |
| SM d33:1 | ND | 17 | ND | 130 | 7 | 104 | ND |
| SM d34:0 | ND | 80 | ND | 543 | 347 | 734 | 646 |
| SM d34:1 | ND | 292 | ND | 544 | 450 | 735 | 645 |
| SM d34:2 | ND | 452 | ND | 251 | 345 | 736 | 644 |
| SM d35:1 | ND | ND | ND | ND | ND | 737 | ND |
| SM d36:0 | ND | 97 | ND | 546 | 5 | 375 | 222 |
| SM d36:1 | ND | 172 | ND | 525 | 283 | 739 | 19 |
| SM d36:2 | ND | 142 | ND | 553 | 96 | 740 | ND |
| SM d37:1 | ND | 29 | ND | 107 | 451 | 741 | 643 |
| SM d38:0 | ND | 12 | ND | ND | ND | 393 | ND |
| SM d38:1 | ND | 202 | ND | ND | 295 | 743 | 642 |
| SM d38:1 \| SM 14:1;2O/24:0 | ND | ND | ND | 554 | ND | ND | ND |
| SM d38:2 | ND | 395 | ND | 163 | 452 | 276 | 641 |
| SM d39:1 | ND | 37 | ND | 87 | 152 | 745 | 672 |
| SM d39:2 | ND | ND | ND | ND | ND | ND | 105 |
| SM d40:0 | ND | 440 | ND | ND | 396 | 7 | 270 |
| SM d40:1 | ND | 199 | ND | 555 | 421 | 747 | 180 |
| SM d40:2 | ND | 122 | ND | 556 | 320 | 748 | 670 |
| SM d41:1 | ND | 302 | ND | ND | 405 | 34 | 212 |
| SM d41:1 \| SM 18:1;2O/23:0 | ND | ND | ND | 557 | ND | ND | ND |
| SM d41:2 | ND | ND | ND | 558 | 406 | 160 | 669 |
| SM d42:0 | ND | ND | ND | ND | 408 | 751 | 285 |
| SM d42:1 | ND | 430 | ND | ND | 407 | 752 | 152 |
| SM d42:1 \| SM 18:1;2O/24:0 | ND | ND | ND | 559 | ND | ND | ND |

| | | | | | | | |
|---|---|---|---|---|---|---|---|
| SM d42:2 | ND | ND | ND | ND | 412 | 753 | 14 |
| SM d42:2\|SM 18:1;2O/24:1 | ND | ND | ND | 560 | ND | ND | ND |
| SM d42:3 | ND | 393 | ND | 561 | 363 | 754 | 249 |
| SM d43:1 | ND | 429 | ND | ND | 409 | 71 | 666 |
| SM d43:2 | ND | 542 | ND | ND | 410 | ND | 665 |
| SM d44:2 | ND | 463 | ND | ND | ND | ND | ND |
| Saccharopine | ND | 194 | ND | ND | ND | ND | ND |
| Ser-Ala | ND | ND | ND | ND | ND | ND | 664 |
| Ser-Arg | ND | ND | ND | ND | 346 | 757 | 663 |
| Ser-Asn | ND | ND | ND | ND | ND | ND | 662 |
| Ser-Gly | ND | ND | ND | ND | ND | ND | 687 |
| Ser-Lys | ND | ND | ND | ND | ND | 758 | 685 |
| Ser-Pro | ND | ND | ND | ND | ND | ND | 684 |
| Ser-Ser | ND | ND | ND | ND | ND | ND | 683 |
| Ser-Thr | ND | ND | ND | ND | ND | ND | 682 |
| Spermidine | ND | ND | ND | ND | ND | 760 | ND |
| Splitomicin | ND | ND | 366 | ND | ND | ND | ND |
| Stachydrine | ND | ND | ND | ND | ND | 761 | ND |
| Stearoyl-L-carnitine | ND | ND | ND | ND | 221 | 762 | 680 |
| Styrene | ND | ND | 376 | ND | ND | ND | ND |
| Sucrose | ND | ND | 238 | 563 | ND | ND | 293 |
| TAG 40:0 | ND | 315 | ND | ND | 286 | ND | 678 |
| TAG 40:1 | ND | 349 | ND | ND | 262 | ND | 677 |
| TAG 42:0 | ND | 249 | ND | ND | ND | ND | 676 |
| TAG 42:1 | ND | ND | ND | ND | 411 | ND | 675 |
| TAG 42:2 | ND | ND | ND | ND | 349 | ND | 674 |
| TAG 42:3 | ND | ND | ND | ND | ND | ND | 117 |
| TAG 44:0 | ND | 459 | ND | ND | 278 | ND | 673 |
| TAG 44:1 | ND | ND | ND | ND | 378 | ND | 702 |
| TAG 44:2 | ND | ND | ND | ND | ND | ND | 231 |
| TAG 46:0 | ND | 201 | ND | ND | 233 | ND | 701 |
| TAG 46:1 | ND | 436 | ND | ND | 368 | ND | 700 |
| TAG 46:2 | ND | 214 | ND | ND | 231 | ND | 699 |
| TAG 46:3 | ND | ND | ND | ND | 383 | ND | 698 |
| TAG 46:4 | ND | ND | ND | ND | ND | ND | 697 |
| TAG 46:5 | ND | ND | ND | ND | ND | ND | 696 |
| TAG 48:0 | ND | 264 | ND | ND | 282 | ND | 695 |
| TAG 48:1 | ND | 485 | ND | ND | 428 | ND | 260 |
| TAG 48:2 | ND | 343 | ND | ND | 429 | ND | 693 |
| TAG 48:3 | ND | ND | ND | ND | 430 | ND | 692 |
| TAG 48:4 | ND | ND | ND | ND | 431 | ND | 691 |
| TAG 49:0 | ND | 378 | ND | ND | 432 | ND | 690 |
| TAG 49:1 | ND | 254 | ND | ND | 433 | ND | 689 |

| | | | | | | | |
|---|---|---|---|---|---|---|---|
| TAG 49:2 | ND | 433 | ND | ND | 434 | ND | 688 |
| TAG 49:3 | ND | 308 | ND | ND | 435 | ND | 717 |
| TAG 50:0 | ND | ND | ND | ND | 301 | ND | 716 |
| TAG 50:1 | ND | 488 | ND | ND | ND | ND | 715 |
| TAG 50:2 | ND | 279 | ND | ND | 111 | ND | 714 |
| TAG 50:3 | ND | 469 | ND | ND | 436 | ND | ND |
| TAG 50:4 | ND | 296 | ND | ND | 437 | ND | 713 |
| TAG 50:5 | ND | ND | ND | ND | 438 | ND | 712 |
| TAG 51:1 | ND | 481 | ND | ND | 439 | ND | 711 |
| TAG 51:2 | ND | 127 | ND | ND | 357 | ND | 710 |
| TAG 51:3 | ND | 396 | ND | ND | 440 | ND | 709 |
| TAG 51:4 | ND | ND | ND | ND | 441 | ND | 708 |
| TAG 51:5 | ND | ND | ND | ND | ND | ND | 707 |
| TAG 52:0 | ND | 407 | ND | ND | 442 | ND | 206 |
| TAG 52:1 | ND | 552 | ND | ND | 443 | ND | ND |
| TAG 52:2 | ND | 559 | ND | ND | ND | ND | 706 |
| TAG 52:3 | ND | 491 | ND | ND | ND | ND | ND |
| TAG 52:4 | ND | 128 | ND | ND | 444 | ND | ND |
| TAG 52:5 | ND | ND | ND | ND | 445 | ND | 705 |
| TAG 52:6 | ND | ND | ND | ND | 446 | ND | 704 |
| TAG 53:0 | ND | 222 | ND | ND | ND | ND | ND |
| TAG 53:1 | ND | 471 | ND | ND | 447 | ND | 703 |
| TAG 53:2 | ND | ND | ND | ND | 448 | ND | 732 |
| TAG 53:3 | ND | ND | ND | ND | 456 | ND | 731 |
| TAG 53:4 | ND | ND | ND | ND | 455 | ND | 730 |
| TAG 53:5 | ND | ND | ND | ND | 454 | ND | 729 |
| TAG 54:0 | ND | 324 | ND | ND | ND | ND | 728 |
| TAG 54:1 | ND | 557 | ND | ND | 337 | ND | 727 |
| TAG 54:2 | ND | 558 | ND | ND | 453 | ND | ND |
| TAG 54:3 | ND | 271 | ND | ND | ND | ND | ND |
| TAG 54:4 | ND | 286 | ND | ND | 459 | ND | 150 |
| TAG 54:5 | ND | 382 | ND | ND | 460 | ND | 726 |
| TAG 54:6 | ND | 448 | ND | ND | 467 | ND | 725 |
| TAG 54:7 | ND | 414 | ND | ND | ND | ND | 724 |
| TAG 54:8 | ND | ND | ND | ND | 468 | ND | 723 |
| TAG 55:1 | ND | 505 | ND | ND | 457 | ND | 722 |
| TAG 55:2 | ND | 60 | ND | ND | 458 | ND | 721 |
| TAG 55:3 | ND | ND | ND | ND | 483 | ND | 720 |
| TAG 56:0 | ND | 391 | ND | ND | ND | ND | ND |
| TAG 56:1 | ND | 518 | ND | ND | 462 | ND | 719 |
| TAG 56:10 | ND | ND | ND | ND | 463 | ND | 718 |
| TAG 56:2 | ND | 554 | ND | ND | 464 | ND | 747 |
| TAG 56:3 | ND | 390 | ND | ND | 465 | ND | 746 |

| | | | | | | | |
|---|---|---|---|---|---|---|---|
| TAG 56:4 | ND | 472 | ND | ND | 1 | ND | 126 |
| TAG 56:5 | ND | ND | ND | ND | 466 | ND | 745 |
| TAG 56:6 | ND | 195 | ND | ND | 515 | ND | 744 |
| TAG 56:7 | ND | 28 | ND | ND | 516 | ND | 743 |
| TAG 56:8 | ND | ND | ND | ND | 469 | ND | 742 |
| TAG 56:9 | ND | ND | ND | ND | 470 | ND | 741 |
| TAG 57:1 | ND | 227 | ND | ND | ND | ND | 740 |
| TAG 57:2 | ND | 131 | ND | ND | 471 | ND | 739 |
| TAG 58:0 | ND | ND | ND | ND | 164 | ND | ND |
| TAG 58:1 | ND | 501 | ND | ND | 472 | ND | 738 |
| TAG 58:10 | ND | ND | ND | ND | 473 | ND | 737 |
| TAG 58:2 | ND | 555 | ND | ND | 474 | ND | 736 |
| TAG 58:3 | ND | 470 | ND | ND | 475 | ND | 735 |
| TAG 58:4 | ND | ND | ND | ND | 476 | ND | 734 |
| TAG 58:5 | ND | 294 | ND | ND | 477 | ND | 733 |
| TAG 58:6 | ND | 447 | ND | ND | 478 | ND | 761 |
| TAG 58:8 | ND | 173 | ND | ND | 479 | ND | 760 |
| TAG 58:9 | ND | 489 | ND | ND | 480 | ND | 759 |
| TAG 59:2 | ND | 383 | ND | ND | 481 | ND | 758 |
| TAG 59:3 | ND | ND | ND | ND | ND | ND | 757 |
| TAG 60:1 | ND | 373 | ND | ND | 482 | ND | 52 |
| TAG 60:11 | ND | 162 | ND | ND | 486 | ND | ND |
| TAG 60:2 | ND | 478 | ND | ND | 485 | ND | 756 |
| TAG 60:3 | ND | 54 | ND | ND | 487 | ND | 755 |
| TAG 60:4 | ND | ND | ND | ND | 488 | ND | 754 |
| TAG 60:5 | ND | ND | ND | ND | ND | ND | 753 |
| TAG 60:6 | ND | ND | ND | ND | 491 | ND | 752 |
| TAG 62:1 | ND | ND | ND | ND | ND | ND | 12 |
| TAG 62:2 | ND | 295 | ND | ND | 492 | ND | 751 |
| TAG 62:3 | ND | 205 | ND | ND | ND | ND | 750 |
| TAG 62:4 | ND | ND | ND | ND | 376 | ND | 749 |
| TAG 64:2 | ND | ND | ND | ND | 489 | ND | 748 |
| TAG 64:3 | ND | ND | ND | ND | 490 | ND | 777 |
| TAG 64:4 | ND | 110 | ND | ND | 499 | ND | 776 |
| TG 36:0 | ND | ND | ND | 532 | ND | ND | ND |
| TG 38:0 \| TG 10:0_12:0_16:0 | ND | ND | ND | 1 | ND | ND | ND |
| TG 38:1 \| TG 10:0_10:0_18:1 | ND | ND | ND | 571 | ND | ND | ND |
| TG 40:0 | ND | ND | ND | ND | ND | 763 | ND |
| TG 40:0 \| TG 10:0_14:0_16:0 | ND | ND | ND | 572 | ND | ND | ND |
| TG 40:1 | ND | ND | ND | ND | ND | 12 | ND |
| TG 41:2 \| TG 12:0_14:0_15:2 | ND | ND | ND | 573 | ND | ND | ND |
| TG 41:4 \| TG 11:0_15:2_15:2 | ND | ND | ND | 574 | ND | ND | ND |
| TG 42:0 \| TG 12:0_14:0_16:0 | ND | ND | ND | 575 | ND | ND | ND |

| | | | | | | | |
|---|---|---|---|---|---|---|---|
| TG 42:1 | ND | ND | ND | ND | ND | 140 | ND |
| TG 42:2 | ND | ND | ND | ND | ND | 35 | ND |
| TG 42:2 \| TG 12:0_12:0_18:2 | ND | ND | ND | 576 | ND | ND | ND |
| TG 42:3 | ND | ND | ND | ND | ND | 767 | ND |
| TG 42:3 \| TG 10:0_14:0_18:3 | ND | ND | ND | 577 | ND | ND | ND |
| TG 43:1 \| TG 9:0_16:0_18:1 | ND | ND | ND | 578 | ND | ND | ND |
| TG 43:2 \| TG 9:0_16:0_18:2 | ND | ND | ND | ND | ND | 768 | ND |
| TG 43:3 \| TG 12:0_16:1_15:2 | ND | ND | ND | 579 | ND | 769 | ND |
| TG 44:0 | ND | ND | ND | ND | ND | 770 | ND |
| TG 44:0 \| TG 14:0_14:0_16:0 | ND | ND | ND | 548 | ND | ND | ND |
| TG 44:1 | ND | ND | ND | 549 | ND | 75 | ND |
| TG 44:2 | ND | ND | ND | ND | ND | 155 | ND |
| TG 44:2 \| TG 10:0_16:0_18:2 | ND | ND | ND | 550 | ND | ND | ND |
| TG 44:3 | ND | ND | ND | ND | ND | 367 | ND |
| TG 44:3 \| TG 10:0_16:0_18:3 | ND | ND | ND | ND | ND | ND | 775 |
| TG 44:3 \| TG 8:0_18:1_18:2 | ND | ND | ND | 551 | ND | ND | ND |
| TG 44:4 \| TG 8:0_18:2_18:2 | ND | ND | ND | 552 | ND | 774 | 774 |
| TG 45:1 \| TG 12:0_15:0_18:1 | ND | ND | ND | 117 | ND | ND | ND |
| TG 45:1 \| TG 14:0_15:0_16:1 | ND | ND | ND | ND | ND | ND | 772 |
| TG 45:2 \| TG 12:0_15:0_18:2 | ND | ND | ND | 586 | ND | ND | ND |
| TG 45:2 \| TG 15:0_14:1_16:1 | ND | ND | ND | ND | ND | ND | 771 |
| TG 45:3 \| TG 9:0_18:1_18:2 | ND | ND | ND | 587 | ND | 775 | ND |
| TG 45:4 \| TG 9:0_18:2_18:2 | ND | ND | ND | 588 | ND | 331 | ND |
| TG 46:0 | ND | ND | ND | ND | ND | 385 | ND |
| TG 46:1 | ND | ND | ND | ND | ND | 320 | ND |
| TG 46:1 \| TG 14:0_16:0_16:1 | ND | ND | ND | 589 | ND | ND | ND |
| TG 46:2 | ND | ND | ND | 590 | ND | 779 | ND |
| TG 46:3 | ND | ND | ND | 591 | ND | 158 | ND |
| TG 46:4 | ND | ND | ND | ND | ND | 275 | ND |
| TG 46:4 \| TG 10:0_18:2_18:2 | ND | ND | ND | 592 | ND | ND | ND |
| TG 46:5 | ND | ND | ND | ND | ND | 55 | ND |
| TG 46:5 \| TG 10:0_18:2_18:3 | ND | ND | ND | 593 | ND | ND | ND |
| TG 47:0 \| TG 14:0_16:0_17:0 | ND | ND | ND | ND | ND | ND | 770 |
| TG 47:1 | ND | ND | ND | 49 | ND | ND | ND |
| TG 47:1 \| TG 15:0_16:0_16:1 | ND | ND | ND | ND | ND | 88 | 769 |
| TG 47:2 | ND | ND | ND | 50 | ND | ND | ND |
| TG 47:2 \| TG 14:0_15:0_18:2 | ND | ND | ND | ND | ND | 74 | ND |
| TG 47:2 \| TG 16:0_15:1_16:1 | ND | ND | ND | ND | ND | ND | 768 |
| TG 47:3 | ND | ND | ND | 594 | ND | ND | ND |
| TG 47:3 \| TG 13:0_16:1_18:2 | ND | ND | ND | ND | ND | 786 | ND |
| TG 47:4 | ND | ND | ND | 98 | ND | 787 | ND |
| TG 48:0 | ND | ND | ND | ND | ND | 204 | ND |
| TG 48:0 \| TG 14:0_16:0_18:0 | ND | ND | ND | 242 | ND | ND | ND |

| | | | | | | | |
|---|---|---|---|---|---|---|---|
| TG 48:1 | ND | ND | ND | ND | ND | 51 | ND |
| TG 48:2 | ND | ND | ND | ND | ND | 105 | ND |
| TG 48:2 \| TG 14:0_16:1_18:1 | ND | ND | ND | 605 | ND | ND | ND |
| TG 48:3 | ND | ND | ND | ND | ND | 145 | ND |
| TG 48:4 | ND | ND | ND | ND | ND | 792 | ND |
| TG 48:4 \| TG 14:1_16:1_18:2 | ND | ND | ND | 607 | ND | ND | ND |
| TG 48:5 | ND | ND | ND | ND | ND | 122 | ND |
| TG 48:5 \| TG 12:0_18:2_18:3 | ND | ND | ND | 608 | ND | ND | ND |
| TG 48:5 \| TG 16:0_16:1_16:4 | ND | ND | ND | 609 | ND | ND | ND |
| TG 49:0 | ND | ND | ND | ND | ND | 254 | ND |
| TG 49:0 \| TG 16:0_16:0_17:0 | ND | ND | ND | 610 | ND | ND | ND |
| TG 49:1 | ND | ND | ND | ND | ND | 350 | ND |
| TG 49:1 \| TG 15:0_16:0_18:1 | ND | ND | ND | 18 | ND | ND | ND |
| TG 49:2 | ND | ND | ND | ND | ND | 796 | ND |
| TG 49:3 | ND | ND | ND | ND | ND | 332 | ND |
| TG 49:4 \| TG 15:1_16:1_18:2 | ND | ND | ND | ND | ND | 124 | 767 |
| TG 49:4 \| TG 16:0_15:2_18:2 | ND | ND | ND | 611 | ND | ND | ND |
| TG 49:5 | ND | ND | ND | 580 | ND | ND | ND |
| TG 49:5 \| TG 13:0_18:2_18:3 | ND | ND | ND | ND | ND | 799 | ND |
| TG 50:1 | ND | ND | ND | ND | ND | 61 | ND |
| TG 50:2 | ND | ND | ND | ND | ND | 249 | ND |
| TG 50:2 \| TG 16:0_16:1_18:1 | ND | ND | ND | 582 | ND | ND | ND |
| TG 50:3 | ND | ND | ND | 583 | ND | 802 | ND |
| TG 50:4 | ND | ND | ND | ND | ND | 107 | ND |
| TG 50:5 | ND | ND | ND | ND | ND | 333 | ND |
| TG 50:7 \| TG 16:1_18:2_16:4 | ND | ND | ND | 43 | ND | ND | ND |
| TG 51:1 | ND | ND | ND | ND | ND | 356 | ND |
| TG 51:2 | ND | ND | ND | ND | ND | 806 | ND |
| TG 51:3 | ND | ND | ND | 691 | ND | 807 | ND |
| TG 51:4 | ND | ND | ND | ND | ND | 808 | ND |
| TG 51:4 \| TG 16:1_17:1_18:2 | ND | ND | ND | 596 | ND | ND | ND |
| TG 51:5 | ND | ND | ND | ND | ND | 246 | ND |
| TG 51:6 \| TG 15:1_18:2_18:3 | ND | ND | ND | ND | ND | 368 | ND |
| TG 51:6 \| TG 15:2_18:2_18:2 | ND | ND | ND | 598 | ND | ND | ND |
| TG 52:0 | ND | ND | ND | ND | ND | 426 | ND |
| TG 52:1 | ND | ND | ND | ND | ND | 46 | ND |
| TG 52:2 | ND | ND | ND | ND | ND | 338 | ND |
| TG 52:2 \| TG 16:0_18:1_18:1 | ND | ND | ND | 600 | ND | ND | ND |
| TG 52:3 | ND | ND | ND | ND | ND | 814 | ND |
| TG 52:3 \| TG 16:0_18:1_18:2 | ND | ND | ND | 180 | ND | ND | ND |
| TG 52:4 | ND | ND | ND | ND | ND | 815 | ND |
| TG 52:4 \| TG 16:1_18:1_18:2 | ND | ND | ND | 601 | ND | ND | ND |
| TG 52:5 | ND | ND | ND | 602 | ND | 816 | ND |

| | | | | | | | |
|---|---|---|---|---|---|---|---|
| TG 52:6 | ND | ND | ND | 603 | ND | ND | ND |
| TG 52:7 \| TG 16:2_18:2_18:3 | ND | ND | ND | 604 | ND | ND | ND |
| TG 52:8 \| TG 14:1_16:1_22:6 | ND | ND | ND | ND | ND | 819 | ND |
| TG 53:0 | ND | ND | ND | ND | ND | 820 | ND |
| TG 53:1 | ND | ND | ND | ND | ND | 821 | ND |
| TG 53:2 | ND | ND | ND | ND | ND | 822 | ND |
| TG 53:3 | ND | ND | ND | ND | ND | 823 | ND |
| TG 53:4 | ND | ND | ND | ND | ND | 824 | ND |
| TG 53:5 | ND | ND | ND | ND | ND | 825 | ND |
| TG 53:7 \| TG 15:0_16:1_22:6 | ND | ND | ND | 616 | ND | 827 | ND |
| TG 54:0 | ND | ND | ND | ND | ND | 828 | ND |
| TG 54:1 | ND | ND | ND | ND | ND | 196 | ND |
| TG 54:1 \| TG 16:0_20:0_18:1 | ND | ND | ND | 617 | ND | ND | ND |
| TG 54:2 | ND | ND | ND | ND | ND | 830 | ND |
| TG 54:3 | ND | ND | ND | ND | ND | 831 | ND |
| TG 54:3 \| TG 18:0_18:1_18:2 | ND | ND | ND | 140 | ND | ND | ND |
| TG 54:4 | ND | ND | ND | ND | ND | 832 | ND |
| TG 54:4 \| TG 18:1_18:1_18:2 | ND | ND | ND | 619 | ND | ND | ND |
| TG 54:5 | ND | ND | ND | 620 | ND | 833 | ND |
| TG 54:6 | ND | ND | ND | 621 | ND | 834 | ND |
| TG 54:7 | ND | ND | ND | 622 | ND | 835 | ND |
| TG 54:8 | ND | ND | ND | ND | ND | 339 | ND |
| TG 54:8 \| TG 16:1_16:1_22:6 | ND | ND | ND | 623 | ND | ND | ND |
| TG 54:8 \| TG 16:1_18:2_20:5 | ND | ND | ND | 658 | ND | ND | ND |
| TG 54:8 \| TG 18:2_18:2_18:3 | ND | ND | ND | 659 | ND | ND | ND |
| TG 54:9 | ND | ND | ND | 628 | ND | ND | ND |
| TG 55:1 | ND | ND | ND | ND | ND | 837 | ND |
| TG 55:1 \| TG 18:0_19:0_18:1 | ND | ND | ND | 629 | ND | ND | ND |
| TG 55:2 | ND | ND | ND | ND | ND | 838 | ND |
| TG 55:2 \| TG 18:0_18:1_19:1 | ND | ND | ND | 630 | ND | ND | ND |
| TG 55:3 | ND | ND | ND | ND | ND | 839 | ND |
| TG 55:4 \| TG 18:1_19:1_18:2 | ND | ND | ND | 632 | ND | 840 | 764 |
| TG 55:8 \| TG 16:1_18:2_21:5 | ND | ND | ND | 636 | ND | ND | 792 |
| TG 56:1 | ND | ND | ND | ND | ND | 845 | ND |
| TG 56:10 | ND | ND | ND | 637 | ND | ND | ND |
| TG 56:1 \| TG 18:0_20:0_18:1 | ND | ND | ND | 638 | ND | ND | ND |
| TG 56:2 | ND | ND | ND | ND | ND | 846 | ND |
| TG 56:2 \| TG 18:0_18:1_20:1 | ND | ND | ND | 639 | ND | ND | ND |
| TG 56:3 | ND | ND | ND | ND | ND | 847 | ND |
| TG 56:3 \| TG 18:1_18:1_20:1 | ND | ND | ND | 640 | ND | ND | ND |
| TG 56:4 | ND | ND | ND | ND | ND | 109 | ND |
| TG 56:5 | ND | ND | ND | ND | ND | 849 | ND |
| TG 56:5 \| TG 18:1_18:2_20:2 | ND | ND | ND | 190 | ND | ND | ND |

| | | | | | | | |
|---|---|---|---|---|---|---|---|
| TG 56:6 | ND | ND | ND | ND | ND | 850 | ND |
| TG 56:6 \| TG 16:0_18:1_22:5 | ND | ND | ND | 644 | ND | ND | ND |
| TG 56:6 \| TG 18:1_18:2_20:3 | ND | ND | ND | 157 | ND | ND | ND |
| TG 56:7 | ND | ND | ND | ND | ND | 851 | ND |
| TG 56:7 \| TG 16:0_18:1_22:6 | ND | ND | ND | 645 | ND | ND | ND |
| TG 56:7 \| TG 18:1_18:2_20:4 | ND | ND | ND | 646 | ND | ND | ND |
| TG 56:8 | ND | ND | ND | ND | ND | 852 | ND |
| TG 56:9 | ND | ND | ND | ND | ND | 853 | ND |
| TG 56:9 \| TG 18:2_18:2_20:5 | ND | ND | ND | 649 | ND | ND | ND |
| TG 57:1 | ND | ND | ND | ND | ND | 854 | ND |
| TG 57:10 \| TG 18:2_18:3_21:5 | ND | ND | ND | 650 | ND | ND | ND |
| TG 57:2 | ND | ND | ND | ND | ND | 855 | ND |
| TG 57:2 \| TG 19:0_18:1_20:1 | ND | ND | ND | 652 | ND | ND | ND |
| TG 57:3 | ND | ND | ND | 653 | ND | ND | ND |
| TG 57:4 \| TG 18:1_21:1_18:2 | ND | ND | ND | 654 | ND | 857 | 790 |
| TG 57:5 \| TG 21:1_18:2_18:2 | ND | ND | ND | 655 | ND | 858 | ND |
| TG 57:7 \| TG 17:0_18:1_22:6 | ND | ND | ND | 656 | ND | ND | ND |
| TG 57:7 \| TG 18:1_18:1_21:5 | ND | ND | ND | 657 | ND | 859 | 789 |
| TG 57:9 \| TG 18:2_18:2_21:5 | ND | ND | ND | 662 | ND | 861 | ND |
| TG 58:1 | ND | ND | ND | ND | ND | 862 | ND |
| TG 58:10 | ND | ND | ND | ND | ND | 863 | ND |
| TG 58:11 \| TG 18:2_18:3_22:6 | ND | ND | ND | 664 | ND | 864 | ND |
| TG 58:11 \| TG 18:2_20:4_20:5 | ND | ND | ND | ND | ND | ND | 788 |
| TG 58:2 | ND | ND | ND | ND | ND | 403 | ND |
| TG 58:3 | ND | ND | ND | ND | ND | 394 | ND |
| TG 58:4 | ND | ND | ND | ND | ND | 868 | ND |
| TG 58:4 \| TG 18:1_22:1_18:2 | ND | ND | ND | 669 | ND | ND | ND |
| TG 58:5 | ND | ND | ND | ND | ND | 138 | ND |
| TG 58:6 | ND | ND | ND | ND | ND | 870 | ND |
| TG 58:7 \| TG 18:1_18:1_22:5 | ND | ND | ND | 673 | ND | 871 | ND |
| TG 58:8 | ND | ND | ND | 674 | ND | 872 | ND |
| TG 58:9 | ND | ND | ND | 702 | ND | 873 | ND |
| TG 59:2 | ND | ND | ND | ND | ND | 875 | ND |
| TG 59:2 \| TG 23:0_18:1_18:1 | ND | ND | ND | 705 | ND | ND | ND |
| TG 59:3 | ND | ND | ND | ND | ND | 211 | ND |
| TG 59:3 \| TG 23:0_18:1_18:2 | ND | ND | ND | 247 | ND | ND | ND |
| TG 59:4 \| TG 18:1_23:1_18:2 | ND | ND | ND | 211 | ND | 877 | 786 |
| TG 59:5 \| TG 23:1_18:2_18:2 | ND | ND | ND | 52 | ND | 280 | 785 |
| TG 59:7 \| TG 19:0_18:1_22:6 | ND | ND | ND | ND | ND | ND | 784 |
| TG 60:10 | ND | ND | ND | 684 | ND | 408 | ND |
| TG 60:10 \| TG 18:2_20:3_22:5 | ND | ND | ND | ND | ND | ND | 783 |
| TG 60:11 | ND | ND | ND | 685 | ND | 880 | ND |
| TG 60:12 \| TG 16:0_22:6_22:6 | ND | ND | ND | 686 | ND | 881 | 782 |

| | | | | | | | |
|---|---|---|---|---|---|---|---|
| TG 60:13 \| TG 16:1_22:6_22:6 | ND | ND | ND | ND | ND | 882 | ND |
| TG 60:2 | ND | ND | ND | ND | ND | 883 | ND |
| TG 60:2 \| TG 18:0_18:1_24:1 | ND | ND | ND | 688 | ND | ND | ND |
| TG 60:3 | ND | ND | ND | ND | ND | 884 | ND |
| TG 60:4 | ND | ND | ND | ND | ND | 264 | ND |
| TG 60:5 | ND | ND | ND | ND | ND | 28 | ND |
| TG 60:6 | ND | ND | ND | ND | ND | 18 | ND |
| TG 60:6 \| TG 18:1_20:1_22:4 | ND | ND | ND | 113 | ND | ND | ND |
| TG 60:7 | ND | ND | ND | ND | ND | 59 | ND |
| TG 60:8 \| TG 18:1_18:2_24:5 | ND | ND | ND | 721 | ND | ND | 781 |
| TG 60:9 | ND | ND | ND | ND | ND | 890 | ND |
| TG 60:9 \| TG 20:2_20:3_20:4 | ND | ND | ND | ND | ND | ND | 780 |
| TG 61:1 \| TG 18:0_25:0_18:1 | ND | ND | ND | 692 | ND | ND | ND |
| TG 61:2 \| TG 25:0_18:1_18:1 | ND | ND | ND | 83 | ND | 891 | ND |
| TG 61:4 \| TG 18:1_25:1_18:2 | ND | ND | ND | 694 | ND | 893 | ND |
| TG 62:13 \| TG 18:2_22:5_22:6 | ND | ND | ND | 697 | ND | ND | 247 |
| TG 62:1 \| TG 16:0_28:0_18:1 | ND | ND | ND | 699 | ND | ND | ND |
| TG 62:2 | ND | ND | ND | ND | ND | 897 | ND |
| TG 62:2 \| TG 26:0_18:1_18:1 | ND | ND | ND | 700 | ND | ND | ND |
| TG 62:3 | ND | ND | ND | ND | ND | 898 | ND |
| TG 62:4 | ND | ND | ND | ND | ND | 182 | ND |
| TG 62:4 \| TG 18:1_26:1_18:2 | ND | ND | ND | 284 | ND | ND | ND |
| TG 62:5 \| TG 26:1_18:2_18:2 | ND | ND | ND | 708 | ND | 900 | ND |
| TG 62:7 \| TG 22:0_18:1_22:6 | ND | ND | ND | ND | ND | ND | 778 |
| TG 63:2 \| TG 16:0_29:0_18:2 | ND | ND | ND | 182 | ND | ND | ND |
| TG 63:3 \| TG 27:0_18:1_18:2 | ND | ND | ND | 74 | ND | 287 | ND |
| TG 63:4 \| TG 18:1_27:1_18:2 | ND | ND | ND | ND | ND | 902 | ND |
| TG 64:11 \| TG 18:2_18:2_28:7 | ND | ND | ND | 709 | ND | 3 | ND |
| TG 64:1 \| TG 16:0_30:0_18:1 | ND | ND | ND | 254 | ND | ND | ND |
| TG 64:2 \| TG 16:0_18:1_30:1 | ND | ND | ND | 711 | ND | ND | ND |
| TG 64:3 | ND | ND | ND | ND | ND | 904 | ND |
| TG 64:3 \| TG 28:0_18:1_18:2 | ND | ND | ND | 712 | ND | ND | ND |
| TG 64:4 | ND | ND | ND | ND | ND | 905 | ND |
| TG 64:4 \| TG 18:1_28:1_18:2 | ND | ND | ND | 235 | ND | ND | ND |
| TG 64:5 \| TG 28:1_18:2_18:2 | ND | ND | ND | ND | ND | 906 | ND |
| TG 65:3 \| TG 16:0_31:1_18:2 | ND | ND | ND | ND | ND | 99 | ND |
| TG 65:3 \| TG 29:0_18:1_18:2 | ND | ND | ND | 713 | ND | ND | ND |
| TG 65:4 \| TG 18:1_29:1_18:2 | ND | ND | ND | 714 | ND | 908 | ND |
| TG 66:3 \| TG 30:0_18:1_18:2 | ND | ND | ND | 715 | ND | 909 | ND |
| TG 66:4 \| TG 18:1_30:1_18:2 | ND | ND | ND | 716 | ND | ND | ND |
| TG 66:4 \| TG 30:0_18:2_18:2 | ND | ND | ND | ND | ND | 910 | ND |
| TG 66:5 \| TG 30:1_18:2_18:2 | ND | ND | ND | 717 | ND | 911 | 807 |
| TG 67:4 \| TG 18:1_31:1_18:2 | ND | ND | ND | 718 | ND | 418 | ND |

| | | | | | | | |
|---|---|---|---|---|---|---|---|
| TG 67:5 \| TG 31:1_18:2_18:2 | ND | ND | ND | ND | ND | 192 | ND |
| TG 68:3;O2 \| TG 16:0_18:1_18:1;O(FA 16:0) | ND | ND | ND | 719 | ND | ND | ND |
| TG 68:4;O2 \| TG 16:0_18:2_16:0;O(FA 18:1) | ND | ND | ND | 282 | ND | ND | ND |
| TG 68:4 \| TG 18:1_32:1_18:2 | ND | ND | ND | ND | ND | 914 | ND |
| TG 68:5 \| TG 32:1_18:2_18:2 | ND | ND | ND | 523 | ND | 915 | 806 |
| TG 69:5 \| TG 33:1_18:2_18:2 | ND | ND | ND | ND | ND | 916 | ND |
| TG 70:3;O2 \| TG 16:0_18:1_18:1;O(FA 18:0) | ND | ND | ND | 521 | ND | ND | 204 |
| TG 70:3 \| TG 18:1_18:1_34:1 | ND | ND | ND | ND | ND | ND | 805 |
| TG 70:4;O2 \| TG 16:0_18:1_18:1;O(FA 18:1) | ND | ND | ND | 519 | ND | ND | ND |
| TG 70:5;O2 \| TG 18:1_18:2_18:1;O(FA 16:0) | ND | ND | ND | 745 | ND | ND | ND |
| TG 70:5 \| TG 34:1_18:2_18:2 | ND | ND | ND | 746 | ND | 917 | 804 |
| TG 70:6;O2 \| TG 16:0_18:2_18:1;O(FA 18:2) | ND | ND | ND | 747 | ND | ND | ND |
| TG 71:4 \| TG 18:1_35:1_18:2 | ND | ND | ND | ND | ND | 918 | ND |
| TG 72:5;O2 \| TG 18:1_18:2_18:1;O(FA 18:0) | ND | ND | ND | 748 | ND | ND | ND |
| TG 72:5 \| TG 36:1_18:2_18:2 | ND | ND | ND | ND | ND | 919 | 803 |
| TG 72:6;O2 \| TG 18:1_18:2_18:1;O(FA 18:1) | ND | ND | ND | 749 | ND | ND | ND |
| TG 72:7;O2 \| TG 18:1_18:2_18:2;O(FA 18:1) | ND | ND | ND | 750 | ND | ND | ND |
| TG O-38:2 | ND | ND | ND | 751 | ND | ND | ND |
| TG O-43:7 \| TG O-13:1_15:3_15:3 | ND | ND | ND | 752 | ND | ND | ND |
| TG O-43:8 \| TG O-9:0_17:4_17:4 | ND | ND | ND | 753 | ND | ND | ND |
| TG O-44:10 \| TG O-8:0_18:5_18:5 | ND | ND | ND | 754 | ND | ND | ND |
| TG O-50:0 | ND | ND | ND | 227 | ND | 920 | ND |
| TG O-50:1 | ND | ND | ND | 22 | ND | 921 | ND |
| TG O-50:2 | ND | ND | ND | 728 | ND | 94 | ND |
| TG O-52:0 | ND | ND | ND | 95 | ND | ND | ND |
| TG O-52:1 | ND | ND | ND | 729 | ND | 923 | ND |
| TG O-52:2 | ND | ND | ND | 730 | ND | 924 | ND |
| TG O-52:3 | ND | ND | ND | 162 | ND | 299 | ND |
| TG O-52:4 | ND | ND | ND | ND | ND | 410 | ND |
| TG O-52:4 \| TG O-19:2_15:0_18:2 | ND | ND | ND | 731 | ND | ND | ND |
| TG O-52:6 | ND | ND | ND | ND | ND | 20 | ND |
| TG O-54:1 | ND | ND | ND | 732 | ND | 928 | ND |
| TG O-54:2 | ND | ND | ND | 733 | ND | 312 | ND |
| TG O-54:3 | ND | ND | ND | 734 | ND | 930 | ND |
| TG O-54:4 | ND | ND | ND | 166 | ND | 96 | ND |
| TG O-54:6 | ND | ND | ND | 518 | ND | 253 | ND |
| TG O-54:7 | ND | ND | ND | ND | ND | 154 | ND |
| TG O-55:1 | ND | ND | ND | ND | ND | 21 | ND |
| TG O-56:2 | ND | ND | ND | 735 | ND | 935 | ND |
| TG O-56:3 | ND | ND | ND | ND | ND | 307 | ND |
| TG O-56:4 | ND | ND | ND | ND | ND | 937 | ND |
| TG O-56:6 | ND | ND | ND | 736 | ND | ND | ND |
| TG O-56:7 | ND | ND | ND | 31 | ND | 45 | ND |

| | | | | | | | |
|---|---|---|---|---|---|---|---|
| TG O-56:8 | ND | ND | ND | ND | ND | 27 | ND |
| TG O-58:4 | ND | ND | ND | ND | ND | 373 | ND |
| TG O-58:6 | ND | ND | ND | ND | ND | 353 | ND |
| TG O-58:8 | ND | ND | ND | ND | ND | 181 | ND |
| Tapentadol | ND | ND | 122 | ND | ND | ND | ND |
| Targinine | ND | ND | ND | ND | ND | ND | 802 |
| Testosterone | ND | 402 | ND | ND | ND | ND | ND |
| Tetraethylene glycol | ND | 55 | 206 | ND | ND | ND | ND |
| Tetraethylene glycol_b | ND | ND | ND | ND | ND | 945 | ND |
| Tetramethylammonium | ND | 56 | ND | ND | ND | ND | ND |
| Thiamine cation | ND | ND | ND | ND | ND | ND | 799 |
| Thiamine monophosphate | ND | ND | ND | ND | ND | ND | 798 |
| Thr-Ala | ND | ND | ND | ND | ND | 388 | ND |
| Thr-Arg | ND | ND | ND | ND | 14 | 948 | 797 |
| Thr-Lys | ND | ND | ND | ND | ND | 950 | 794 |
| Thr-Ser | ND | ND | ND | ND | ND | 951 | ND |
| Thr-Thr | ND | 412 | ND | ND | ND | ND | ND |
| Threonine | ND | ND | 386 | 545 | 503 | ND | 793 |
| Trehalose | ND | ND | ND | ND | ND | ND | 835 |
| Triallyl cyanurate | ND | ND | 251 | ND | ND | ND | ND |
| Tributyl phosphate | ND | 169 | ND | ND | ND | ND | ND |
| Tridodecylmethylammonium | ND | ND | ND | 222 | ND | ND | ND |
| Trigonelline | ND | ND | ND | ND | ND | 42 | ND |
| Trimethylamine N-oxide | ND | ND | ND | ND | ND | 305 | ND |
| Tripelennamine | ND | 482 | 324 | 225 | ND | ND | ND |
| Tripropylamine | ND | 563 | ND | ND | ND | ND | ND |
| Tris(2-butoxyethyl) phosphate | ND | 11 | ND | ND | ND | ND | ND |
| Tryptophan | ND | 551 | 391 | 569 | ND | ND | ND |
| Tyr-Arg | ND | ND | ND | 581 | 108 | 954 | 821 |
| Tyr-His | ND | ND | ND | ND | 505 | ND | 820 |
| Tyrosine | ND | ND | ND | 626 | ND | ND | 819 |
| Tyrosine butyl ester | ND | ND | 190 | ND | ND | ND | ND |
| UDP GlcNAc | ND | ND | ND | ND | 394 | ND | ND |
| UDP-N-acetylglucosamine | ND | 288 | ND | ND | ND | ND | 818 |
| UDP-glucuronic acid | ND | ND | ND | ND | 95 | ND | 817 |
| UMP | ND | 523 | ND | ND | 140 | ND | 816 |
| Urea | ND | ND | 369 | 682 | ND | 955 | ND |
| Uridine 5'-monophosphate | ND | ND | ND | ND | ND | 956 | ND |
| Uridine-5-diphosphoacetylgalactosamine | ND | ND | ND | ND | 386 | ND | 814 |
| Uridine-5-diphosphoacetylglucosamine | ND | 548 | ND | ND | ND | ND | ND |
| Val-Ala | ND | ND | ND | ND | ND | ND | 217 |
| Val-Arg | ND | 530 | 344 | 706 | 399 | 957 | 813 |
| Val-Glu_a | ND | ND | ND | ND | ND | 222 | ND |

| | | | | | | | |
|---|---|---|---|---|---|---|---|
| Val-His | ND | 549 | ND | ND | 506 | 417 | 812 |
| Val-Leu | ND | ND | ND | ND | ND | 961 | ND |
| Val-Lys | ND | ND | ND | ND | ND | ND | 811 |
| Val-Phe | ND | ND | ND | ND | ND | 80 | ND |
| Val-Ser | ND | ND | ND | ND | ND | 173 | ND |
| Val-Thr | ND | ND | ND | ND | ND | ND | 810 |
| Val-Val | ND | ND | ND | ND | ND | 964 | ND |
| Valinamide | ND | ND | ND | ND | ND | 201 | ND |
| [(4,6-Dimethyl-2-pyrimidinyl)amino]acetic acid | ND | 92 | ND | ND | ND | ND | ND |
| acetaminophen | ND | ND | ND | ND | 115 | ND | ND |
| acetylcarnitine | ND | ND | 345 | ND | ND | ND | ND |
| aconitic acid | ND | ND | ND | ND | ND | 328 | 136 |
| adenine | ND | ND | ND | ND | ND | 197 | ND |
| adenosine | ND | 87 | ND | ND | ND | ND | ND |
| adenosine-5-monophosphate | ND | 289 | ND | ND | ND | 281 | ND |
| adipic acid | ND | 115 | ND | ND | 168 | 340 | 98 |
| alanine | ND | 153 | ND | ND | 162 | 184 | 261 |
| alpha-Galactosamine-1-phosphate | ND | 497 | ND | 742 | ND | ND | ND |
| alpha-aminoadipic acid | ND | 277 | ND | ND | 73 | ND | 165 |
| alpha-ketoglutarate | ND | ND | ND | ND | ND | ND | 93 |
| aminomalonate | ND | 209 | ND | ND | 174 | ND | ND |
| aminomalonic acid | ND | ND | ND | ND | ND | 286 | 838 |
| arabinose | ND | ND | ND | ND | 121 | ND | ND |
| arabitol | ND | 287 | ND | ND | 80 | 972 | ND |
| arachidic acid | ND | ND | ND | ND | ND | 370 | 124 |
| asparagine | ND | 325 | ND | ND | ND | ND | ND |
| aspartate | ND | ND | ND | ND | ND | 363 | ND |
| aspartic acid | ND | ND | ND | ND | 332 | ND | ND |
| azelaic acid | ND | 300 | ND | ND | 126 | 261 | 143 |
| behenic acid | ND | ND | ND | ND | ND | 325 | ND |
| benzoic acid | ND | 237 | ND | ND | ND | ND | 837 |
| beta-Glycerolphosphate | ND | 364 | ND | ND | ND | ND | ND |
| beta-Glycerophosphate | ND | ND | ND | ND | 12 | ND | ND |
| beta-Homomethionine | ND | ND | ND | ND | ND | 337 | ND |
| beta-Nicotinamide adenine dinucleotide | ND | 510 | ND | ND | 197 | 400 | ND |
| beta-alanine | ND | 274 | ND | ND | 97 | 123 | 118 |
| beta-gentiobiose | ND | ND | ND | ND | ND | 981 | ND |
| beta-glycerolphosphate | ND | ND | ND | ND | ND | ND | 189 |
| beta.-Homoglutamine | ND | 226 | ND | ND | ND | ND | ND |
| butane-2,3-diol | ND | ND | ND | ND | ND | 982 | ND |
| butyrolactam | ND | 251 | ND | ND | ND | ND | 836 |
| cadaverine | ND | ND | ND | ND | ND | ND | 139 |
| caproic acid | ND | ND | ND | ND | ND | ND | 133 |

| | | | | | | | |
|---|---|---|---|---|---|---|---|
| caproic acid HMDB | ND | ND | ND | ND | ND | 329 | ND |
| caprylic acid | ND | ND | ND | ND | ND | ND | 85 |
| cellobiose | ND | ND | ND | ND | ND | 207 | 233 |
| cerotinic acid | ND | ND | ND | ND | ND | 985 | ND |
| citric acid | ND | 326 | ND | ND | 244 | 425 | 869 |
| conduritol-beta-epoxide | ND | 72 | ND | ND | ND | 164 | 251 |
| conduritol-beta-expoxide | ND | ND | ND | ND | 177 | ND | ND |
| creatinine | ND | 90 | ND | ND | 171 | 988 | ND |
| cyanoalanine | ND | ND | ND | ND | ND | 989 | 246 |
| cysteine | ND | 139 | ND | ND | 114 | 404 | 282 |
| cysteine sulfonic acid | ND | ND | ND | ND | ND | ND | 860 |
| cysteine-glycine | ND | ND | ND | ND | 153 | ND | ND |
| cytidine-5-monophosphate | ND | 310 | ND | ND | ND | ND | ND |
| dAMP | ND | 341 | ND | ND | ND | ND | ND |
| decane | ND | ND | ND | ND | ND | ND | 123 |
| dehydroascorbic acid | ND | 75 | ND | ND | ND | 290 | 859 |
| deoxypentitol | ND | ND | ND | ND | 68 | ND | 858 |
| diethanolamine | ND | ND | ND | ND | ND | ND | 244 |
| digalacturonic acid | ND | ND | ND | ND | ND | 992 | ND |
| diglycerol | ND | ND | ND | ND | ND | 317 | ND |
| dimethyl-PE 38:4 | ND | ND | ND | ND | ND | ND | 856 |
| dimethyl-lysine | ND | ND | 374 | ND | ND | ND | ND |
| dodecanol | ND | ND | ND | ND | ND | 241 | ND |
| erythritol | ND | 406 | ND | ND | 89 | 415 | 855 |
| erythronic acid | ND | ND | ND | ND | 43 | ND | ND |
| erythronic acid lactone | ND | 240 | ND | ND | ND | ND | ND |
| erythronic acid lactone isomer | ND | ND | ND | ND | ND | ND | 854 |
| erythrose major | ND | ND | ND | ND | ND | ND | 853 |
| ethanolamine | ND | 93 | ND | ND | 155 | ND | 852 |
| flavin adenine | ND | ND | ND | ND | ND | ND | 154 |
| free fatty acid 14:0 (myristic acid) | ND | ND | 153 | ND | ND | ND | ND |
| free fatty acid 15:1 | ND | ND | 39 | ND | ND | ND | ND |
| free fatty acid 16:0 (palmitic acid) | ND | ND | 95 | ND | ND | ND | ND |
| free fatty acid 16:1 (palmitoleic acid) | ND | ND | 182 | ND | ND | ND | ND |
| free fatty acid 16:2 | ND | ND | 144 | ND | ND | ND | ND |
| free fatty acid 16:3 | ND | ND | 85 | ND | ND | ND | ND |
| free fatty acid 16:4 | ND | ND | 138 | ND | ND | ND | ND |
| free fatty acid 17:1 | ND | ND | 131 | ND | ND | ND | ND |
| free fatty acid 17:2 | ND | ND | 14 | ND | ND | ND | ND |
| free fatty acid 18:1 | ND | ND | 260 | ND | ND | ND | ND |
| free fatty acid 18:2 (linoleic acid) | ND | ND | 284 | ND | ND | ND | ND |
| free fatty acid 18:3 | ND | ND | 164 | ND | ND | ND | ND |
| free fatty acid 18:4 | ND | ND | 38 | ND | ND | ND | ND |

| | | | | | | | |
|---|---|---|---|---|---|---|---|
| free fatty acid 19:0 | ND | ND | 36 | ND | ND | ND | ND |
| free fatty acid 19:1 | ND | ND | 232 | ND | ND | ND | ND |
| free fatty acid 19:2 | ND | ND | 89 | ND | ND | ND | ND |
| free fatty acid 20:1 (eicosenoic acid) | ND | ND | 47 | ND | ND | ND | ND |
| free fatty acid 20:2 (eicosadienoic acid) | ND | ND | 83 | ND | ND | ND | ND |
| free fatty acid 20:3 (homo-gamma-linolenic acid) | ND | ND | 90 | ND | ND | ND | ND |
| free fatty acid 20:4 (arachidonic acid) | ND | ND | 29 | ND | ND | ND | ND |
| free fatty acid 20:5 (eicosapentaenoic acid) | ND | ND | 46 | ND | ND | ND | ND |
| free fatty acid 21:1 | ND | ND | 81 | ND | ND | ND | ND |
| free fatty acid 21:5 | ND | ND | 103 | ND | ND | ND | ND |
| free fatty acid 22:1 (erucic acid) | ND | ND | 119 | ND | ND | ND | ND |
| free fatty acid 22:2 (docosadienoic acid) | ND | ND | 256 | ND | ND | ND | ND |
| free fatty acid 22:3 | ND | ND | 197 | ND | ND | ND | ND |
| free fatty acid 22:4 | ND | ND | 93 | ND | ND | ND | ND |
| free fatty acid 22:5 | ND | ND | 45 | ND | ND | ND | ND |
| free fatty acid 22:6 (docosahexaenoic acid) | ND | ND | 86 | ND | ND | ND | ND |
| free fatty acid 23:1 | ND | ND | 5 | ND | ND | ND | ND |
| free fatty acid 24:1 (nervonic acid) | ND | ND | 230 | ND | ND | ND | ND |
| free fatty acid 24:2 | ND | ND | 186 | ND | ND | ND | ND |
| free fatty acid 24:4 | ND | ND | 221 | ND | ND | ND | ND |
| free fatty acid 24:5 | ND | ND | 105 | ND | ND | ND | ND |
| free fatty acid 24:6 | ND | ND | 63 | ND | ND | ND | ND |
| free fatty acid 25:1 | ND | ND | 71 | ND | ND | ND | ND |
| free fatty acid 28:7 | ND | ND | 7 | ND | ND | ND | ND |
| free fatty acid 34:1 | ND | ND | 74 | ND | ND | ND | ND |
| free fatty acid 44:10 | ND | ND | 34 | ND | ND | ND | ND |
| fructose | ND | 392 | ND | ND | 144 | 233 | 851 |
| fructose-1-phosphate | ND | ND | ND | ND | 172 | ND | 884 |
| fructose-6-phosphate | ND | ND | ND | ND | 154 | 997 | 234 |
| fucose | ND | ND | ND | ND | ND | ND | 881 |
| fumaric acid | ND | 281 | ND | ND | 184 | 203 | 880 |
| galactinol | ND | ND | ND | ND | 228 | ND | 185 |
| galactitol | ND | ND | ND | ND | ND | 999 | ND |
| galactonic acid | ND | ND | ND | ND | 327 | 148 | 879 |
| galacturonic acid | ND | ND | ND | ND | 375 | ND | ND |
| gamma-Aminobutyric Acid (GABA) | ND | ND | ND | ND | ND | ND | 878 |
| gamma-Glutamyl-cysteine | ND | ND | ND | ND | ND | ND | 877 |
| gamma-Glutamyl-glutamic acid | ND | ND | ND | ND | ND | 169 | ND |
| gamma.-L-Glutamyl-L-alanine | ND | ND | ND | ND | ND | ND | 875 |
| gamma.-Muricholic acid | ND | ND | ND | ND | ND | ND | 296 |
| gluconic acid | ND | 256 | ND | ND | 260 | ND | 276 |
| gluconic acid lactone | ND | ND | ND | ND | 18 | ND | ND |
| glucose | ND | 342 | ND | ND | 185 | 1002 | 872 |

| | | | | | | | |
|---|---|---|---|---|---|---|---|
| glucose-1-phosphate | ND | 329 | ND | ND | ND | 1003 | ND |
| glucose-6-phosphate | ND | 350 | ND | ND | 25 | 1004 | 286 |
| glucuronic acid | ND | ND | ND | ND | 230 | ND | 241 |
| glutamic acid | ND | ND | ND | ND | ND | 344 | 870 |
| glutamine | ND | ND | ND | ND | ND | 1006 | ND |
| glutaric acid | ND | 242 | ND | ND | 147 | 273 | 220 |
| glutathione | ND | ND | ND | ND | ND | 422 | ND |
| glyceraldehyde | ND | ND | ND | ND | ND | ND | 894 |
| glyceric acid | ND | 467 | ND | ND | 167 | 1009 | 893 |
| glycerol | ND | 168 | ND | ND | 100 | 1010 | 179 |
| glycerol-3-galactoside | ND | ND | ND | ND | 205 | 362 | 280 |
| glycine | ND | 104 | ND | ND | 558 | 141 | 890 |
| glycocyamine | ND | ND | ND | ND | ND | 1014 | ND |
| glycolic acid | ND | 252 | ND | ND | 275 | 365 | 228 |
| glycyl-glycine | ND | ND | ND | ND | ND | ND | 889 |
| guanidinosuccinate | ND | 61 | ND | ND | 143 | ND | ND |
| guanine | ND | 301 | ND | ND | ND | ND | ND |
| histidine | ND | ND | ND | ND | 194 | ND | ND |
| homocitrulline | ND | 76 | ND | ND | ND | ND | ND |
| homoserine | ND | 269 | ND | ND | ND | ND | ND |
| hydroxycarbamate | ND | ND | ND | ND | ND | 1016 | ND |
| hypotaurine | ND | 327 | ND | ND | ND | 1017 | 142 |
| hypoxanthine | ND | 369 | ND | ND | ND | ND | ND |
| indole-3-lactate | ND | ND | ND | ND | ND | ND | 888 |
| inosine | ND | 148 | ND | ND | ND | 1018 | ND |
| inositol-4-monophosphate | ND | 319 | ND | ND | 187 | 1019 | 194 |
| isoleucine | ND | 79 | ND | ND | 560 | 306 | 887 |
| isomaltose | ND | ND | ND | ND | 263 | ND | 274 |
| isooctanol | ND | ND | ND | ND | ND | 252 | 229 |
| isothreonic acid | ND | 450 | ND | ND | 297 | 1022 | 885 |
| itaconic acid | ND | 49 | ND | ND | ND | ND | ND |
| lactamide | ND | ND | ND | ND | ND | 407 | 839 |
| lactic acid | ND | 166 | ND | ND | 215 | 291 | 906 |
| lactose | ND | ND | ND | ND | 40 | ND | 271 |
| leucine | ND | 100 | ND | ND | ND | 1025 | 904 |
| levoglucosan | ND | 280 | ND | ND | 212 | 1026 | 170 |
| lignoceric acid | ND | ND | ND | ND | 49 | 1027 | ND |
| linoleic acid | ND | ND | ND | ND | 352 | 1028 | 50 |
| lysine | ND | ND | ND | ND | ND | 1029 | ND |
| lysophosphatidylcholine 16:0 | ND | ND | 244 | ND | ND | ND | ND |
| lysophosphatidylcholine 18:0 | ND | ND | 23 | ND | ND | ND | ND |
| lysophosphatidylcholine 18:1 | ND | ND | 109 | ND | ND | ND | ND |
| lysophosphatidylcholine 18:2 | ND | ND | 125 | ND | ND | ND | ND |

| | | | | | | | |
|---|---|---|---|---|---|---|---|
| lysophosphatidylcholine 20:4 | ND | ND | 217 | ND | ND | ND | ND |
| lysophosphatidylethanolamine 16:0 | ND | ND | 52 | ND | ND | ND | ND |
| lysophosphatidylethanolamine 18:0 | ND | ND | 333 | ND | ND | ND | ND |
| lysophosphatidylethanolamine 18:1 | ND | ND | 150 | ND | ND | ND | ND |
| lysophosphatidylethanolamine 18:2 | ND | ND | 124 | ND | ND | ND | ND |
| lysophosphatidylethanolamine 20:4 | ND | ND | 268 | ND | ND | ND | ND |
| lysophosphatidylethanolamine 22:6 | ND | ND | 76 | ND | ND | ND | ND |
| lysophosphatidylethanolamine O-16:1 | ND | ND | 307 | ND | ND | ND | ND |
| lysophosphatidylinositol 18:0 | ND | ND | 246 | ND | ND | ND | ND |
| lyxitol | ND | 223 | ND | ND | 90 | ND | ND |
| maleic acid | ND | 411 | ND | ND | 192 | 424 | 903 |
| maleimide | ND | 339 | ND | ND | ND | ND | 257 |
| malic acid | ND | 458 | ND | ND | 576 | 238 | 94 |
| malonamide | ND | ND | ND | ND | ND | ND | 901 |
| maltose | ND | 420 | ND | ND | ND | 1032 | ND |
| maltotriose | ND | ND | ND | ND | ND | 1033 | ND |
| mannitol | ND | ND | ND | ND | ND | ND | 301 |
| mannose | ND | ND | ND | ND | 338 | 341 | 899 |
| melezitose | ND | ND | ND | ND | ND | ND | 267 |
| melibiose | ND | ND | ND | ND | 110 | 1035 | ND |
| methanolphosphate | ND | 232 | ND | ND | 577 | 217 | 897 |
| methionine sulfoxide | ND | 401 | ND | ND | 276 | ND | ND |
| methylmalonic acid | ND | ND | ND | ND | ND | 278 | ND |
| myo-inositol | ND | 273 | ND | ND | 578 | 83 | 896 |
| n-Propyl gallate | ND | ND | 285 | ND | ND | ND | ND |
| nicotinamide | ND | ND | ND | ND | 317 | ND | 895 |
| nicotinic acid | ND | ND | ND | ND | 176 | 1040 | 226 |
| octadecanol | ND | 134 | ND | ND | 240 | 1041 | 201 |
| oleamide | ND | ND | ND | ND | 300 | 161 | 129 |
| oleoylcarnitine | ND | ND | 183 | ND | ND | ND | ND |
| ornithine | ND | ND | ND | ND | ND | 1043 | ND |
| oxalic acid | ND | 106 | ND | ND | ND | 142 | 874 |
| oxoproline | ND | 388 | ND | ND | ND | 1045 | ND |
| p-Aminobenzoic acid | ND | ND | ND | ND | ND | ND | 873 |
| p-hydroxylphenyllactic acid | ND | ND | ND | ND | ND | 1046 | ND |
| palmitoylcarnitine | ND | ND | 212 | ND | ND | ND | ND |
| panose | ND | ND | ND | ND | ND | 1047 | 157 |
| pantothenic acid | ND | ND | ND | ND | ND | ND | 871 |
| parabanic acid | ND | ND | ND | ND | ND | 1048 | 868 |
| pentose | ND | ND | ND | ND | 272 | ND | ND |
| phenol | ND | ND | ND | ND | ND | ND | 237 |
| phenylalanine | ND | 188 | ND | ND | 381 | 73 | 863 |
| phenylethylamine | ND | ND | ND | ND | ND | ND | 862 |

| | | | | | | | |
|---|---|---|---|---|---|---|---|
| phosphate | ND | 321 | ND | ND | 419 | 1050 | 242 |
| phosphatidylcholine 30:0 | ND | ND | 338 | ND | ND | ND | ND |
| phosphatidylcholine 32:0 | ND | ND | 301 | ND | ND | ND | ND |
| phosphatidylcholine 34:0 | ND | ND | 355 | ND | ND | ND | ND |
| phosphatidylcholine 34:2 | ND | ND | 278 | ND | ND | ND | ND |
| phosphatidylcholine 35:1 | ND | ND | 292 | ND | ND | ND | ND |
| phosphatidylcholine 35:2 | ND | ND | 108 | ND | ND | ND | ND |
| phosphatidylcholine 36:3 | ND | ND | 340 | ND | ND | ND | ND |
| phosphatidylcholine 36:4 | ND | ND | 308 | ND | ND | ND | ND |
| phosphatidylcholine 36:5 | ND | ND | 199 | ND | ND | ND | ND |
| phosphatidylcholine 37:2 | ND | ND | 21 | ND | ND | ND | ND |
| phosphatidylcholine 38:3 | ND | ND | 273 | ND | ND | ND | ND |
| phosphatidylcholine 38:5 | ND | ND | 225 | ND | ND | ND | ND |
| phosphatidylcholine 38:6 | ND | ND | 136 | ND | ND | ND | ND |
| phosphatidylcholine 40:5 | ND | ND | 290 | ND | ND | ND | ND |
| phosphatidylcholine 40:6 | ND | ND | 261 | ND | ND | ND | ND |
| phosphatidylcholine 40:7 | ND | ND | 319 | ND | ND | ND | ND |
| phosphatidylcholine 40:8 | ND | ND | 314 | ND | ND | ND | ND |
| phosphatidylcholine O-34:1 | ND | ND | 169 | ND | ND | ND | ND |
| phosphatidylcholine O-34:2 | ND | ND | 121 | ND | ND | ND | ND |
| phosphatidylcholine O-36:4 | ND | ND | 133 | ND | ND | ND | ND |
| phosphatidylethanolaminentaethylene glycol | ND | ND | 156 | ND | ND | ND | ND |
| phosphatidylethanolaminerindopril | ND | ND | 114 | ND | ND | ND | ND |
| phosphoenolpyruvate | ND | ND | ND | ND | ND | ND | 861 |
| phosphogluconic acid | ND | ND | ND | ND | 232 | ND | 845 |
| phthalic acid | ND | 196 | ND | ND | ND | ND | ND |
| pimelic acid | ND | ND | ND | ND | ND | 361 | 279 |
| pinitol | ND | ND | ND | ND | 170 | 39 | 218 |
| pipecolinic acid | ND | ND | ND | ND | ND | ND | 654 |
| proline | ND | 218 | ND | ND | 281 | 224 | 647 |
| pseudo uridine | ND | 183 | ND | ND | 329 | 1055 | 76 |
| putrescine | ND | 313 | ND | ND | 188 | 547 | 621 |
| pyrophosphate | ND | 381 | ND | ND | 159 | 1073 | 248 |
| pyrrole-2-carboxylic acid | ND | ND | ND | ND | ND | 1074 | ND |
| pyruvic acid | ND | ND | ND | ND | ND | ND | 616 |
| raffinose | ND | ND | ND | ND | ND | ND | 299 |
| ribitol | ND | 147 | ND | ND | 360 | 1075 | 614 |
| ribonic acid | ND | 137 | ND | ND | 358 | 1076 | 613 |
| ribose | ND | 70 | ND | ND | 208 | 316 | 946 |
| ribose-5-phosphate | ND | ND | ND | ND | ND | ND | 306 |
| ribulose | ND | ND | ND | ND | ND | 260 | ND |
| ribulose-5-phosphate | ND | ND | ND | ND | 312 | ND | ND |
| salicylaldehyde | ND | 424 | ND | ND | ND | ND | 230 |

| | | | | | | | |
|---|---|---|---|---|---|---|---|
| salicylic acid | ND | ND | ND | ND | ND | 1079 | ND |
| sarcosine | ND | ND | ND | ND | 268 | ND | ND |
| serine | ND | 149 | ND | ND | ND | 1080 | ND |
| shikimic acid | ND | 248 | ND | ND | ND | 1081 | ND |
| sophorose | ND | ND | ND | ND | 55 | 198 | 930 |
| sorbitol | ND | 358 | ND | ND | 29 | 1083 | 931 |
| spermidine | ND | 255 | ND | ND | ND | ND | ND |
| sphingomyelin d33:1 | ND | ND | 104 | ND | ND | ND | ND |
| sphingomyelin d34:0 | ND | ND | 270 | ND | ND | ND | ND |
| sphingomyelin d34:1 | ND | ND | 111 | ND | ND | ND | ND |
| sphingomyelin d34:2 | ND | ND | 327 | ND | ND | ND | ND |
| sphingomyelin d36:1 | ND | ND | 213 | ND | ND | ND | ND |
| sphingomyelin d36:2 | ND | ND | 352 | ND | ND | ND | ND |
| sphingomyelin d37:1 | ND | ND | 112 | ND | ND | ND | ND |
| sphingomyelin d38:2 | ND | ND | 171 | ND | ND | ND | ND |
| sphingomyelin d40:2 | ND | ND | 49 | ND | ND | ND | ND |
| sphingomyelin d41:2 | ND | ND | 27 | ND | ND | ND | ND |
| sphingomyelin d42:3 | ND | ND | 356 | ND | ND | ND | ND |
| stearic acid | ND | ND | ND | ND | ND | 1084 | ND |
| suberic acid | ND | ND | ND | ND | ND | 1085 | ND |
| succinate semialdehyde | ND | 185 | ND | ND | 326 | 1086 | 886 |
| succinic acid | ND | 405 | ND | ND | 265 | 1072 | 658 |
| sucrose | ND | 303 | ND | ND | 156 | ND | ND |
| sulfuric acid | ND | ND | ND | ND | ND | 581 | ND |
| taurine | ND | ND | ND | ND | ND | ND | 929 |
| terephthalic acid | ND | ND | ND | ND | ND | 364 | ND |
| tetracosane | ND | ND | ND | ND | ND | ND | 147 |
| tetracosanol | ND | ND | ND | ND | ND | ND | 309 |
| threitol | ND | ND | ND | ND | 158 | ND | ND |
| threonic acid | ND | 24 | ND | ND | 333 | 58 | 927 |
| threonine | ND | 517 | ND | ND | ND | 551 | ND |
| thymidine | ND | ND | ND | ND | ND | 553 | ND |
| thymine | ND | ND | ND | ND | ND | ND | 264 |
| trans-4-hydroxyproline | ND | 233 | ND | ND | 304 | 555 | 925 |
| triacylglycerol 34:0 | ND | ND | 10 | ND | ND | ND | ND |
| triacylglycerol 38:0 | ND | ND | 147 | ND | ND | ND | ND |
| triacylglycerol 40:0 | ND | ND | 315 | ND | ND | ND | ND |
| triacylglycerol 40:1 | ND | ND | 305 | ND | ND | ND | ND |
| triacylglycerol 42:1 | ND | ND | 19 | ND | ND | ND | ND |
| triacylglycerol 42:2 | ND | ND | 214 | ND | ND | ND | ND |
| triacylglycerol 42:3 | ND | ND | 313 | ND | ND | ND | ND |
| triacylglycerol 43:3 | ND | ND | 275 | ND | ND | ND | ND |
| triacylglycerol 44:1 | ND | ND | 254 | ND | ND | ND | ND |

| | | | | | | | |
|---|---|---|---|---|---|---|---|
| triacylglycerol 44:2 | ND | ND | 247 | ND | ND | ND | ND |
| triacylglycerol 44:3 | ND | ND | 349 | ND | ND | ND | ND |
| triacylglycerol 46:1 | ND | ND | 155 | ND | ND | ND | ND |
| triacylglycerol 46:2 | ND | ND | 9 | ND | ND | ND | ND |
| triacylglycerol 46:3 | ND | ND | 56 | ND | ND | ND | ND |
| triacylglycerol 46:4 | ND | ND | 318 | ND | ND | ND | ND |
| triacylglycerol 46:5 | ND | ND | 328 | ND | ND | ND | ND |
| triacylglycerol 48:0 | ND | ND | 98 | ND | ND | ND | ND |
| triacylglycerol 48:1 | ND | ND | 174 | ND | ND | ND | ND |
| triacylglycerol 48:3 | ND | ND | 18 | ND | ND | ND | ND |
| triacylglycerol 48:4 | ND | ND | 157 | ND | ND | ND | ND |
| triacylglycerol 49:0 | ND | ND | 216 | ND | ND | ND | ND |
| triacylglycerol 49:2 | ND | ND | 245 | ND | ND | ND | ND |
| triacylglycerol 49:4 | ND | ND | 294 | ND | ND | ND | ND |
| triacylglycerol 50:2 | ND | ND | 239 | ND | ND | ND | ND |
| triacylglycerol 50:3 | ND | ND | 24 | ND | ND | ND | ND |
| triacylglycerol 50:4 | ND | ND | 31 | ND | ND | ND | ND |
| triacylglycerol 50:5 | ND | ND | 296 | ND | ND | ND | ND |
| triacylglycerol 50:6 | ND | ND | 118 | ND | ND | ND | ND |
| triacylglycerol 51:1 | ND | ND | 209 | ND | ND | ND | ND |
| triacylglycerol 51:2 | ND | ND | 57 | ND | ND | ND | ND |
| triacylglycerol 51:3 | ND | ND | 54 | ND | ND | ND | ND |
| triacylglycerol 51:4 | ND | ND | 96 | ND | ND | ND | ND |
| triacylglycerol 51:5 | ND | ND | 193 | ND | ND | ND | ND |
| triacylglycerol 52:0 | ND | ND | 145 | ND | ND | ND | ND |
| triacylglycerol 52:1 | ND | ND | 220 | ND | ND | ND | ND |
| triacylglycerol 52:2 | ND | ND | 8 | ND | ND | ND | ND |
| triacylglycerol 52:3 | ND | ND | 116 | ND | ND | ND | ND |
| triacylglycerol 52:4 | ND | ND | 180 | ND | ND | ND | ND |
| triacylglycerol 52:5 | ND | ND | 293 | ND | ND | ND | ND |
| triacylglycerol 52:7 | ND | ND | 332 | ND | ND | ND | ND |
| triacylglycerol 53:1 | ND | ND | 312 | ND | ND | ND | ND |
| triacylglycerol 53:2 | ND | ND | 185 | ND | ND | ND | ND |
| triacylglycerol 53:3 | ND | ND | 97 | ND | ND | ND | ND |
| triacylglycerol 53:5 | ND | ND | 60 | ND | ND | ND | ND |
| triacylglycerol 54:1 | ND | ND | 235 | ND | ND | ND | ND |
| triacylglycerol 54:2 | ND | ND | 250 | ND | ND | ND | ND |
| triacylglycerol 54:3 | ND | ND | 69 | ND | ND | ND | ND |
| triacylglycerol 54:6 | ND | ND | 178 | ND | ND | ND | ND |
| triacylglycerol 54:7 | ND | ND | 287 | ND | ND | ND | ND |
| triacylglycerol 54:9 | ND | ND | 134 | ND | ND | ND | ND |
| triacylglycerol 55:1 | ND | ND | 342 | ND | ND | ND | ND |
| triacylglycerol 55:2 | ND | ND | 143 | ND | ND | ND | ND |

| | | | | | | | |
|---|---|---|---|---|---|---|---|
| triacylglycerol 55:3 | ND | ND | 195 | ND | ND | ND | ND |
| triacylglycerol 56:1 | ND | ND | 264 | ND | ND | ND | ND |
| triacylglycerol 56:10 | ND | ND | 236 | ND | ND | ND | ND |
| triacylglycerol 56:2 | ND | ND | 346 | ND | ND | ND | ND |
| triacylglycerol 56:3 | ND | ND | 330 | ND | ND | ND | ND |
| triacylglycerol 56:5 | ND | ND | 26 | ND | ND | ND | ND |
| triacylglycerol 56:6 | ND | ND | 194 | ND | ND | ND | ND |
| triacylglycerol 56:7 | ND | ND | 51 | ND | ND | ND | ND |
| triacylglycerol 56:8 | ND | ND | 58 | ND | ND | ND | ND |
| triacylglycerol 56:9 | ND | ND | 347 | ND | ND | ND | ND |
| triacylglycerol 57:2 | ND | ND | 240 | ND | ND | ND | ND |
| triacylglycerol 57:4 | ND | ND | 282 | ND | ND | ND | ND |
| triacylglycerol 58:10 | ND | ND | 78 | ND | ND | ND | ND |
| triacylglycerol 58:3 | ND | ND | 234 | ND | ND | ND | ND |
| triacylglycerol 58:4 | ND | ND | 204 | ND | ND | ND | ND |
| triacylglycerol 58:5 | ND | ND | 298 | ND | ND | ND | ND |
| triacylglycerol 58:6 | ND | ND | 184 | ND | ND | ND | ND |
| triacylglycerol 58:9 | ND | ND | 177 | ND | ND | ND | ND |
| triacylglycerol 59:3 | ND | ND | 258 | ND | ND | ND | ND |
| triacylglycerol 60:11 | ND | ND | 191 | ND | ND | ND | ND |
| triacylglycerol 60:4 | ND | ND | 154 | ND | ND | ND | ND |
| triacylglycerol 60:5 | ND | ND | 55 | ND | ND | ND | ND |
| triacylglycerol 60:6 | ND | ND | 12 | ND | ND | ND | ND |
| triacylglycerol 68:4;O2 | ND | ND | 126 | ND | ND | ND | ND |
| triacylglycerol 70:3;O2 | ND | ND | 351 | ND | ND | ND | ND |
| triacylglycerol 70:4;O2 | ND | ND | 357 | ND | ND | ND | ND |
| triacylglycerol 70:5;O2 | ND | ND | 226 | ND | ND | ND | ND |
| triacylglycerol 72:5;O2 | ND | ND | 362 | ND | ND | ND | ND |
| triacylglycerol 72:6;O2 | ND | ND | 326 | ND | ND | ND | ND |
| triacylglycerol 72:7;O2 | ND | ND | 335 | ND | ND | ND | ND |
| triacylglycerol O-38:2 | ND | ND | 146 | ND | ND | ND | ND |
| triacylglycerol O-50:1 | ND | ND | 271 | ND | ND | ND | ND |
| triacylglycerol O-50:2 | ND | ND | 135 | ND | ND | ND | ND |
| triacylglycerol O-52:0 | ND | ND | 173 | ND | ND | ND | ND |
| triacylglycerol O-52:1 | ND | ND | 337 | ND | ND | ND | ND |
| triacylglycerol O-52:2 | ND | ND | 117 | ND | ND | ND | ND |
| triacylglycerol O-52:3 | ND | ND | 16 | ND | ND | ND | ND |
| triacylglycerol O-54:1 | ND | ND | 127 | ND | ND | ND | ND |
| triacylglycerol O-54:2 | ND | ND | 289 | ND | ND | ND | ND |
| triacylglycerol O-54:3 | ND | ND | 269 | ND | ND | ND | ND |
| triacylglycerol O-58:2 | ND | ND | 304 | ND | ND | ND | ND |
| tryptophan | ND | ND | ND | ND | 183 | 556 | 924 |
| tyrosine | ND | 69 | ND | ND | 404 | 561 | ND |

| | | | | | | | |
|---|---|---|---|---|---|---|---|
| uracil | ND | 435 | ND | ND | ND | 259 | ND |
| urea | ND | 212 | ND | ND | 420 | ND | 298 |
| uric acid | ND | ND | ND | ND | 149 | 562 | 181 |
| uridine | ND | ND | ND | ND | ND | 565 | 922 |
| valine | ND | ND | ND | ND | 379 | 566 | ND |
| xanthine | ND | 198 | ND | ND | ND | 234 | ND |
| xanthosine | ND | ND | ND | ND | ND | ND | 921 |
| xylitol | ND | 417 | ND | ND | 401 | 188 | 920 |
| xylonic acid | ND | ND | ND | ND | 291 | ND | ND |
| xylonic acid isomer | ND | 336 | ND | ND | ND | ND | ND |
| xylonolactone | ND | ND | ND | ND | ND | 572 | 919 |
| xylose | ND | 304 | ND | ND | 203 | ND | 252 |
| xylulose | ND | 418 | ND | ND | 580 | 550 | 915 |
