## Supplemental Figures for "Multi-Tissue Metabolomic Signatures of Five Longevity Interventions Converge on Ergothioneine and Lipid Remodeling in Male UM-HET3 Mice"

**Supplementary table 1.** Number of metabolites (features) in each tissue dataset used in this study. Tissues: brain, perigonadal fat, inguinal fat, kidney, liver, muscle, plasma.

| Tissue | Number of Features |
|---|---|
| Brain | 572 |
| Perigonadal Fat | 395 |
| Inguinal Fat | 755 |
| Kidney | 946 |
| Liver | 580 |
| Muscle | 1,087 |
| Plasma | 1,051 |

**Supplemental Table 2**. Category over-representation by tissue at 90% feature explainability power for each individual tissue. The "Dataset Count" is the number of times the metabolite category is found in the whole dataset for a particular tissue. The "Observed Count" is the number of times that the metabolite category is observed in the top 90% of predictive features, as given in the "Metabolites needed to explain 90% Prediction column". The statistical test performed on the data was a binomial test for skew; raw and Benjamini-Hochberg adjusted ("FDR-BH") p-values are reported.

| Tissue | Metabolites needed to explain 90% Prediction | Metabolite Category | Observed Count | Dataset Count | Binomial Skew P Value | FDR BH |
|---|---|---|---|---|---|---|
| Brain | 64 | Biogenic amine | 5 | 12 | 1.1E-02 | 1.2E-01 |
| Perigonadal Fat | 58 | FA (fatty acid) | 17 | 55 | 2.0E-03 | 9.9E-03* |
| Perigonadal Fat | 58 | DG (diacylglycerol) | 7 | 22 | 4.2E-02 | 1.0E-01 |
| Kidney | 11 | PC (phosphatidylcholine) | 5 | 105 | 4.4E-03 | 8.7E-03* |
| Liver | 9 | SM (sphingomyelin) | 2 | 23 | 4.7E-02 | 1.2E-01 |
| Muscle | 96 | DG (diacylglycerol) | 9 | 42 | 1.2E-02 | 1.3E-01 |
| Muscle | 96 | Ceramide / Glucosylceramide | 5 | 21 | 3.9E-02 | 2.1E-01 |
| Plasma | 52 | TAG (triacylglycerol) | 19 | 170 | 4.1E-04 | 2.4E-03* |
| Plasma | 52 | DG (diacylglycerol) | 6 | 23 | 9.4E-04 | 2.8E-03* |

**Supplementary table 3.** Number of iterations until convergence for the XGBoost algorithm applied to each tissue type. Tissues: brain, perigonadal fat, inguinal fat, kidney, liver, muscle, plasma.

| Tissue | Convergence Iterations XGBoost Gain |
|---|---|
| Brain | 300 |
| Perigonadal Fat | 400 |
| Inguinal Fat | 400 |
| Kidney | 500 |
| Liver | 300 |
| Muscle | 500 |
| Plasma | 400 |

**Supplementary table 4. Ratio of total phosphatidylcholine (PC) in pooled male longevity interventions versus control, by tissue.** For each tissue, metabolite columns classified as phosphatidylcholine (PC) were identified (LysoPC and sphingomyelin excluded). Linear-scale intensities (log2_intensity) were summed over all PC species and over samples in control or in pooled male treatment groups (rapamycin, acarbose, canagliflozin, 17α-estradiol, calorie restriction). The table reports the ratio of total treatment PC to total control PC and its $log_2$ value (log2_PC_ratio). “n PC species” is the number of PC metabolites detected per tissue.

| Tissue | n PC species | PC ratio | log2 PC ratio |
|---|---|---|---|
| Brain | 49 | 2.90 | 1.54 |
| Perigonadal Fat | 23 | 3.72 | 1.90 |
| Inguinal Fat | 40 | 4.07 | 2.02 |
| Kidney | 105 | 2.80 | 1.49 |
| Liver | 72 | 2.87 | 1.52 |
| Muscle | 138 | 3.39 | 1.76 |
| Plasma | 117 | 2.46 | 1.30 |

**Supplementary Table 5. Cross-tissue metabolite feature importance rankings from 1,000 XGBoost gain iterations.** Attached as a separate file.

**Supplementary Table 6 Convergence of XGBoost Gain feature importance rankings across tissues.** Convergence iteration indicates the first step at which Spearman rank correlation ($\rho_T$) >= 0.98 and total variation distance ($D_T$) <= 0.02 were met for two successive iteration steps, evaluated on a fixed top-feature set defined at 1,000 iterations. All tissues achieved near-perfect rank stability ($\rho_T > 0.99$, $D_T < 0.01$) by 1,000 iterations.

| Tissue | N Features | Convergence Iteration | Final ρT (1,000 iter.) | Final DT (1,000 iter.) |
|---|---|---|---|---|
| Brain | 572 | 300 | 0.9994 | 0.0037 |
| Liver | 580 | 300 | 1.0000 | 0.0038 |
| Plasma | 1,051 | 400 | 0.9992 | 0.0057 |
| Gonadal Fat | 395 | 400 | 0.9981 | 0.0054 |
| Inguinal Fat | 755 | 400 | 0.9964 | 0.0049 |
| Kidney | 946 | 500 | 0.9964 | 0.0078 |
| Muscle | 1,087 | 500 | 0.9979 | 0.0076 |
